# Quasi-continuous cotranslational compaction and folding of a multidomain protein

**DOI:** 10.64898/2026.02.04.703729

**Authors:** Spyridoula Mitsikosta, Justin M. Westerfield, Fátima Pardo-Avila, Michael Levitt, Gunnar von Heijne, Ane Metola

## Abstract

Most proteins start to fold cotranslationally as they come off the ribosome. So far, studies of cotranslational folding have focused mainly on small, single-domain proteins. Here, we have used Force Profile Analysis to study the cotranslational folding of Firefly luciferase, a complex 550-residue protein composed of an N-terminal domain (NTD) encompassing two split Rossmann folds (RF-1, RF-2) and a β-roll, and a flexibly attached C-terminal domain (CTD). The folding process is characterized by a quasi-continuous series of compaction/folding steps that generate intermediate-size pulling forces on the nascent chain, punctuated by a prominent high-force event that represents the folding of the RF-2 domain, and a few low-force instances that likely indicate the formation of distinct folding intermediates. Trigger Factor interacts extensively with the nascent chain when the central part of RF-2 and the early parts of the CTD are synthesized. Our analysis uncovers a cotranslational compaction/folding process that is rich in detail and not just a simple succession of a few distinct, cooperative folding transitions.

## Introduction

While it has long been acknowledged that many proteins may begin to fold cotranslationally *in vivo*, it is only relatively recently that a range of methodological advances have opened up this area to more detailed studies. Early work mainly used assays based on protease resistance^1,2^, while today powerful new biochemical and biophysical methods such as force-profile analysis (FPA)^3^, pulse-proteolysis^4^, NMR^5^, hydrogen-deuterium exchange mass-spectrometry (HDX-MS)^6^, cryo-electron microscopy (cryo-EM)^3^, optical tweezers^7^, fluorescence-resonance energy transfer (FRET)^8^, and computational approaches^9^ have advanced the field to a new level.

Thus, over the past ∼10 years, it has been shown that small protein domains (up to ∼100 residues in size) can fold wholly or in part inside the ribosome exit tunnel (ET)^10,11^, that unfolded protein domains held in proximity to the ribosome can interact non-specifically with the surface of the ribosome and that on-ribosome folding reduces the entropic penalty of folding and promotes formation of partially folded intermediates^12^, that folding pathways can differ dramatically when folding happens on-ribosome as compared to off-ribosome^13^, and that a large proportion of proteins likely start to fold cotranslationally^14^. Moreover, cotranslational nascent chain—chaperone interactions have been shown to play important roles during protein folding *in vivo*^15^.

Many of these studies have focused on single-domain proteins that are small enough to be able to fold inside the ET, or in the vestibule that defines the exit port at the distal end of the ET. Larger proteins, however, especially those that are composed of identifiable but tightly interacting or interdigitating subdomains, are more challenging and only a handful such studies have been published to date^13^. We have now applied FPA to one such protein, Firefly Luciferase (FLuc), a 550-residue protein that has been much studied in the context of chaperone-mediated folding^16–19^. FLuc is composed of a large N-terminal domain (NTD) encompassing two split Rossmann folds (RF-1, RF-2) and a β-roll, and a flexibly attached C-terminal α/β domain (CTD), Fig. 1. FLuc folds efficiently during *in vitro* translation even in the absence of chaperones^20^, but requires chaperones to refold from a denatured state^21^. Cotranslational folding of FLuc was previously proposed to proceed via the formation of an N-terminal protease-resistant 21 kDa intermediate^2^, and a recent HDX-MS study has shown that, when synthesized on *E. coli* ribosomes, FLuc adopts a close-to-native structure before, or at the latest at, the time when the NTD has emerged from the ET^22^.

**Figure 1.**
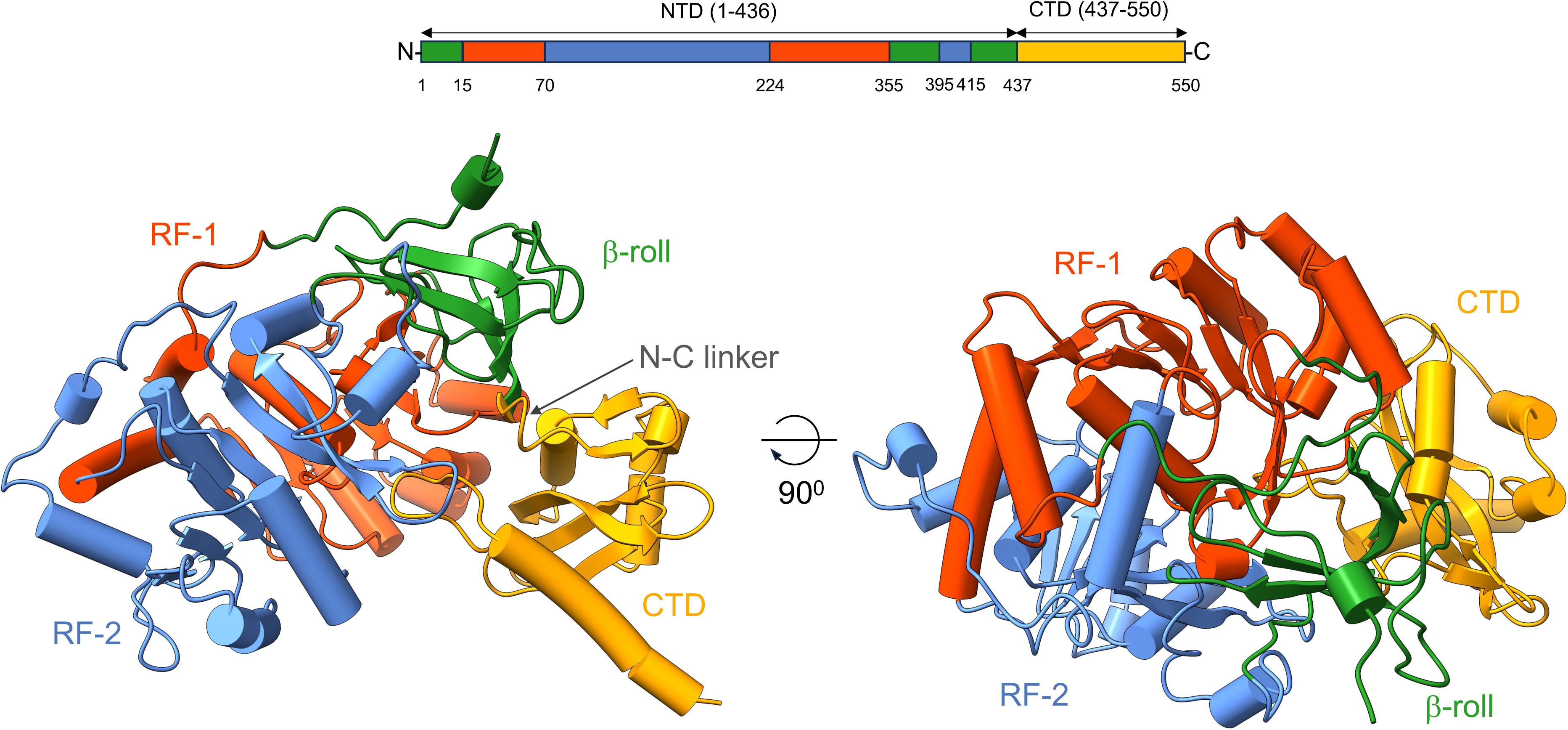
FLuc domain structure. FLuc 3D structure (AlphaFold DB^61^ AF-P08659-F1-v4). The two Rossmann folds are colored in orange (RF-1) and blue (RF-2), the β-roll domain in green, and the C-terminal domain (CTD) in yellow. Approximate domain boundaries are indicated at the top.

Here, we report the first high-resolution FPA study of the cotranslational folding of FLuc. We find that FLuc undergoes a quasi-continuous series of cotranslational compaction/folding steps, punctuated by a prominent folding event and a few quiescent periods that likely correspond to the formation of distinct folding intermediates. The chaperone Trigger Factor (TF) strongly affects the folding of a portion of the first half of the NTD and of an early portion of the CTD, but otherwise has no or only highly localized effects. The quasi-continuous character of the folding process raises questions about what should be considered a cotranslational folding intermediate and how such intermediates may relate to cooperative folding events during translation.

## Results

### Force Profile Analysis

FPA is based on the original discovery^23^ of translational arrest peptides (APs) – short polypeptide sequences in nascent chains (NCs) that make specific interactions with ribosomal rRNA in the vicinity of the peptidyl transferase center (PTC) and thereby induce slight changes in the relative orientation of the A- and P-site tRNAs that lead to strong reductions in the rate of transpeptidation^24^. An external pulling force acting on the AP can dislodge it from its binding site and thereby restore the normal translation rate^25–27^; APs can thus be used as force sensors to report on various cotranslational processes that generate force on the AP such as protein folding^3^, membrane protein biogenesis^28^, or protein translocation^29^. Exceptionally strong APs can even be used to irreversibly stall a NC on the ribosome^30^, making it possible to purify stable ribosome-nascent chain complexes (RNCs) for analysis by techniques such as cryo-EM^3^, NMR^12^, and HDX-MS^22^.

In a typical FPA experiment^3^, an intermediate-strength AP (e.g., the *E. coli* SecM(*Ec*) AP) is fused via a linker to the C-terminal end of the target protein, and a C-terminal tail is added behind the AP, Fig. 2a. A series of constructs is made by truncating one or a few residues at a time from the C-terminal end of the linker sequence and continuing into the target protein. Each construct is then translated and pulse-labelled with [^35^S]-Met for a set time, either *in vivo* or in an *in vitro* translation system such as PURExpress™, and analyzed by SDS-PAGE, Fig. 2b. The bands corresponding to arrested (*A*) and full-length (*FL*) protein are quantitated on a Phosphorimager, and the fraction full-length protein (*f_FL_* = *I_FL_*/(*I_FL_*+*I_A_*), where *I_A_* and *I_FL_* are the measured intensities of the respective bands) is calculated for each construct. *f_FL_* serves as a convenient proxy for the pulling force acting on the AP, and can in principle be converted into force measured in pN^31^; the higher the pulling force, the higher is *f_FL_*.

**Figure 2.**
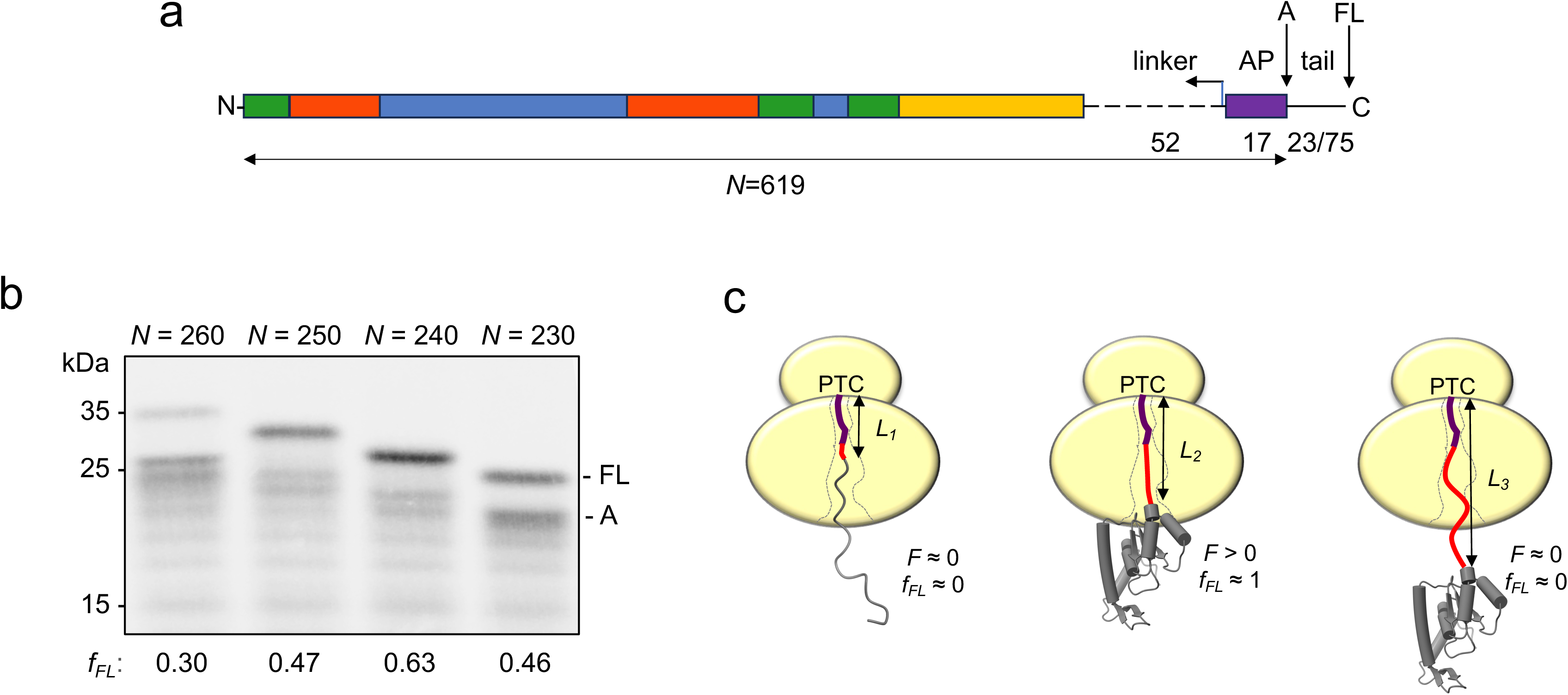
Force Profile Analysis of FLuc. (a) Construct design. In the construct shown (*N* = 619 residues), full-length FLuc (coloring as in Fig. 1) is fused to a 52-residue GS-linker, the 17-residue SecM(*Ec*) AP (FSTPVWISQAQGIRAGP), and a 75-residue C-terminal tail; in constructs with *N* ≤ 250 residues, the C-terminal tail is 23 residues long (see Supplemental Table S1 for sequences). The full set of constructs was made by deleting 5 residues at a time starting from the C-terminal end of the linker (left-pointing arrow). The shortest construct (*N* = 70 residues) thus contains FLuc residues 1-50, a 3-residue GS-linker, the 17-residue SecM(*Ec*) AP, and a 23-residue tail. The 17-residue AP is constant in all constructs. Arrested (*A*) and full-length (*FL*) products are indicated. (b) Example SDS-PAGE gel showing constructs *N* = 230-260 residues. The fraction full-length protein is calculated as *f_FL_* = *I_FL_*/(*I_FL_*+*I_A_*), where *I_FL_* and *I_A_* are the intensities of the *FL* and *A* bands, respectively. The band immediately below the *A* product indicates a ribosome stacked behind the arrested ribosome. (c) Cartoon model illustrating how cotranslational folding of a protein domain generates force on the AP. In constructs with a short linker (red), the C-terminal end of the domain is located deep inside the ET at the point when the ribosome reaches the C-terminal end of the AP (purple), there is not enough room for the domain to fold, and hence little force is generated (*F* ≈ 0, *f_FL_* ≈ 0; left). At intermediate linker lengths, the domain starts to fold at the point when the ribosome reaches the C-terminal end of the AP, generating force on the AP by stretching the NC (*F* > 0, *f_FL_* ≈ 1; middle). Finally, in constructs with long linkers, the domain has already folded at the point when the ribosome reaches the C-terminal end of the AP, and again no force is generated (*F* ≈ 0, *f_FL_* ≈ 0; right).

Finally, a force profile (FP) is obtained by plotting *f_FL_* against *N*, the number of residues between the N-terminal end of the construct and the C-terminal end of the AP. In the case of cotranslational protein folding, peaks in the FP correspond to constructs in which a (sub)domain can just reach a region in or immediately outside the ET where there is enough space for it to fold^3,10^, Fig. 2c. Some of the free energy of folding will thus be stored as elastic energy in the linker (red in Fig. 2c) that attaches the subdomain to the AP, generating a pulling force on the AP and a high *f_FL_* value^32^.

### A FLuc FP at five-residue resolution

A set of FLuc constructs was prepared starting from a design with full-length FLuc (residues 1-550) fused to a 52-residue Gly-Ser (GS) linker, a 17-residue SecM(*Ec*) AP (FSTPVWISQAQGIRAGP), and a 75-residue C-terminal tail, Fig. 2a. Shorter constructs were designed by deleting five residues at a time starting from the C-terminal end of the GS-linker and continuing until only FLuc residues 1-50 remained; constructs with *N* ≤ 250 residues were designed with a shorter, 23-residue C-terminal tail. All in all, the full set consists of 128 constructs, with each construct referred to by its *N*-value, *i.e*., the number of residues from the N-terminal end of FLuc to the C-terminal Pro residue in the AP. All constructs were transcribed and translated *in vitro* in the *E. coli*-derived PURExpress™ system, and analyzed by SDS-PAGE. The resulting FLuc FP, obtained at five-residue resolution (and two-residue resolution around a few peaks and dips), is shown in Fig. 3 (blue curve). The FP extends from *N* = 70 residues (FLuc residues 1-50 fused via a 3-residue GS-linker to the 17-residue AP and a 23-residue C-terminal tail) to *N* = 619 (FLuc residues 1-550 fused via a 52-residue GS-linker to the 17-residue AP and a 75-residue C-terminal tail).

**Figure 3.**
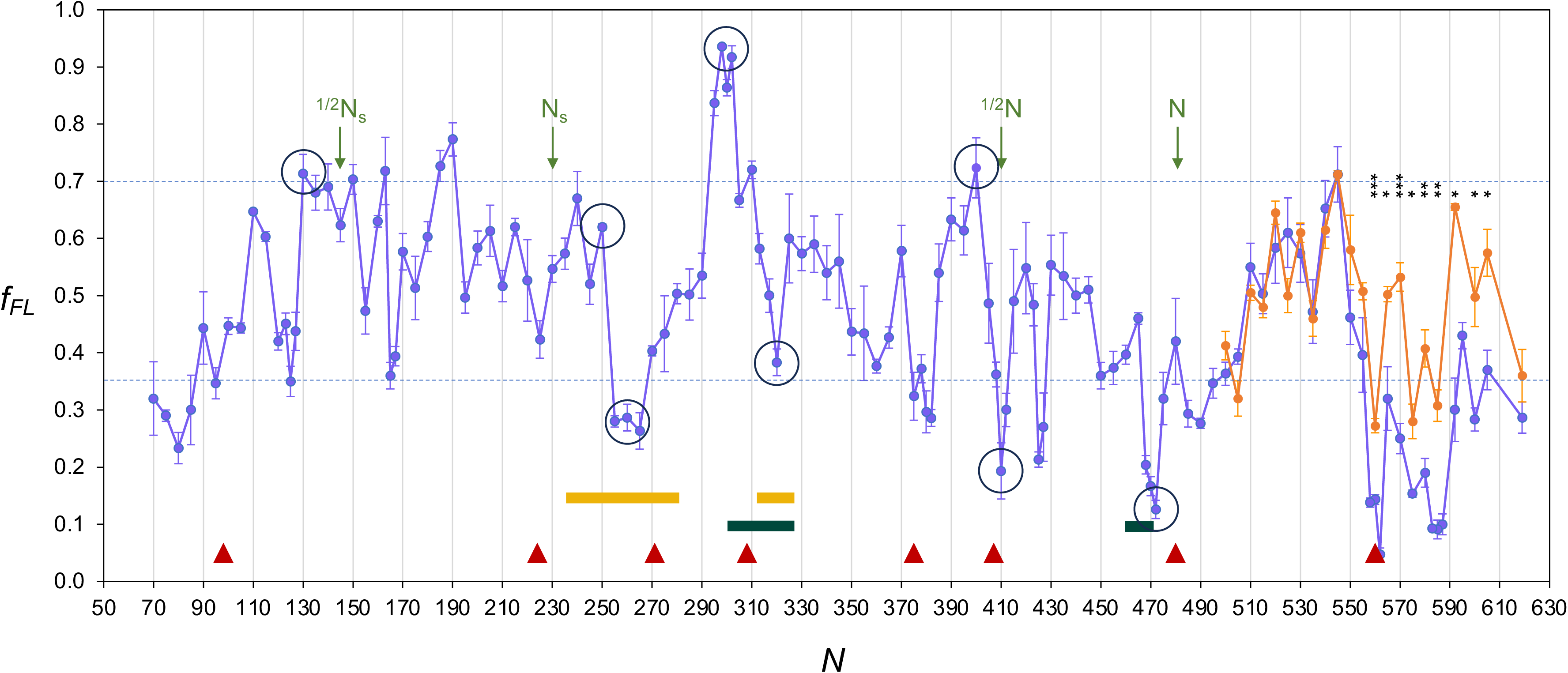
FLuc force profile. Average *f_FL_* values plotted against *N* for FLuc constructs (in blue), and for the CTD expressed by itself (orange). Peaks and dips discussed in the main text are encircled. The full set of subdomain boundaries predicted by the SWORD2 algorithm^35^ are indicated by red triangles, peptides showing increased deuterium uptake compared to folded full-length FLuc in the stalled ^1/2^N construct^22^ are indicated by yellow bars, cotranslational folding intermediates predicted by the Jacobs and Shakhnovitch method^36^ are indicated by dark green bars, and four constructs previously analyzed by HDX-MS^22^ are indicated by their names (^1/2^N_s_, N_s_, ^1/2^N, N; in green). Significant differences between the FLuc and CTD data are indicated by *p*-values calculated by Student’s t-test (*p* < 0.05: *; *p* < .01: **, *p* < .001: ***). Error bars indicate the standard error of the mean (SEM).

The FP is characterized by stretches with many small peaks and dips of intermediate amplitude (0.35 ≲*f_FL_* ≲ 0.7) punctuated by a few high peaks and deep dips. Previous work on a range of single-domain proteins have shown that cooperative folding events happening in or just outside the ribosome exit tunnel (ET) give rise to distinct peaks in the FP^3,10,31–34^. Conversely, we hypothesize that deep dips in the FP correspond to points during translation where a (semi)stable folding intermediate has just been formed while the following part of the nascent chain (NC) has not yet grown sufficiently in size to re-initiate folding.

To look for correlations between the FP and possible folding intermediates, we used the SWORD2 server^35^ to parse the FLuc 3D structure into subdomains (red triangles indicate predicted domain boundaries) and the Jacobs & Shakhnovic method^36^ to predict the formation of potential cotranslational folding intermediates (dark green bars), Fig. 3 and Supplemental Fig. S1. Because the ET sequesters ∼40 residues of the NC^37^ and protein domains composed of more than ∼120 amino acids typically start to fold when their C-terminus is ∼40 residues away from the peptidyl-transferase center (PTC) in the ribosome^1,10^, we estimated *N*-values for the predicted subdomain boundaries and folding intermediates by adding 40 residues to the predicted boundaries (*e.g*., a predicted boundary at FLuc residue 265 is plotted at *N* = 305 residues in Fig. 3). The SWORD2 boundaries correlate quite well with dips in the FP, and the two predicted folding intermediates correspond to the main peak-dip features at *N* ≈ 300-325 residues and *N* ≈ 460-470 residues.

The first main peak at *N* = 130 residues, and the first major dip at *N* = 255 residues, suggest that some degree of compaction/folding commences approximately when FLuc residues 1-95 have emerged outside the ET and continues until what appears to be a compact subdomain comprising residues 1-215 (±5 residues) has formed outside the ET, Fig. 3. This subdomain corresponds well to the domain boundary predicted at *N* = 271 residues. Available HDX-MS data for stalled *E. coli* FLuc RNCs corresponding to constructs *N* = 145 (^1/2^N_S_, Fig. 3) and *N* = 230 (N_S,_ Fig. 3) indicate that the parts exposed outside the ribosome have not adopted the native structure in the former, and that native-like structure is seen only for FLuc residues ∼160-190 in the latter^22^. Early work demonstrated the cotranslational formation of a protease-resistant 21 kDa FLuc intermediate comprising residues 1-190^2^; a recent study has shown that this may represent a non-native homodimer produced during the proteolysis experiment^22^. We thus propose that FLuc residues 1-215 (±5) define an early compacted folding intermediate with some native-like structure.

The prominent dip in the FP at *N* = 255-265 residues indicates that the segment containing FLuc residues ∼215-225 does not fold onto the 1-215 folding intermediate, but rather remains in a flexible conformation as the NC elongates. Interestingly, HDX-MS data^22^ for the stalled ^1/2^N construct (corresponding to our *N* = 410 construct) shows increased deuterium uptake compared to folded full-length FLuc for peptides covering FLuc residues 195-238 and 272-287, matching the dips in the FP at *N* = 255-265 and 315-325 residues (yellow bars, Fig. 3). Thus, FLuc residues ∼215-225 and ∼275-285 remain in a flexible conformation at least until translation has proceeded to a NC-length of 410 residues.

A very strong pulling force is apparent at *N* = 295-310 residues, signaling a new folding event. This peak has the highest amplitude of all. To precisely map its origin, we made a “GS-scan”^33^ where we replaced an increasing number of residues at the C-terminal end of the FLuc part in the *N* = 300 construct by a repeating Gly-Ser sequence. As seen in Fig. 4a, the amplitude of the peak is reduced when ∼20 or more C-terminal FLuc residues are replaced by an equally long GS-stretch. Because the SecM(*Ec*) AP part is 20 residues long (Fig. 2a), this means that the C-terminal end of the part of FLuc that folds in this construct is ∼40 residues distant from the PTC, as anticipated. In addition, a series of N-terminal deletions in the *N* = 300 residues construct shows that residues 4-80 can be deleted with no effect on the amplitude of the *N* = 300 peak, while longer deletions ending at residues 90 to 170 lead to a gradual reduction in peak amplitude, Fig. 4b. The peak at *N* = 295-310 residues is thus generated by the folding of residues ∼80 to ∼260, and the potential folding intermediate corresponding to the dip at *N* = 320 residues neatly corresponds to the FLuc RF-2 domain (except for the last, non-contiguous RF-2 segment M^396^–G^416^) and a small part of RF-1 (most notably the hydrophobic helix G^246^–C^258^), Fig. 5. We cannot tell from the FP-data whether the N-terminal part of RF-1 (residues 16-70) is also folded in the *N* = 320 construct, only that it does not contribute to the generation of the *N* = 300 peak. Notably, however, proteinase K treatment of stalled ^1/2^N ribosomes (corresponding to our *N* = 410 construct, Fig. 3) yields a ∼35 kDa PK-resistant N-terminal FLuc fragment and HDX-MS analysis of ^1/2^N RNCs shows that FLuc residues 1 to ∼270 mostly adopt native-like structure^22^, arguing that the first half of RF-1 and most of RF-2 are folded (except for residues 215-225 and 275-285, as noted above). Consistent with this, the coarse-grained structure-based model of cotranslational folding developed by Jacobs & Shaknovich^36,38^ predicts that a first, stable folding intermediate should form when FLuc residues ∼260-280 emerge from the ET at *N* ≈ 300 residues, Supplemental Fig. S1. We conclude that the RF-2 domain folds into a close-to-native structure at *N* ≈ 300 residues and that the first half of the RF-1 domain also is folded at this stage, although the RF-2 domain can fold in its absence.

**Figure 4.**
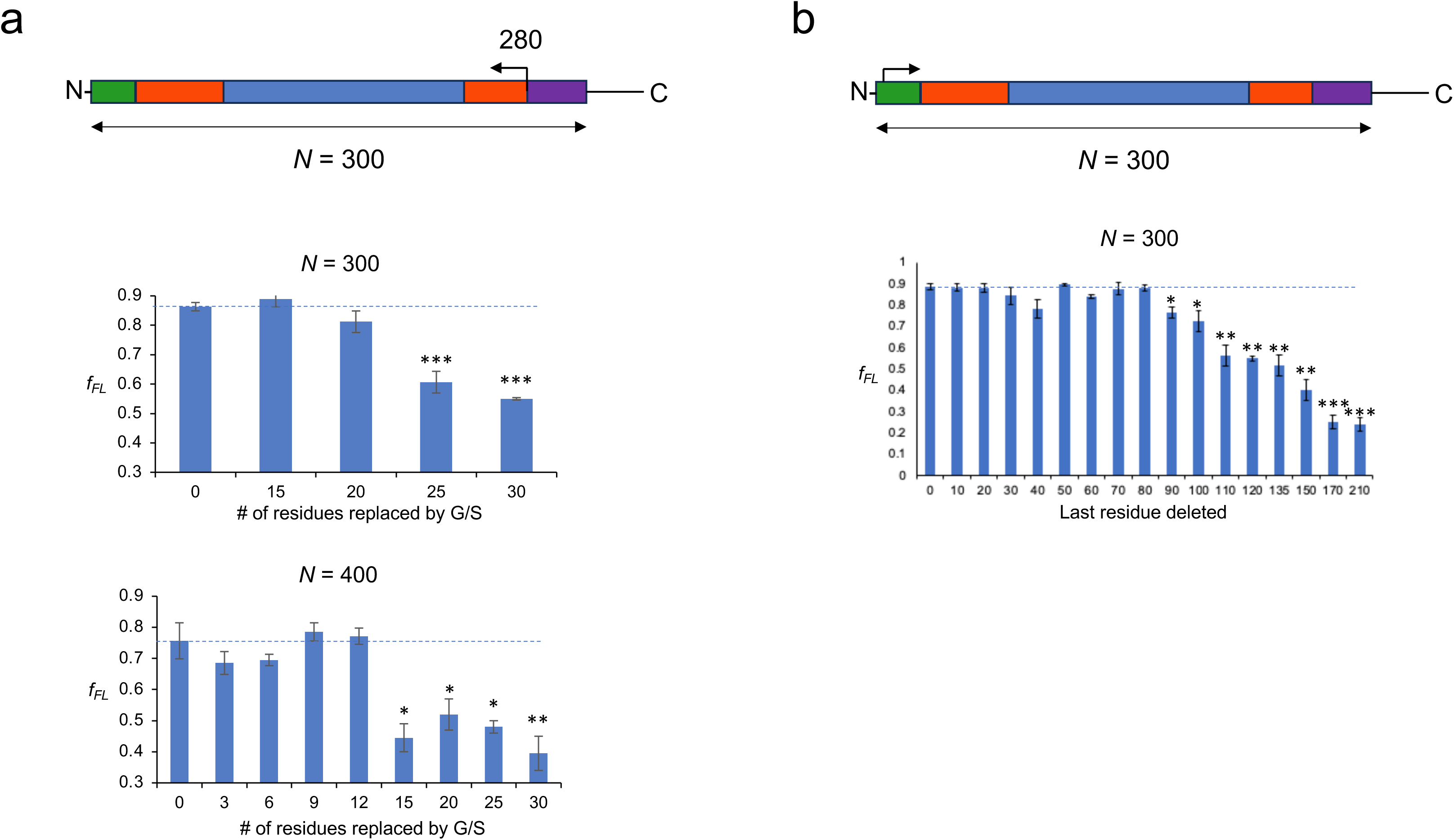
GS- and deletion-mapping of the N = 300 and N = 400 residues peaks in the FLuc force profile. (a) FLuc residues immediately upstream of the AP (purple) are progressively replaced in the C-to-N-terminal direction by alternating G and S residues, as indicated by the arrow. For the *N* = 300 residues construct, the replacements start at FLuc residue 280. GS-mapping results for the *N* = 300 and *N* = 400 residues peaks in the FLuc FP are shown below. Significant differences between the *f_FL_* values of the non-mutated and mutated constructs are indicated ((*p* < 0.05 (*), *p* < 0.01 (**), *p* < 0.001 (***); calculated by Student’s t-test). Error bars indicate the standard error of the mean (SEM). (b) N-terminal residues (starting at FLuc residue Ala^4^) are progressively deleted in the N-to-C-terminal direction, as indicated by the arrow. N-terminal deletion results for the N = 300 residues peak in the FLuc FP are shown below. Significant differences between the *f_FL_* values of the original and the deletion constructs are indicated (*p* < 0.05 (*), *p* < 0.01 (**), *p* < 0.001 (***); calculated by Student’s t-test). Error bars indicate the standard error of the mean (SEM).

**Figure 5.**
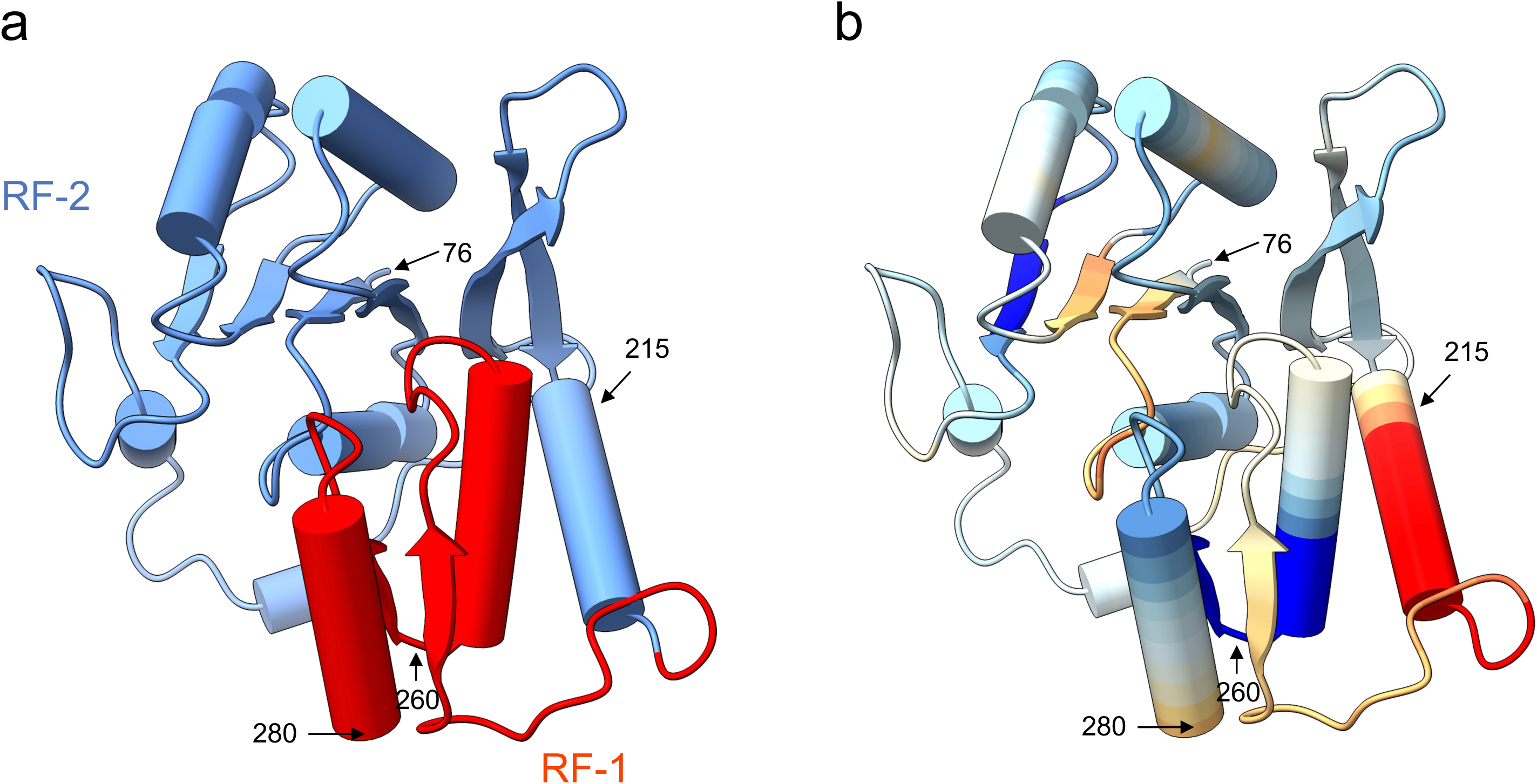
FLuc residues 76-280. (a) Colored by domain structure, c.f., Fig. 1. (b) Colored according to *f_FL_* values (low-to-high red-yellow-blue spectrum). The coloring of a given residue represents the instantaneous pulling force on the AP at the point when this residue is located 40 residues away from the PTC. Thus, the dip in the FP at *N* = 255-265 residues (Fig. 3) corresponds to the emergence of the α-helix starting at residue 215 (red), and the peak in the FP at *N* ≈ 300 residues corresponds to the emergence of the α-helix ending at *N* = 260 residues (deep blue), from the ET.

The gradual reduction in the peak amplitude seen for the progressively longer N-terminal deletions ending between residues 90 to 150 (Fig. 4b), corresponding to the consecutive removal of RF-2 strands β1, β2, β3, and β4 (see Fig. 5), is puzzling at first sight. However, full-atom MD simulations show that a fragment of the FLuc native structure that corresponds to residues 81-282 is quite stable during a 1 μs simulation, with the six-stranded β-sheet and the central core around the helix formed by residues 89-98 remaining intact, Supplemental Fig. S2 and Supplemental Movie M1. A fragment encompassing residues 111-282 (lacking β1, the core helix, and β2) adopts conformations in which the central β-sheet is broken into two 2-stranded β-sheets and the central core is extensively repacked because of the missing helix (residues 89-98), Supplemental Movie M2. The RMSD and RMSF values remain small, however, suggesting a fairly stable folded state, Fig. S2. In contrast, a fragment encompassing residues 151-282 (lacking most of the N-terminal half of the RF-2 domain) adopts rapidly interchanging conformations and is characterized by high RMSD and RMSF values, Fig. S2 and Supplemental Movie M3. Thus, the MD results are broadly consistent with the gradual drop in *f_FL_* values seen in Fig. 4b.

The peak at *N* = 400 residues and the deep dip at *N* = 410 residues suggest another folding intermediate. GS-mapping of the *N* = 400 residues construct shows that the C-terminal end of the parts that fold is ∼35 residues away from the PTC, Fig. 4a, i.e., close to the border between the RF-1 domain and the β-roll domain.

Finally, the deep dip at *N* = 468-472 residues corresponds to the full emergence of the core of the β-roll domain outside the ET, and perfectly matches the predicted folding intermediate at *N* = 460-470 residues, Supplemental Fig. S1. The absence of high peaks in the FP during the exit of the β-roll domain from the ET (*N* ≈ 410-475 residues; FLuc residues ∼370-435) suggests that it may not be as stably folded as the other subdomains, in agreement with results obtained by HDX-MS^22^.

Since, given its flexible attachment to the N-domain, it seemed likely that the CTD can fold by itself, we recorded the last part of the FP both for full-length FLuc and for a FLuc segment (residues 423-550) encompassing the isolated CTD. Notably, while the two FPs are identical within experimental error between *N* = 500-555 residues, the isolated CTD FP consistently has significantly higher *f_FL_* values (by about 0.2, on average) between *N* = 560-605 residues, while the shapes of the two FPs remain very similar, Supplemental Fig. S3a. Thus, the presence of the NTD causes a significant destabilization of the second half of the CTD in the ribosome-bound NC, again in agreement with HDX-MS data^22^.

### Cryo-EM analysis of FLuc RNCs

To further characterize the early part of the FP, we picked three constructs corresponding to the peaks at *N* = 110, 130, and 190 residues (Fig. 3) for analysis by cryo-EM. RNCs carrying these constructs stalled by the strong SecM(3W) AP^30^ were purified from *E. coli* cells, and strcutures were determined at an average resolution of 2.3 Å. For all three constructs, stalled NCs were readily apparent in the ET, with only small compact densities evident near the uL24 loop in the exit port, Fig. 6. The absence of more extensive folded domains in the ET shows that the FP peaks are not generated by the formation of local substructures within the ET.

**Figure 6.**
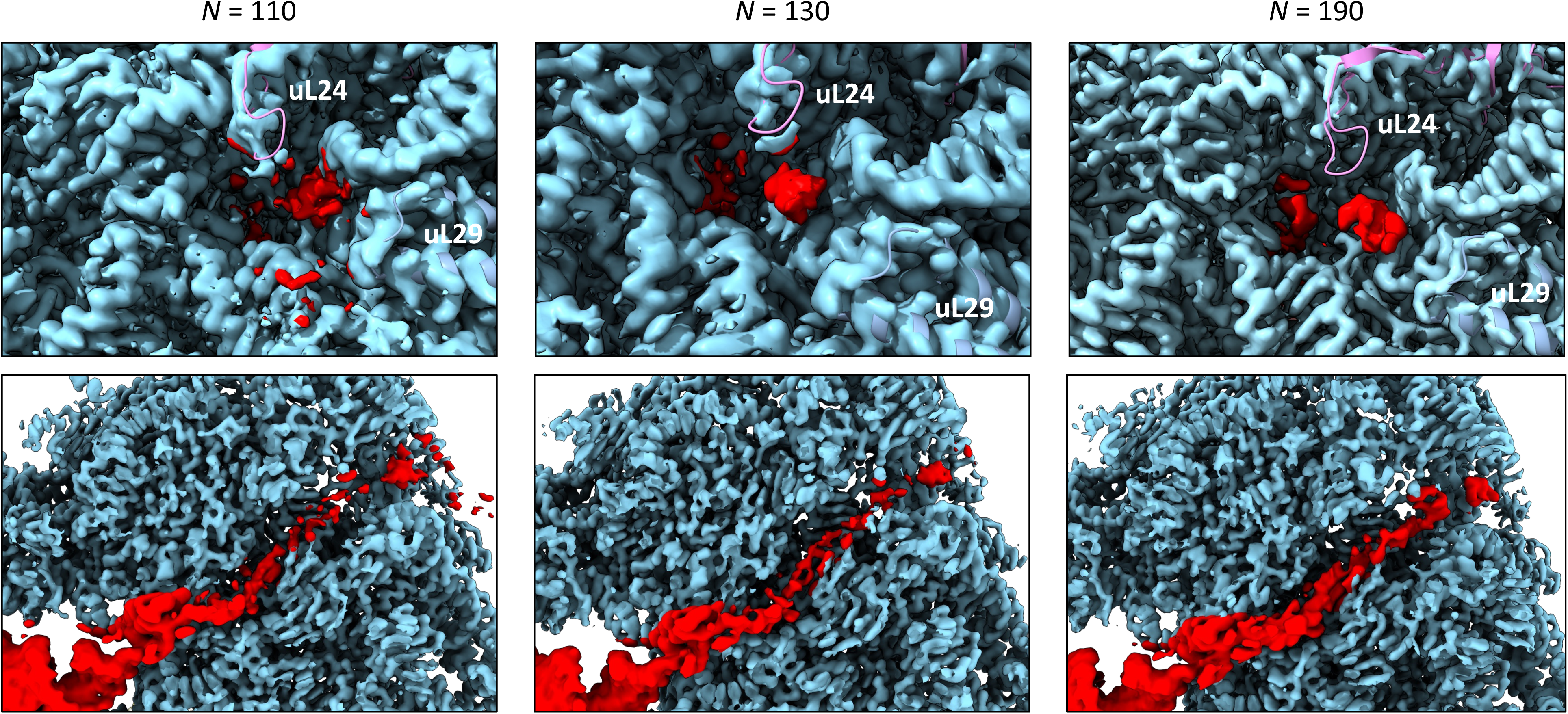
Cryo-EM densities of E. coli ribosomes with Fluc constructs N = 110, 130, and 190 residues stalled in the exit tunnel. Composite maps are shown with NC-density in the exit tunnel colored red. Ribosomal proteins uL24 and uL29 from PDB:3JBU (rigid body-docked) are shown in ribbon representation. Top panels: the ribosome exit port area, bottom panels: cross-section showing the NC-density (red) in the ET (part of the tRNA density is seen in the lower left corner).

### Effects of TF and TF[2xRBD] on the FLuc FP

In *E. coli*, Trigger Factor (TF) is the first chaperone that meets a NC as it emerges from the exit port^39^. NC-TF interactions can typically be detected when the NC is 80-120 residues long^40^. A host of studies have shown that TF preferentially binds compacted/partially folded substrates in a wide cavity located ∼50-100 Å away from the ribosome exit port^6,41–45^. FP analysis of the cotranslational folding of DHFR has shown that TF reduces the amplitude of the peak that represents the main folding event^46^. Given the multiple compaction/folding events apparent in the FLuc FP, we tested the effects of TF and of the TF[2xRBD] variant, in which duplication of the ribosome-binding domain positions the substrate-binding cavity farther from the exit port than in wild-type TF^6^.

The FLuc FP obtained in the presence of TF, Fig. 7 (red curve), is overall quite similar to the original FP but with some interesting differences. The most dramatic difference is for the ∼40-residue long region *N* = 165-205 residues, where the *f_FL_* values are markedly reduced (by about 0.25, on average) in the presence of TF, even though the shapes of the two FPs remain similar, Supplemental Fig. S3b. Significant but more localized reductions in *f_FL_* are seen at *N* ≈ 110, 250, 300, 400, and 525 residues, and significant increases in *f_FL_* at *N* ≈ 125 and 365 residues. The apparent reduction at *N* = 135-145 is not statistically significant on the *p* < 0.01 level.

**Figure 7.**
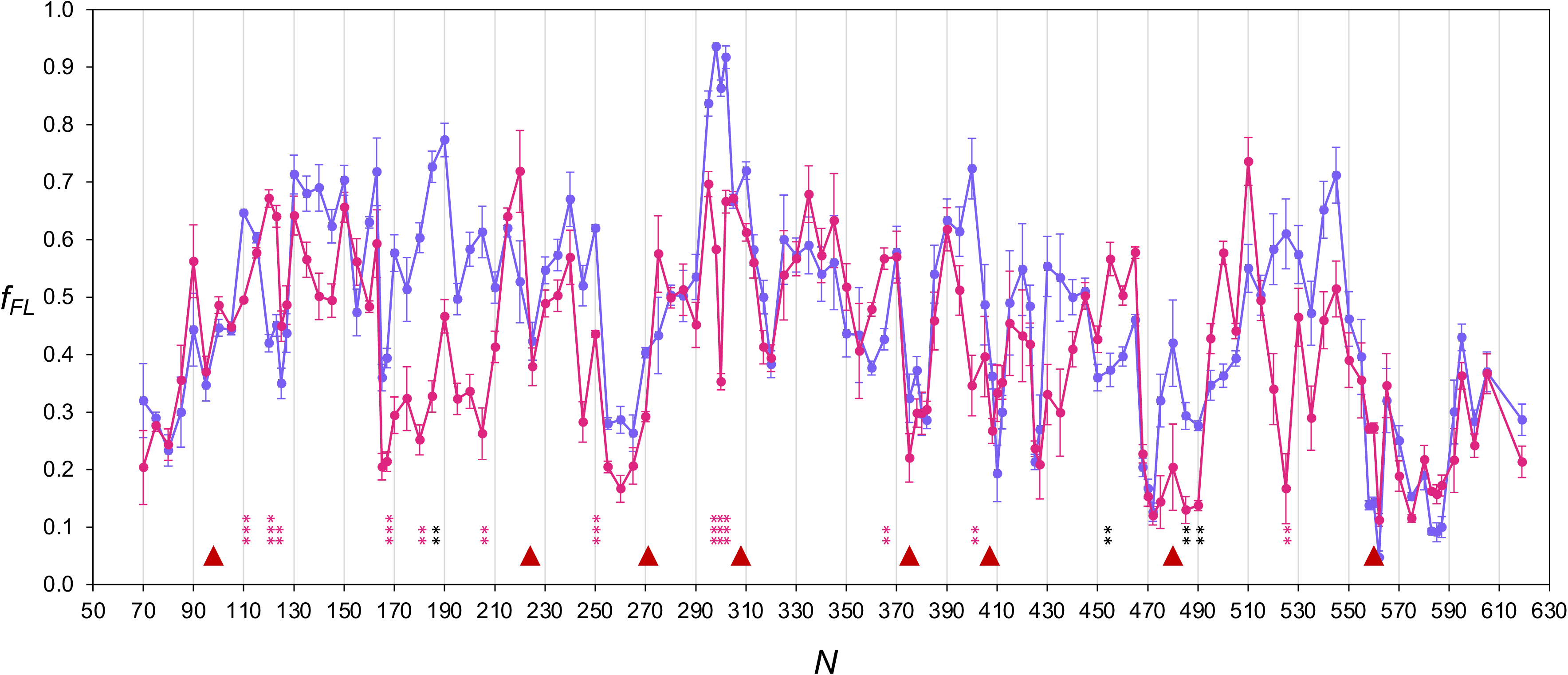
Effects of trigger factor on the FLuc force profile. FLuc FP obtained in the presence of 4 μM TF (dark red), superimposed on the original FP from Fig. 3 (blue). Constructs for which the difference between the ±TF *f_FL_* values are significant on the *p* < 0.01 (**) or *p* < 0.001 (***) levels (calculated by Student’s t-test) are indicated. When marked in red, the differences are also significant on the *p* < 0.05 level according to the Holm-Bonferroni method^60^, see Materials and Methods. Error bars indicate the standard error of the mean (SEM).

The partial FP recorded in the presence of TF[2xRBD] (which we focused on areas where there are significant TF effects), Supplemental Fig. S4, is even more similar to the original FP, and the dramatic TF effects seen at *N* = 165-205 are not seen with TF[2xRBD]. Notably, however, the amplitudes of the main peaks at *N* = 300, 400, and 525 residues are markedly reduced by TF[2xRBD], albeit the effects are smaller than seen with TF.

Interpretation of the FLuc-TF and FLuc-TF[2xRBD] FPs is complicated by the fact that the extended TF binding cavity is located far from the ribosome exit port where the compaction/folding events that are reflected in the FP occur. Depending on where a particular substrate domain binds in the binding cavity, a minimum of ∼15-20 NC residues in an extended conformation would be required to reach from the exit port to the TF-binding cavity^47,48^. Indeed, crosslinking analysis of interactions between folding intermediates in *E. coli* β-galactosidase NCs and TF show that the number of NC residues between the PTC and the most C-terminal NC residue found to crosslink to TF typically ranges from ∼80 up to ∼150 residues but can be as short as ∼60 residues^6^, in agreement with ribosome profiling data^40^.

Thus, the *f_FL_* value recorded at a given *N* value in the presence of TF should reflect the combined effects of a TF-binding event happening at least ∼20 residues away from the exit port and simultaneous folding/compaction events taking place at the exit port. Previous fluorescence-based studies have shown that a ribosome-stalled FLuc translation intermediate of length 77 residues (comparable to our *N* = 77 construct) does not interact with TF, while a translation intermediate of length 164 residues (comparable to our *N* = 164 construct) both recruits TF to the ribosome and interacts with the binding cavity, albeit considerably less efficiently than a long, 520-residue intermediate^49^. This fits nicely with the TF-induced reduction in *f_FL_* values starting at *N* ≈ 165 residues, strongly suggesting that binding in the TF cavity competes with (but, because the FP shape stays the same, does not completely eliminate) compaction/folding at the exit port between *N* ≈ 165-210 residues.

Assuming that 40 residues are sequestered in the ET, a possible interpretation of the early parts of the +TF FP is that compaction at the exit port commences no later than when ∼90 residues have emerged from the ET (*N* ≈ 130 residues), that the compacting NC can reach the TF-binding cavity when ∼125 residues have emerged from the ET (*N* = 165 residues) at which point TF-binding starts to compete with continued compaction at the exit port (thus reducing *f_FL_*), and that formation of a more stable compacted state at the exit port then outcompetes TF-binding, starting at *N* ≈ 210 residues. The compacted part increases in size until the first main compacted intermediate, encompassing residues ∼1-210, is complete at *N* ≈ 250 residues, Fig. 7. Given recent data demonstrating that TF binds compacted/partially folded rather than unfolded regions in a NC^6^, the TF-induced reduction in *f_FL_* values starting at *N* = 165 residues is a strong indication that the FLuc NC forms a compacted intermediate no later than when ∼125 residues have emerged from the ET. This further implies that the intermediate-amplitude region in the -TF FP from *N* = 130-250 residues reflects a continuous compaction/folding process, rather than an abrupt, cooperative folding event. A similar scenario might apply for the *N* ≈ 520-545 residues region in the CTD.

The strong, narrowly localized reductions in *f_FL_* caused by TF at the maxima of the major peaks at *N* ≈ 300 and 400 residues are seen also with TF[2xRBD], which is surprising since the binding cavity is at different distances away from the exit port in TF and TF[2xRBD]. A possible way out of this conundrum is if these TF effects are caused by interactions between the NC and the TF RBD rather than the central TF cavity; some evidence for the existence of a NC binding site in the RBD has recently been obtained by chemical crosslinking^6^.

## Discussion

Cotranslational folding of multi-domain proteins has so far been studied mainly by biophysical methods such as NMR^5^, HDX-MS^22^, and optical tweezers^50^. Data obtained with these kinds of methods often have rather straight-forward structural interpretations, but are laborious to obtain and hence normally derived from the analysis of a limited number of stalled ribosome-nascent chain complexes (RNCs) for any given target protein.

Force-profile analysis (FPA) allows more fine-grained study of cotranslational folding—in principle with up to single-residue resolution—but has not yet been applied to multi-domain proteins, except in a high-throughput fluorescence-based *in vivo* format^51^. Here, we have applied the “classical”, gel-based FPA method^3,27^ to map the cotranslational folding of the 550-residue multi-domain protein FLuc at five-residue resolution, both in the absence and presence of the TF chaperone. FLuc has an interesting architecture, with a relatively large NTD flexibly attached to a relatively small CTD. The NTD in turn is composed of two intertwined Rossmann folds and a β-roll domain, and it has been unclear to what extent identifiable folding intermediates, in addition to the semi-independent NTD and CTD, can form during translation.

Overall, the FLuc FP (Fig. 3) is characterized by rather long regions of fluctuating *f_FL_* values of intermediate amplitude, punctuated by one major peak at N ≈ 300 residues and a few deep dips that we propose represent the formation of transient folding intermediates.

The early part of the FP, up to *N* = 250 residues, shows an intricate pattern of medium-amplitude peaks and dips, quite distinct from the characteristic high-amplitude, sharp peaks previously seen for cooperatively folding of single-domain proteins^3,10,32,52^. This region also shows clear signs of TF-binding (Fig. 7), suggesting the formation of a compacted/partially folded NC. This in agreement with HDX-MS data for the ^1/2^N_s_ and N_s_ RNCs (corresponding to our *N* = 145 and *N* = 230 constructs)^22^, and with the general observation that on-ribosome folding promotes formation of partially folded intermediates^12^.

As shown by N-terminal deletion and C-terminal GS-replacement analysis, the main peak at *N* ≈ 300 residues signals a cooperative folding event involving FLuc residues ∼80-260, precisely matching the RF-2 domain plus an additional helix located at the border between the RF-2 domain and the second half of the RF-1 domain (Fig. 5). Interestingly, deletion of the first half of the RF-1 domain, which most likely is also folded at *N* ≈ 300 residues, has no effect on the amplitude of the *N* ≈ 300 peak. In itself, this part of the RF-1 domain therefore does not seem to contribute much to the stability of the RF-2 domain.

The intermediate amplitude regions in the FP imply an almost continuous series of small compaction/folding events during chain elongation. The absence of distinct peaks and deep dips in these regions suggests that these events are non-cooperative, possibly reflecting ongoing compaction and/or the addition of small pieces of structure to a more or less stably folded core, each piece providing only a marginal contribution to the overall stability.

The effect of TF on the FP is broadly consistent with this picture of the cotranslational folding process, especially in the region *N* = 165-210 residues. The large reduction in *f_FL_* over this region implies a strong interaction with TF, in turn implying that the NC is compacted/partially folded (as also shown by the HDX-MS data^22^ for the N_S_ construct). Nevertheless, for *N* ≥ 215 residues, compaction at the exit port apparently outcompetes TF-binding. A similar pattern is seen for the CTD, where TF markedly reduces *f_FL_* for the region *N* = 520-545 residues, presumably by binding to the early parts of the CTD and thereby preventing folding at the exit port. In addition, the FP data has uncovered narrowly localized effects of TF on two of the main peaks in the FP, at *N* = 300 and 400 residues. These effects are also seen with TF[2xRBD], hinting at a possible involvement of the TF RBD. The overall cotranslational compaction/folding process is summarized schematically in Fig. 8.

**Figure 8.**
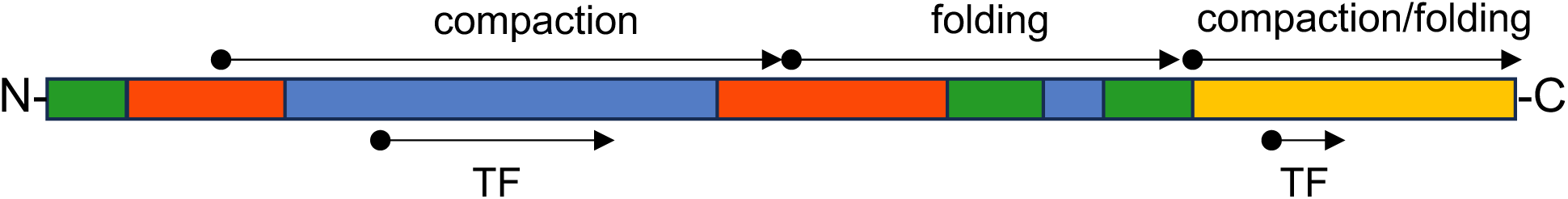
Schematic model for cotranslational folding of FLuc. Compaction of the NC begins when FLuc residue ∼70 emerges from the exit port at *N* ≈ 110 residues (c.f., Fig. 3). Folding into the native structure initiates when residue ∼260, close to the end of the RF-2 domain, emerges at *N* ≈ 300 residues. The CTD undergoes a similar compaction/folding process as it emerges from the exit port. FLuc starts to interact with the TF chaperone when ∼125 residues have emerged outside the ribosome (at *N* ≈ 165 residues).

While our analysis here has focused on the main peaks and dips in the FLuc FP, we surmise that the intricate medium-amplitude features seen throughout the FP also hold important information on the compaction and folding of the growing NC. Single-residue resolution FP studies, together with mutational analysis and computational modelling, may eventually make it possible to gain a detailed, residue-level understanding of both the initial compaction/partial folding processes evident in the first half of the FLuc FP and the later growth of the folded domains in the second half. Promising steps in this direction have already been taken for the small single-domain protein HemK-NTD^53^.

## Online Methods

### Enzymes and chemicals

Cloning enzymes, reagents, and buffers were acquired from New England BioLabs (Ipswich, Massachusetts, USA) and Thermo Fisher Scientific (Waltham, Massachusetts, USA). Oligonucleotides were acquired from Eurofins (Germany). [^35^S] L-Methionine was acquired from Revvity (Waltham, MA, USA). All other chemicals were acquired from Sigma-Aldrich (subsidiary of Merck Life Science, Darmstadt, Germany) unless otherwise stated.

### Purification of E. coli Trigger Factor

TF was expressed from the plasmid pCA528^54^ containing an Ulp1-cleavable N-terminal His_6_-SUMO tag. Protein expression and purification was carried out by the Protein Science Facility at Karolinska Institutet, Stockholm, Sweden, as described previously^55^.

### Cloning and mutagenesis

All FLuc plasmids were produced by Invitrogen GeneArt Services, Thermo Fisher Scientific. The full-length pET19b plasmid contains—after modification—the 550-residue long FLuc gene originated from *Photinus pyralis* (UniProt Q27758), a 52-residue GS-linker sequence, the *E. coli* SecM AP FSTPVWISQAQGIRAGP, and an extra C-terminal tail derived from the *lepB* gene, under the control of a T7 promoter^23,26,27,56^. Stepwise internal deletions upstream of the AP resulted in 109 truncated constructs that differ in length by 5 amino acids. Further mutagenesis was performed in-house by a combination of Q5 PCR and Gibson assembly^57^ with a homemade reaction mixture.

### In vitro transcription and translation

The FLuc-encoding regions, including a 5’ T7 promoter, were individually amplified by PCR from the plasmid set. PCR products were purified and subjected to *in vitro* transcription and translation according to the PURExpress^TM^ protocol from New England Biolabs. The PURExpress^TM^ kit contains the essential components required for *in vitro* transcription and translation isolated from *E. coli*. Wild-type and mutant Trigger Factor proteins (TF, TF[2xRBD])^6,58^ were prepared at 40 μM in PURE system buffer (9 mM magnesium acetate, 5 mM potassium phosphate pH 7.3, 95 mM potassium glutamate, 5 mM NH4Cl, 1 mM spermidine, 8 mM putrescine, 1 mM DTT) and included in the *in vitro* reactions at a final concentration of 4 μM. After addition of 0.8 μL [^35^S]-Methionine, PURE reactions were incubated for 15 min at 37 °C with shaking at 700 rpm in an Eppendorf Thermomixer. The reaction was stopped by the addition of 10% ice-cold TCA, and samples were incubated for 30 min on ice and then centrifuged for 5 min at 20,800 x g at 4 °C. The pellets were dissolved in sample buffer (67 mM Tris, 3.3% (w/v) SDS, 0.012% (w/v) bromophenol blue, 10mM EDTA-KOH pH 8.0, 6.75% (v/v) glycerol, and 100 mM DTT) and treated with 0.25 mg/mL RNase at 37 °C for 15 min. The mixture was then separated by SDS/PAGE and the radiolabeled proteins were detected using a Fuji FLA-3000 scanner.

### Quantification

Full-length and arrested band intensity profiles were extracted with ImageJ^59^. The profiles were then fit to one or more Gaussian curves using an in-house python script to determine the integrated band intensities. *f*_FL_ was calculated as *f_FL_* = *I_FL_*/(*I_FL_*+*I_A_*), where *I_A_* and *I_FL_* are the integrated intensities of the arrested and full-length bands, respectively. For certain constructs where band assignment was ambiguous, control constructs were cloned and expressed. Full-length control constructs were made by mutating the C-terminal Pro in SecM(*Ec*) to Ala, which prevents SecM(*Ec*)-induced stalling. Arrested controls were made by mutating the C-terminal Pro in the SecM(*Ec*) to a stop codon.

### Statistics

To correct for multiple testing and correlations in the FP when comparing the FLuc[±TF] FPs (Fig. 7), the autocorrelation function (ACF) was first calculated for the FLuc[-TF] FP (109 data points; only data points for *N-*values that are multiples of 5 were included). The ACF showed significant values for lags 1-3 (*i.e*., for residue separations up to 15 residues), Supplemental Fig. S5. The Holm-Bonferroni method^60^ was then used to identify the *N*-values for which the FLuc[±TF] *f_FL_* values differed on the α = 0.05 level, with *p*-values calculated by Student’s t-test and the number of independent tests set to *m* = 109/4 = 27.

### Molecular dynamics simulations

Three models, Δ4-80, Δ4-110, and Δ4-150, were created based on the AlphaFold DB^61^ model AF-P08659-F1-v4. The model structure was trimmed, and the N-terminal modified to match the sequences shown below:

Δ4-80:

MEDCSENSLQFFMPVLGALFIGVAVAPANDIYNERELLNSMGISQPTVVFVSKKGLQKILNVQKKLPIIQKIIIMDSKTDYQGFQSMYTFVTSHLPPGFNEYDFVPESFDRDKTIALIMNSSGSTGLPKGVALPHRTACVRFSHARDPIFGNQIIPDTAILSVVPFHHGFGMFTTLGYLICGFRVVLMYRFEEELFLRSLQDY

Δ4-110:

MEDERELLNSMGISQPTVVFVSKKGLQKILNVQKKLPIIQKIIIMDSKTDYQGFQSMYTFVTSHLPPGFNEYDFVPESFDRDKTIALIMNSSGSTGLPKGVALPHRTACVRFSHARDPIFGNQIIPDTAILSVVPFHHGFGMFTTLGYLICGFRVVLMYRFEEELFLRSLQDY

Δ4-150

MEDIMDSKTDYQGFQSMYTFVTSHLPPGFNEYDFVPESFDRDKTIALIMNSSGSTGLPKGVALPHRTACVRFSHARDPIFGNQIIPDTAILSVVPFHHGFGMFTTLGYLICGFRVVLMYRFEEELFLRSLQDY

For all three models, the residues at the N-terminus were mutated using PyMOL^62^ to match the “MED” sequence. The terminal residues were capped with ACE (N-acetyl) and NME (N-methylamide).

Molecular dynamics simulations were performed using the Amber99SB-ILDN force field^63^ with GROMACS 2024.4^64^. Each protein was solvated in a dodecahedron box of explicit TIP3P^65^ water, with a 1.5 nm distance from the solute to the edge of the box. The systems were neutralized with Na^+^ or Cl^-^ ions.

The systems were energy minimized using steepest descent and then subjected to the equilibration protocol described below:

0–500 ps: NVT and position restraints on all heavy atoms (force constant of 1000 kJ mol^−1^nm^−2^), temperature coupling to 310 K using V-rescale thermostat^66^.

500ps–1 ns: NPT with a Berendsen barostat^67^ with a coupling constant τp = 0.5 ps and an isotropic compressibility of 4.5·10^−5^bar^−1^, temperature coupling to 310 K using V-rescale thermostat.

After equilibration, for each system, 3 simulations of 1 μs each were performed in the NPT ensemble, with periodic boundary conditions. Temperature coupling was done with the Nose-Hoover thermostat (310 K)^68^, and pressure coupling was done with the Parrinello-Rahman barostat (τp = 5ps and an isotropic compressibility of 4.5·10^−5^bar^−1^)^69^. A 10 Å cut-off was used for van der Waals and short-range electrostatic interactions. The Particle-Mesh Ewald (PME) summation method was used for long-range electrostatic interactions^70^. Verlet cut-off scheme was used^71^.Covalent bonds were constrained using the LINCS algorithm^72^. The integration time step was 2 fs for all steps.

To analyze the simulations, trajectories were filtered by extracting a frame every 100 ps and the root mean square fluctuation (RMSF) and root mean square deviation (RMSD) relative to an average conformation for the simulation period between 0.1 and 1 μs were calculated, as described below:

The RMSF of the C_α_ atoms of each frame in the filtered trajectories, relative to the average structure, was computed using the “gmx rmsf” function in GROMACS. The average conformation was calculated from the sampled conformations between 0.1 and 1 μs for each trajectory. The average structure for each trajectory was saved as a PDB file, and the mean RMSF value and standard deviation for each C_α_ atom was calculated for each trajectory.

The root mean squared distance (RMSD) to the average structure (obtained during the RMSF calculation) was calculated using the “gmx rms” function in GROMACS and fitting for both rotation and translation. To create the RMSD plot, the “histplot” function implemented in seaborn^73^ was used to calculate the distributions from the three trajectories for each system.

Both plots were generated in Python 3.10 with matplotlib^74^ and seaborn^73^. Structure images were created using ChimeraX^75^. Movies were generated using VMD^76^.

### Cryo-electron microscopy

Constructs from *in vitro* transcription and translation experiments were modified for cryo-electron microscopy (cryo-EM). First, the SecM(3W) AP (FSTPVWIWWWPPIRGSP) was used to enhance stalling^30^. Second, a 6xHis tag was added to the N-terminus of FLuc to facilitate purification of RNCs (full sequences in Supplementary Table S1). Specifically, we chose three constructs at *N* = 110, 130, and 190 residues. RNCs were purified according to^77^, and expression from pET19b was induced with 1 mM IPTG in *E.coli* BL21(DE3). Purified RNCs were added to grids using a Vitrobot (Thermo Fisher Scientific, Waltham, Massachusetts, USA), plunge-freezing with a 3 second blot time and 15 second wait time in 100% humidity at 4°C. Data were collected on a Krios G3i microscope (Thermo Fisher Scientific, Waltham, Massachusetts, USA) equipped with a Gatan K3 DED at 300 kV with a pixel size of 0.825 Å/pixel, over 40 frames with a total dose of 40 e^−^/Å^2^. Processing was performed using CryoSPARC 4.3.0. Briefly, after motion correction and CTF estimation, particles were picked using either the built-in “blob” picker or based on templates generated from the data. After 2D classification, selected particles were subjected to 3D homogeneous refinement. Refined particles were then subtracted such that the only density remaining was of the ribosomal groove containing the tRNA. Then, particles were 3D classified based on the tRNA conformation. Finally, particles representing ribosomes with P-site tRNA were selected and refined to generate the final maps. There were slight processing differences between the three constructs. Full workflows are provided in Supplementary Fig. S6. The data was collected at the Swedish National Cryo-EM Facility funded by the Knut and Alice Wallenberg, Family Erling Persson and Kempe Foundations, SciLifeLab, Stockholm University and Umeå University.

## Author Contributions

Conceptualization, A.M., J.W., and G.v.H.; investigation, A.M. S.M., J.W., and F.P.A.; validation, A.M., J.W., and G.v.H.; formal analysis, A.M., J.W., F.P.A., and G.v.H.; visualization, A.M., S.M., J.W., F.P.A., and G.v.H.; supervision, A.M., M.L., and G.v.H.; funding acquisition, M.L., G.v.H.; writing – original draft, A.M. S.M., J.W., F.P.A., M.L., and G.v.H.; writing – review & editing, A.M., S.M., J.W., F.P.A., M.L., and G.v.H.

## Data availability

Cryo-EM maps have been deposited in the Electron Microscopy Data Bank (EMDB) under accession numbers EMD-56368, EMD-56453, and EMD-56462.

## Acknowledgments

This work was supported by grants from the Knut and Alice Wallenberg Foundation (2022.0241), the Novo Nordisk Fund (NNF18OC0032828), and the Swedish Research Council (2025-03943) to GvH, from the Stanford Data Science Initiative to FPA, and from the National Institute of Health (R35 GM122543) to ML. The plasmid encoding TF was a gift from Dr. Günter Kramer, Heidelberg. The FLuc gene and purified TF[2xRBD] were gifts from Dr David Balchin, London. We thank Dr. Mathieu Coinçon at the Swedish National Cryo-EM Facility for help with data collection, and Drs. Grant Pellowe and David Balchin, London, for discussions and sharing data prior to publication.

---

**Supplemental Figure S1.**
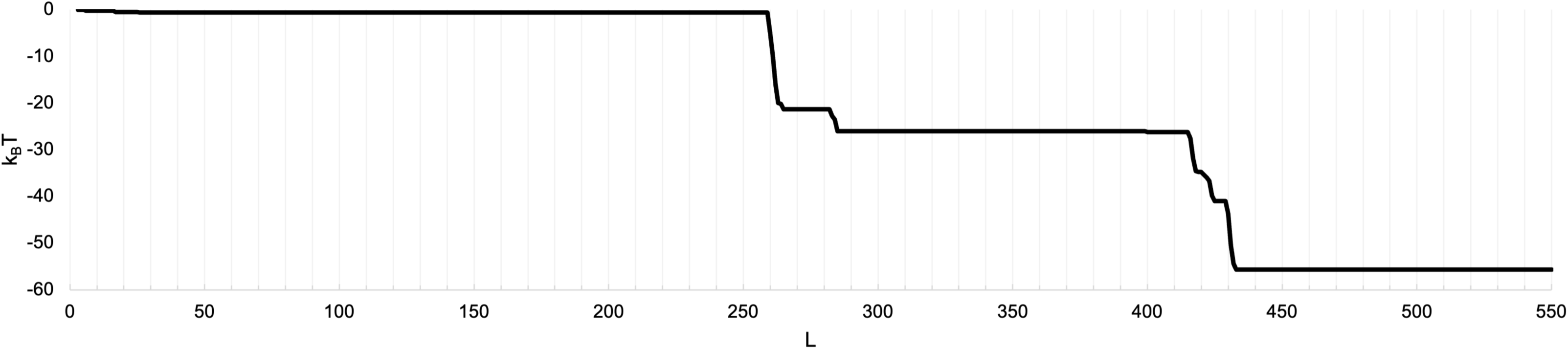
Cotranslational folding intermediates predicted by the Jacobs & Shakhnovic method^36^, using the full-length FLuc sequence (Supplemental Table S1) and the two PDB structures 5DV9 and 5WYS as input. Folding intermediates are predicted to form at FLuc lengths *L* ≈ 260-285, and 415-430 residues, corresponding to *N* ≈ 300-325 and 455-470 residues.

**Supplemental Figure S2.**
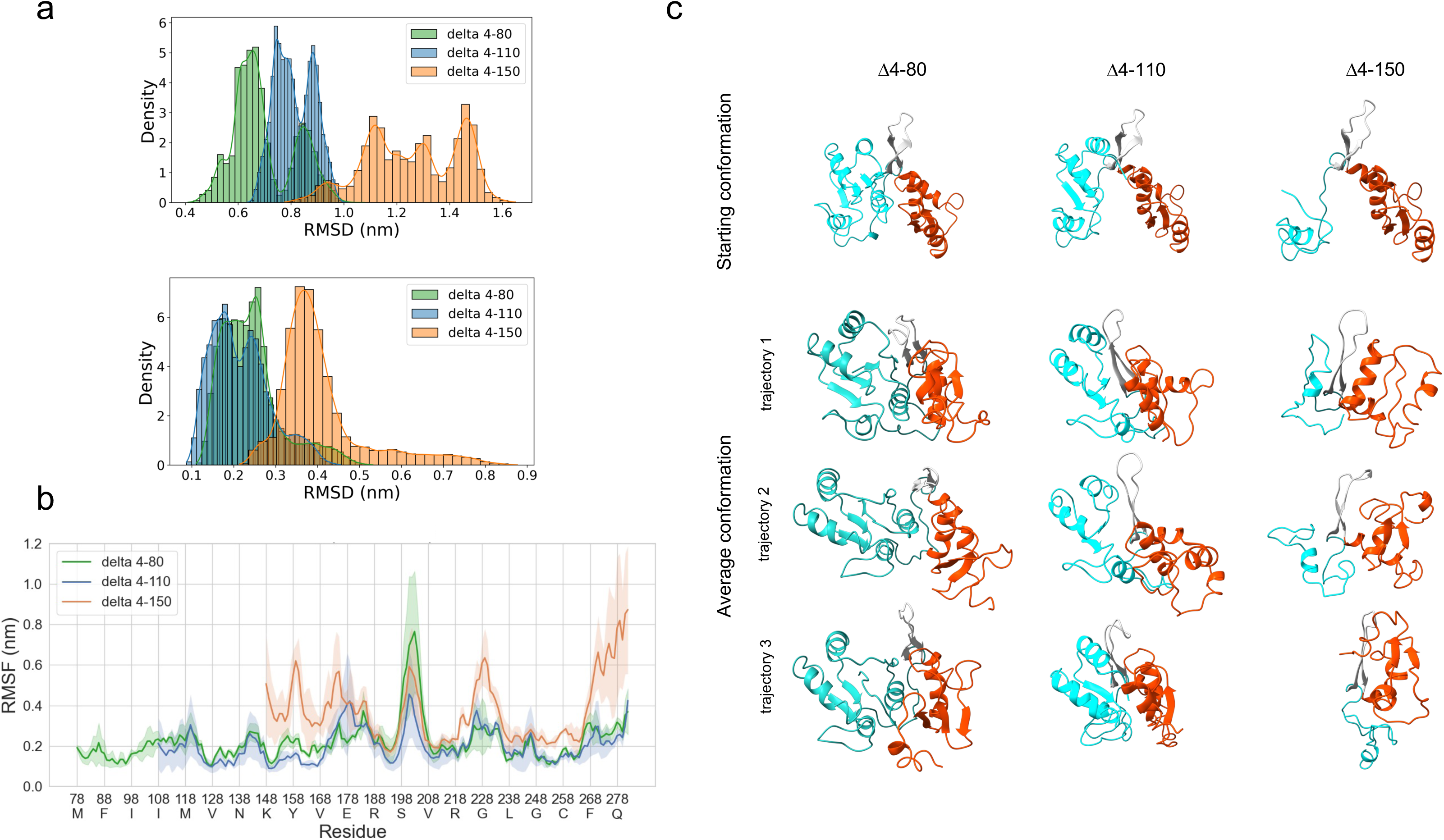
Full-atom molecular dynamics simulations of FLuc residues 1-282, with residues 4-80, 4-110, and 4-150 deleted. Three 1 μs simulations were performed for each of the three FLuc fragments. (a) RMSD distributions for the Δ4-80 (green), Δ4-110 (blue), and Δ4-150 (orange) FLuc fragments. RMSD values were calculated for each 100 ps simulation frame in the interval 0.1-1 μs using either the starting conformation (top panel) or the average conformation of all frames in the same interval (bottom; c.f. panel *c*) as the reference. The results for the three simulations for each fragment were pooled. (b) RMSF profiles for the Δ4-80 (green), Δ4-110 (blue), and Δ4-150 (orange) FLuc fragments. C_α_ RMSF values were calculated for each 100 ps simulation frame in the interval 0.1-1 μs using the average conformation of all frames in the same interval as the reference, and the results for the three simulations performed for each FLuc fragment were averaged (solid line). The shaded area indicates the standard deviation. (c) Starting conformation (initial models after energy minimization, c.f., Fig. 5) and the three average conformations (calculated over the interval 0.1-1 μs) for the Δ4-80, Δ4-110, and Δ4-150 FLuc simulations. Structures are colored as in Supplementary Movies M1-M3 (N-terminal part in cyan, β-hairpin residues 192-211 in gray, C-terminal part in red). Starting conformations were obtained by deleting the relevant residues from the complete FLuc structure (AF-P08659-F1-model_v4), followed by a steepest-descent energy minimization step.

**Supplemental Figure S3.**
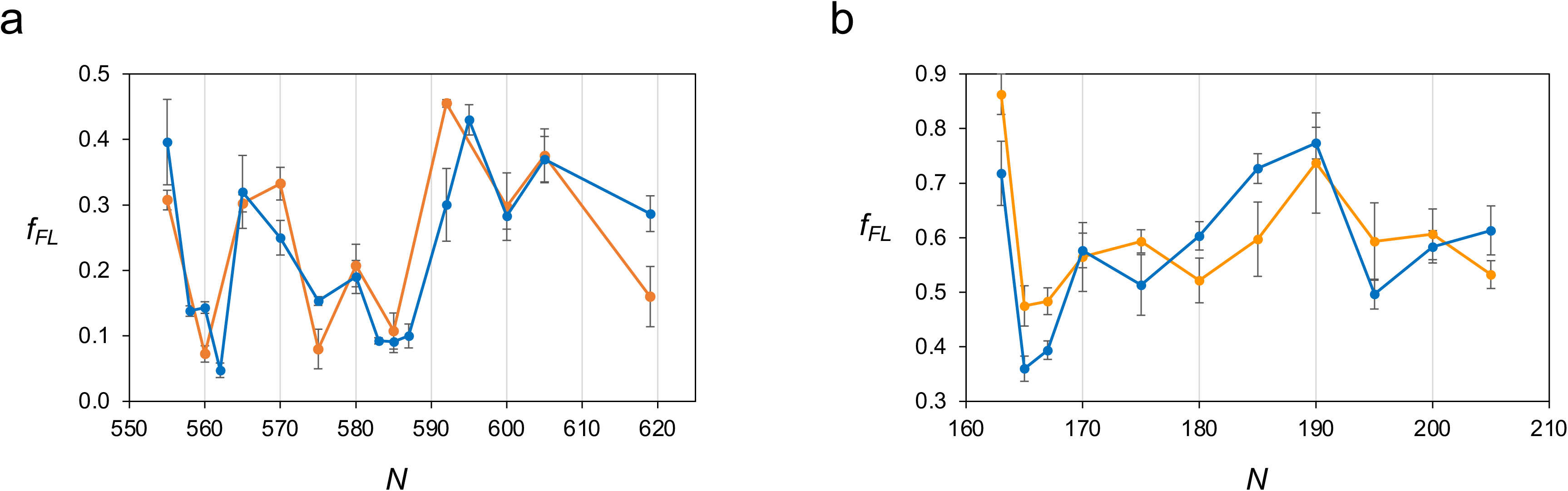
(a) Shape overlap between the FLuc FP (blue) and the CTD FP (orange) in the region *N* = 555-619 residues. The *f_FL_* values for the latter have been reduced by 0.2 compared to Fig. 3. Error bars indicate the standard error of the mean (SEM). (b) Shape overlap between the FLuc FP (blue) and the FLuc+TF FP (orange) in the region *N* = 163-205 residues. The *f_FL_* values for the latter have been increased by 0.27 compared to Fig. 7. Error bars indicate the standard error of the mean (SEM).

**Supplemental Figure S4.**
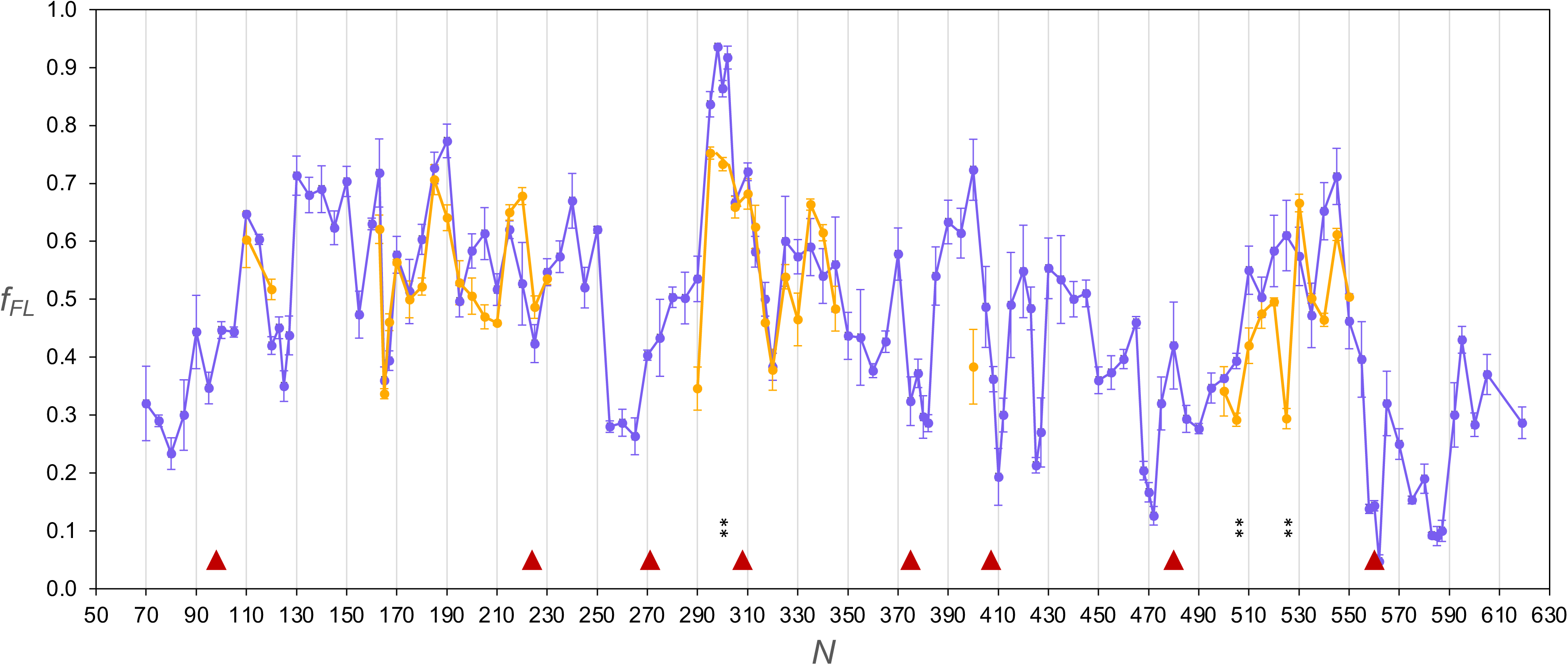
Effects of TF[2xRBD] on the FLuc force profile. FLuc FP obtained in the presence of 4 μM TF[2xRBD] (orange), superimposed on the original FP from Fig. 3 (blue). Constructs for which the difference between the ±TF[2xRBD] *f_FL_* values are significant on the *p* < 0.01 (**) level (calculated by Student’s t-test) are indicated. Error bars indicate the standard error of the mean (SEM).

**Supplemental Figure S5.**
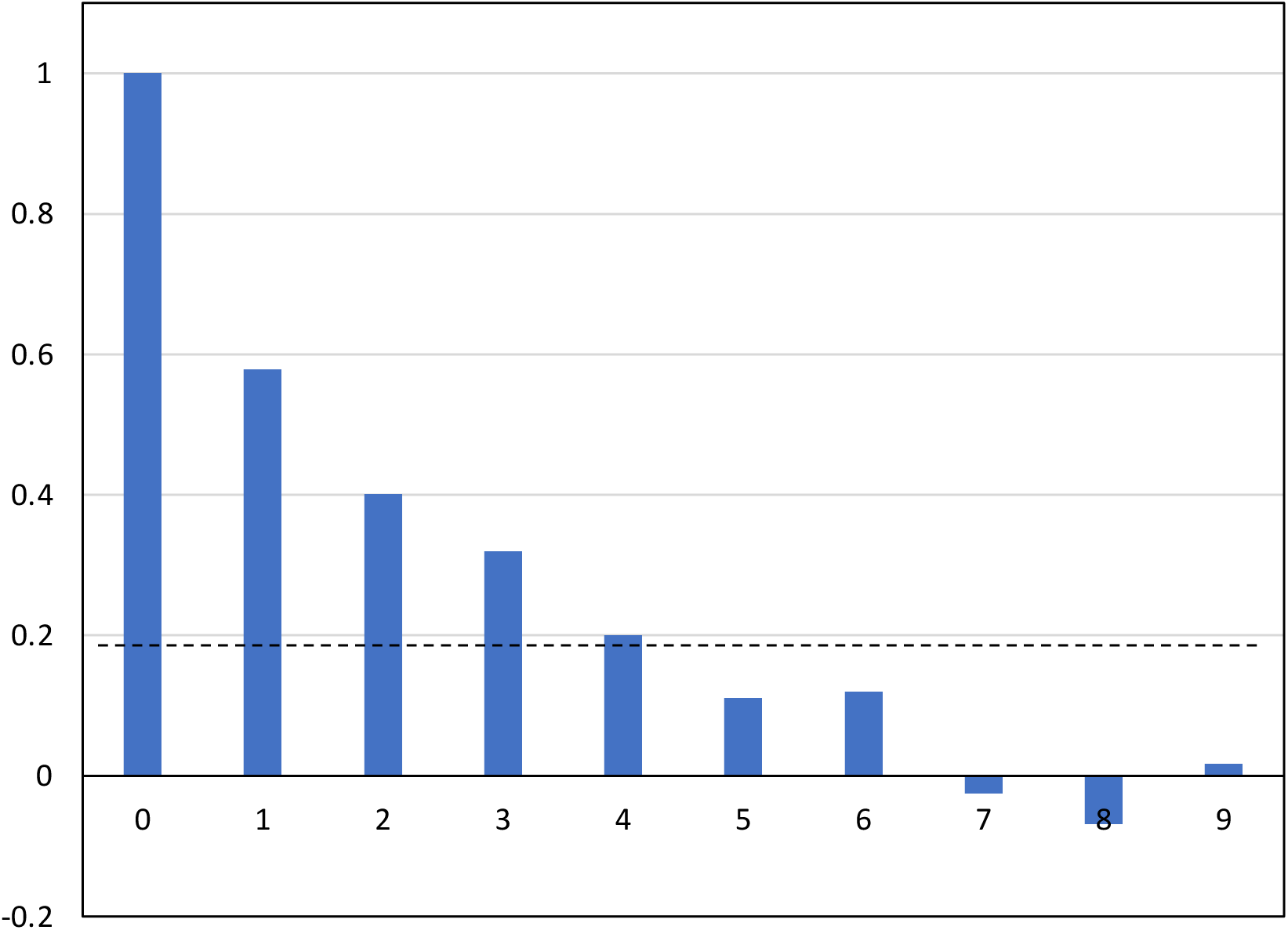
Autocorrelation Function calculated for the FLuc FP in Fig. 3 (see Materials and Methods). The dotted line indicates the confidence interval (±2/√%, where *n* = 109).

**Supplemental Figure S6.**
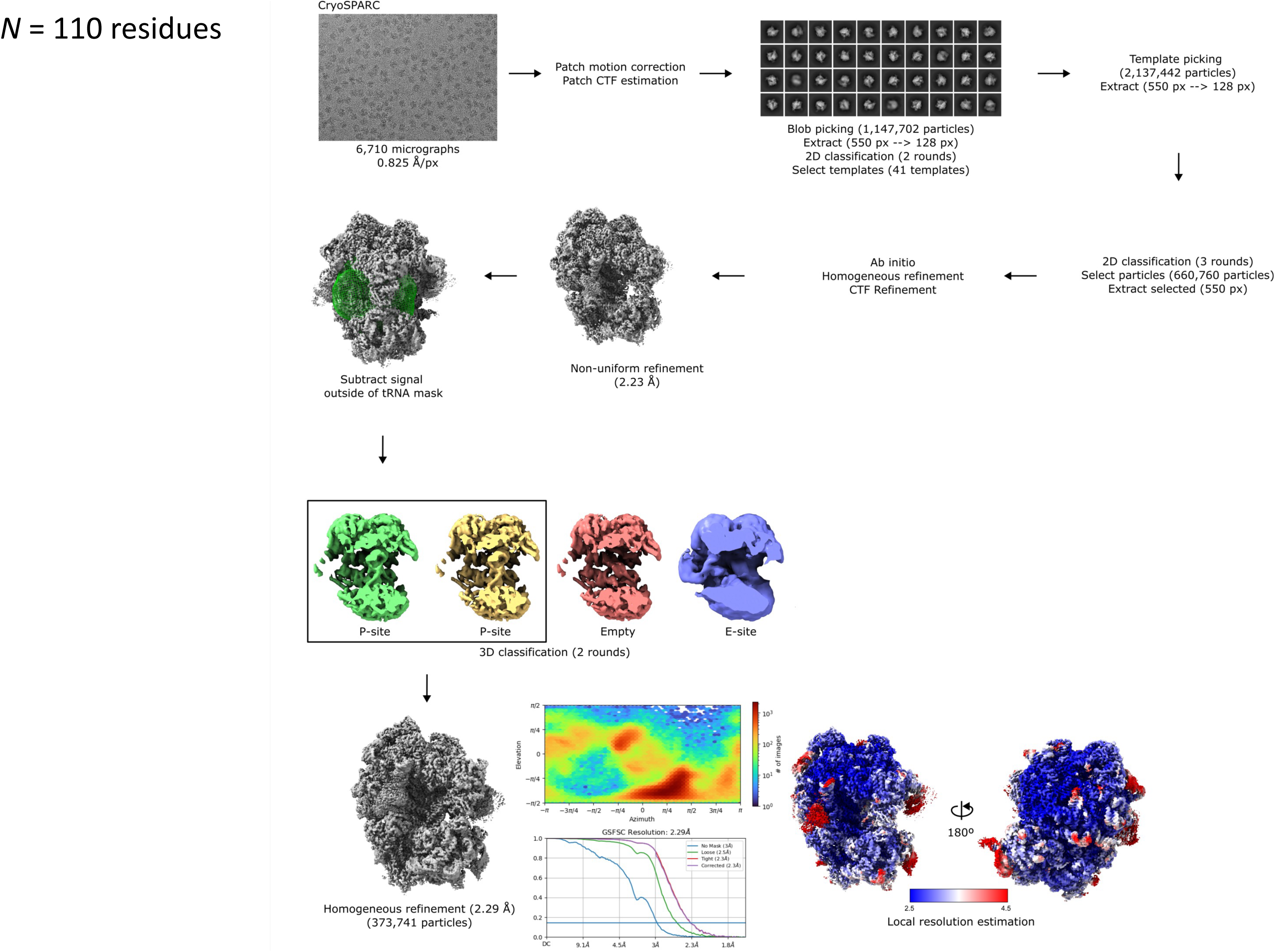
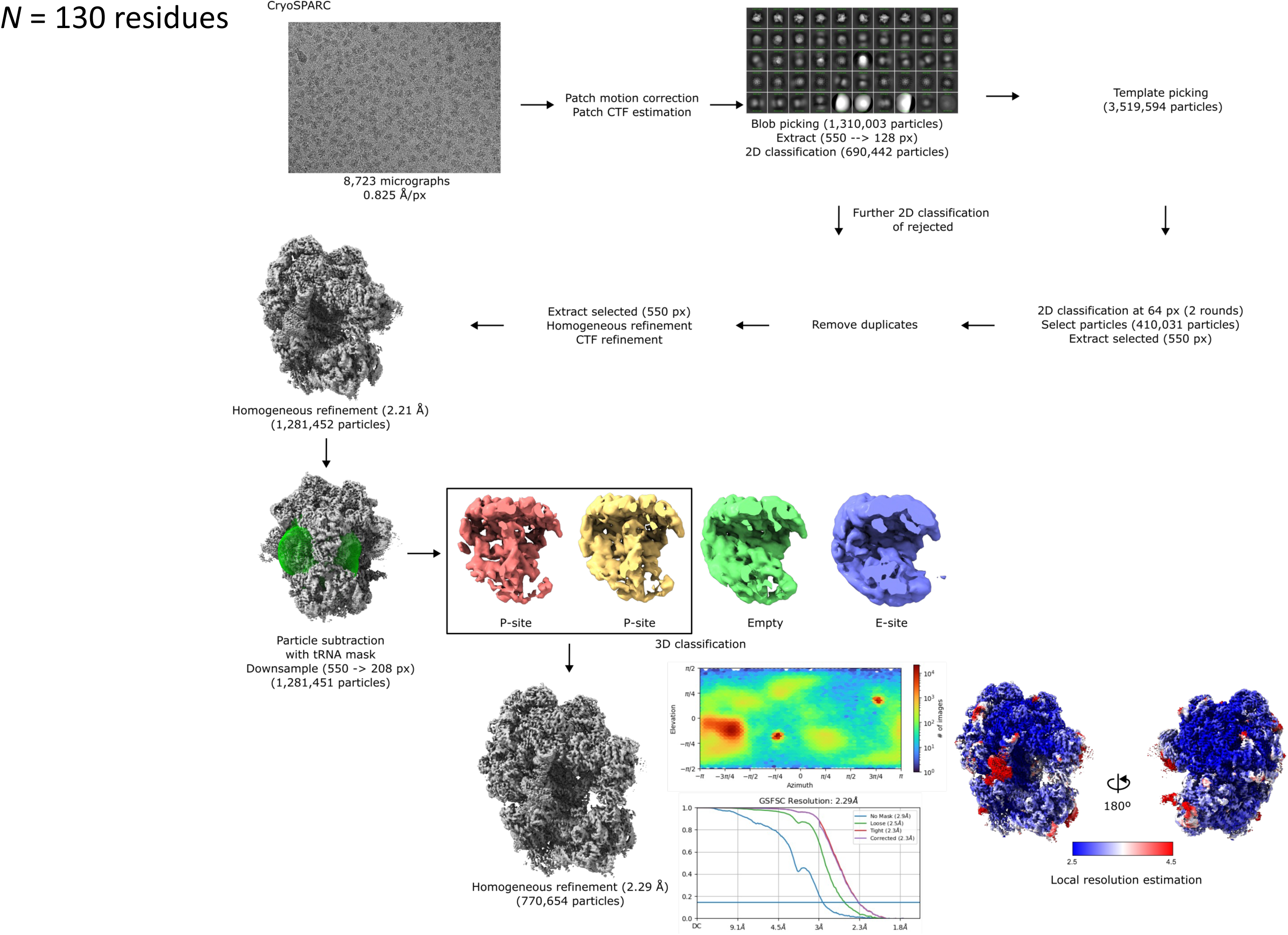
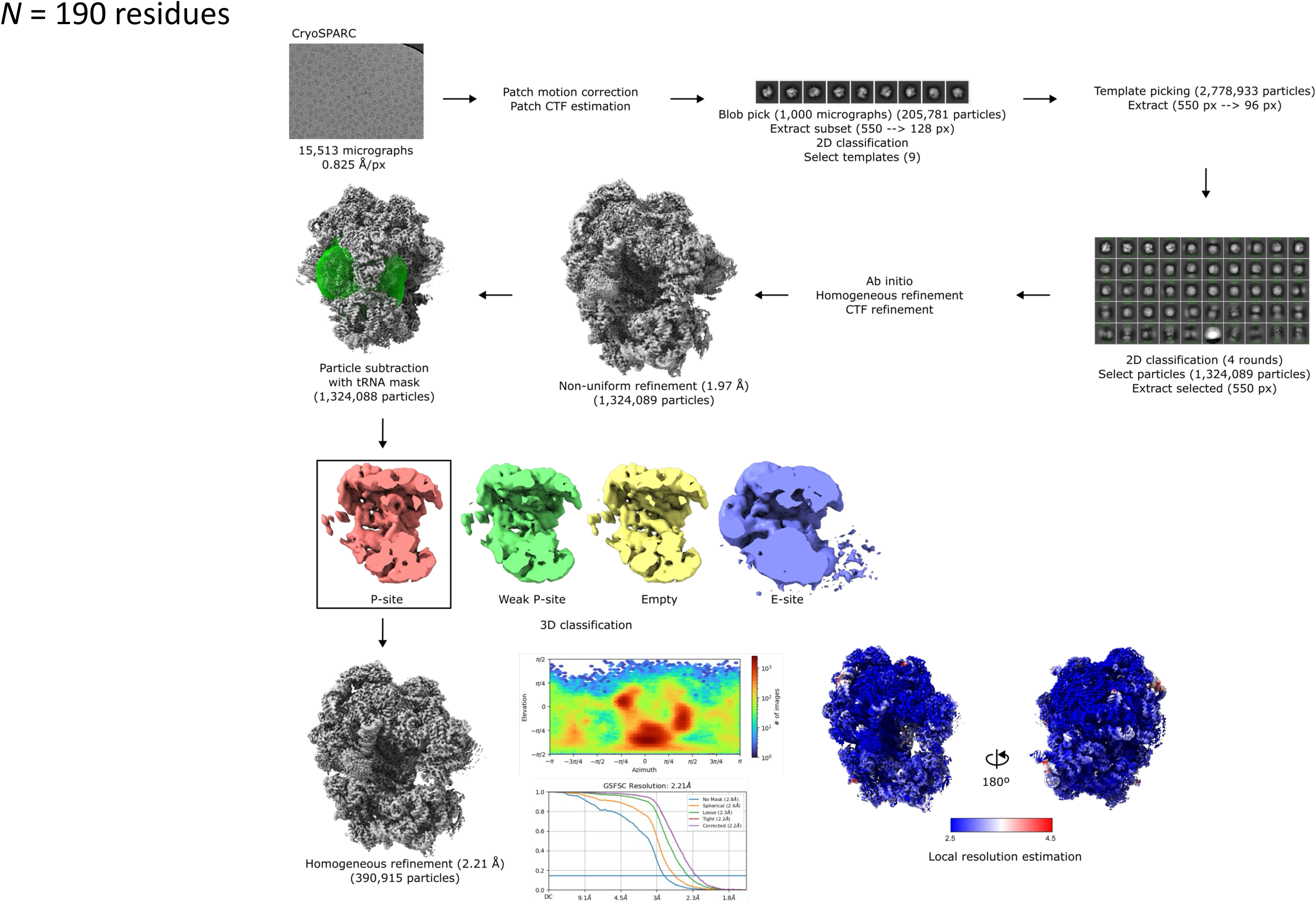
Cryo-EM processing pipelines. (a) *N* = 110 residues RNCs. (b) *N* = 130 residues RNCs. (c) *N* = 190 residues RNCs.

## Legend for Supplemental Movies M1-M3

Full trajectories for the MD simulations carried out for FLuc fragments Δ4-80 (Movie M1), Δ4-110 (Movie M2), and Δ4-150 (Movie M3) have been deposited in Zenodo^78^. Each movie contains three concatenated 1 μs trajectories, all with the same starting conformation as given in Supplemental Fig. S2.

