## Supplemental Table S1 for "Quasi-continuous cotranslational compaction and folding of a multidomain protein"

### **Supplemental Table S1.** Sequences of constructs

**Color code:** FLuc Linker sequence SecM(*Ec*) AP C-tail

FLuc SecM(*Ec*) *N* = 619

MEDAKNIKKGPAPFYPLEDGTAGEQLHKAMKRYALVPGTIAFTDAHIEVDITYAEYFEMSVRLAEAMKRYGLNTNHRIVVC
SENSLQFFMPVLGALFIGVAVAPANDIYNERELLNSMGISQPTVVFVSKKGLQKILNVQKKLPIIQKIIIMDSKTDYQGFQS
MYTFVTSHLPPGFNEYDFVPESFDRDKTIALIMNSSGSTGLPKGVALPHRTACVRFSHARDPIFGNQIIPDTAILSVVPFHH
GFGMFTTLGYLICGFRVVLMYRFEEELFLRSLQDYKIQSALLVPTLFSFFAKSTLIDKYDLSNLHEIASGGAPLSKEVGEAVA
KRFHLPGIRQGYGLTETTSAILITPEGDDKPGAVGKVVPFFEAKVVDLDTGKTLGVNQRGELCVRGPMIMSGYVNNPEAT
NALIDKDGWLHSGDIAYWDEDEHFFIVDRLKSLIKYKGYQVAPAELESILLQHPNIFDAGVAGLPDDDAGELPAAVVVLE
HGKTMTEKEIVDYVASQVTTAKKLRGGVVFVDEVPKGLTGKLDARKIREILIKAKKGGKIAVSGSGSGSGSGSGSGSGS
GSGSGGSGSGSGSGSGSGSGSGSGSGSGSGSGSFSTPVWISQAQGIRAGPDYIKRAVGLPGDKVTYDPVSKELTIQPG
CSSGQACENALPVTYSNVEPSDFVGSSDKQEGEWPTGLRLSRIGGIH

FLuc SecM(*Ec*) *N* = 605

MEDAKNIKKGPAPFYPLEDGTAGEQLHKAMKRYALVPGTIAFTDAHIEVDITYAEYFEMSVRLAEAMKRYGLNTNHRIVVC
SENSLQFFMPVLGALFIGVAVAPANDIYNERELLNSMGISQPTVVFVSKKGLQKILNVQKKLPIIQKIIIMDSKTDYQGFQS
MYTFVTSHLPPGFNEYDFVPESFDRDKTIALIMNSSGSTGLPKGVALPHRTACVRFSHARDPIFGNQIIPDTAILSVVPFHH
GFGMFTTLGYLICGFRVVLMYRFEEELFLRSLQDYKIQSALLVPTLFSFFAKSTLIDKYDLSNLHEIASGGAPLSKEVGEAVA
KRFHLPGIRQGYGLTETTSAILITPEGDDKPGAVGKVVPFFEAKVVDLDTGKTLGVNQRGELCVRGPMIMSGYVNNPEAT
NALIDKDGWLHSGDIAYWDEDEHFFIVDRLKSLIKYKGYQVAPAELESILLQHPNIFDAGVAGLPDDDAGELPAAVVVLE
HGKTMTEKEIVDYVASQVTTAKKLRGGVVFVDEVPKGLTGKLDARKIREILIKAKKGGKIAVSGSGSGSGSGSGSGSGS
GSGSGGSGSGSGSGSGSGSFSTPVWISQAQGIRAGPDYIKRAVGLPGDKVTYDPVSKELTIQPGCSSGQACENALPVT
YSNVEPSDFVGSSDKQEGEWPTGLRLSRIGGIH

FLuc SecM(*Ec*) *N* = 600

MEDAKNIKKGPAPFYPLEDGTAGEQLHKAMKRYALVPGTIAFTDAHIEVDITYAEYFEMSVRLAEAMKRYGLNTNHRIVVC
SENSLQFFMPVLGALFIGVAVAPANDIYNERELLNSMGISQPTVVFVSKKGLQKILNVQKKLPIIQKIIIMDSKTDYQGFQS
MYTFVTSHLPPGFNEYDFVPESFDRDKTIALIMNSSGSTGLPKGVALPHRTACVRFSHARDPIFGNQIIPDTAILSVVPFHH
GFGMFTTLGYLICGFRVVLMYRFEEELFLRSLQDYKIQSALLVPTLFSFFAKSTLIDKYDLSNLHEIASGGAPLSKEVGEAVA
KRFHLPGIRQGYGLTETTSAILITPEGDDKPGAVGKVVPFFEAKVVDLDTGKTLGVNQRGELCVRGPMIMSGYVNNPEAT
NALIDKDGWLHSGDIAYWDEDEHFFIVDRLKSLIKYKGYQVAPAELESILLQHPNIFDAGVAGLPDDDAGELPAAVVVLE
HGKTMTEKEIVDYVASQVTTAKKLRGGVVFVDEVPKGLTGKLDARKIREILIKAKKGGKIAVSGSGSGSGSGSGSGSGS
GSGSGGSGSGSGSGFSTPVWISQAQGIRAGPDYIKRAVGLPGDKVTYDPVSKELTIQPGCSSGQACENALPVTYSNVEP
SDFVGSSDKQEGEWPTGLRLSRIGGIH

FLuc SecM(*Ec*) *N* = 592

MEDAKNIKKGPAPFYPLEDGTAGEQLHKAMKRYALVPGTIAFTDAHIEVDITYAEYFEMSVRLAEAMKRYGLNTNHRIVVC
SENSLQFFMPVLGALFIGVAVAPANDIYNERELLNSMGISQPTVVFVSKKGLQKILNVQKKLPIIQKIIIMDSKTDYQGFQS
MYTFVTSHLPPGFNEYDFVPESFDRDKTIALIMNSSGSTGLPKGVALPHRTACVRFSHARDPIFGNQIIPDTAILSVVPFHH
GFGMFTTLGYLICGFRVVLMYRFEEELFLRSLQDYKIQSALLVPTLFSFFAKSTLIDKYDLSNLHEIASGGAPLSKEVGEAVA
KRFHLPGIRQGYGLTETTSAILITPEGDDKPGAVGKVVPFFEAKVVDLDTGKTLGVNQRGELCVRGPMIMSGYVNNPEAT
NALIDKDGWLHSGDIAYWDEDEHFFIVDRLKSLIKYKGYQVAPAELESILLQHPNIFDAGVAGLPDDDAGELPAAVVVLE
HGKTMTEKEIVDYVASQVTTAKKLRGGVVFVDEVPKGLTGKLDARKIREILIKAKKGGKIAVSGSGSGSGSGSGSGSGS
GSGSGGFSTPVWISQAQGIRAGPDYIKRAVGLPGDKVTYDPVSKELTIQPGCSSGQACENALPVTYSNVEPSDFVGSSD
KQEGEWPTGLRLSRIGGIH

FLuc SecM(*Ec*) *N* = 587

MEDAKNIKKGPAPFYPLEDGTAGEQLHKAMKRYALVPGTIAFTDAHIEVDITYAEYFEMSVRLAEAMKRYGLNTNHRIVVC
SENSLQFFMPVLGALFIGVAVAPANDIYNERELLNSMGISQPTVVFVSKKGLQKILNVQKKLPIIQKIIIMDSKTDYQGFQS
MYTFVTSHLPPGFNEYDFVPESFDRDKTIALIMNSSGSTGLPKGVALPHRTACVRFSHARDPIFGNQIIPDTAILSVVPFHH
GFGMFTTLGYLICGFRVVLMYRFEEELFLRSLQDYKIQSALLVPTLFSFFAKSTLIDKYDLSNLHEIASGGAPLSKEVGEAVA
KRFHLPGIRQGYGLTETTSAILITPEGDDKPGAVGKVVPFFEAKVVDLDTGKTLGVNQRGELCVRGPMIMSGYVNNPEAT
NALIDKDGWLHSGDIAYWDEDEHFFIVDRLKSLIKYKGYQVAPAELESILLQHPNIFDAGVAGLPDDDAGELPAAVVVLE
HGKTMTEKEIVDYVASQVTTAKKLRGGVVFVDEVPKGLTGKLDARKIREILIKAKKGGKIAVSGSGSGSGSGSGSGSGSGS
GFSTPVWISQAQGIRAGPDYIKRAVGLPGDKVTYDPVSKELTIQPGCSSGQACENALPVTYSNVEPSDFVGSSDKQEGE
WPTGLRLSRIGGIH

FLuc SecM(*Ec*) *N* = 585

MEDAKNIKKGPAPFYPLEDGTAGEQLHKAMKRYALVPGTIAFTDAHIEVDITYAEYFEMSVRLAEAMKRYGLNTNHRIVVC
SENSLQFFMPVLGALFIGVAVAPANDIYNERELLNSMGISQPTVVFVSKKGLQKILNVQKKLPIIQKIIIMDSKTDYQGFQS
MYTFVTSHLPPGFNEYDFVPESFDRDKTIALIMNSSGSTGLPKGVALPHRTACVRFSHARDPIFGNQIIPDTAILSVVPFHH
GFGMFTTLGYLICGFRVVLMYRFEEELFLRSLQDYKIQSALLVPTLFSFFAKSTLIDKYDLSNLHEIASGGAPLSKEVGEAVA
KRFHLPGIRQGYGLTETTSAILITPEGDDKPGAVGKVVPFFEAKVVDLDTGKTLGVNQRGELCVRGPMIMSGYVNNPEAT
NALIDKDGWLHSGDIAYWDEDEHFFIVDRLKSLIKYKGYQVAPAELESILLQHPNIFDAGVAGLPDDDAGELPAAVVVLE
HGKTMTEKEIVDYVASQVTTAKKLRGGVVFVDEVPKGLTGKLDARKIREILIKAKKGGKIAVSGSGSGSGSGSGSGSGSGF
STPVWISQAQGIRAGPDYIKRAVGLPGDKVTYDPVSKELTIQPGCSSGQACENALPVTYSNVEPSDFVGSSDKQEGEWP
TGLRLSRIGGIH

FLuc SecM(*Ec*) *N* = 583

MEDAKNIKKGPAPFYPLEDGTAGEQLHKAMKRYALVPGTIAFTDAHIEVDITYAEYFEMSVRLAEAMKRYGLNTNHRIVVC
SENSLQFFMPVLGALFIGVAVAPANDIYNERELLNSMGISQPTVVFVSKKGLQKILNVQKKLPIIQKIIIMDSKTDYQGFQS
MYTFVTSHLPPGFNEYDFVPESFDRDKTIALIMNSSGSTGLPKGVALPHRTACVRFSHARDPIFGNQIIPDTAILSVVPFHH
GFGMFTTLGYLICGFRVVLMYRFEEELFLRSLQDYKIQSALLVPTLFSFFAKSTLIDKYDLSNLHEIASGGAPLSKEVGEAVA
KRFHLPGIRQGYGLTETTSAILITPEGDDKPGAVGKVVPFFEAKVVDLDTGKTLGVNQRGELCVRGPMIMSGYVNNPEAT
NALIDKDGWLHSGDIAYWDEDEHFFIVDRLKSLIKYKGYQVAPAELESILLQHPNIFDAGVAGLPDDDAGELPAAVVVLE
HGKTMTEKEIVDYVASQVTTAKKLRGGVVFVDEVPKGLTGKLDARKIREILIKAKKGGKIAVSGSGSGSGSGSGSGSGFST
PVWISQAQGIRAGPDYIKRAVGLPGDKVTYDPVSKELTIQPGCSSGQACENALPVTYSNVEPSDFVGSSDKQEGEWPTG
LRLSRIGGIH

FLuc SecM(*Ec*) *N* = 580

MEDAKNIKKGPAPFYPLEDGTAGEQLHKAMKRYALVPGTIAFTDAHIEVDITYAEYFEMSVRLAEAMKRYGLNTNHRIVVC
SENSLQFFMPVLGALFIGVAVAPANDIYNERELLNSMGISQPTVVFVSKKGLQKILNVQKKLPIIQKIIIMDSKTDYQGFQS
MYTFVTSHLPPGFNEYDFVPESFDRDKTIALIMNSSGSTGLPKGVALPHRTACVRFSHARDPIFGNQIIPDTAILSVVPFHH
GFGMFTTLGYLICGFRVVLMYRFEEELFLRSLQDYKIQSALLVPTLFSFFAKSTLIDKYDLSNLHEIASGGAPLSKEVGEAVA
KRFHLPGIRQGYGLTETTSAILITPEGDDKPGAVGKVVPFFEAKVVDLDTGKTLGVNQRGELCVRGPMIMSGYVNNPEAT
NALIDKDGWLHSGDIAYWDEDEHFFIVDRLKSLIKYKGYQVAPAELESILLQHPNIFDAGVAGLPDDDAGELPAAVVVLE
HGKTMTEKEIVDYVASQVTTAKKLRGGVVFVDEVPKGLTGKLDARKIREILIKAKKGGKIAVSGSGSGSGSGSFSTPVWI
SQAQGIRAGPDYIKRAVGLPGDKVTYDPVSKELTIQPGCSSGQACENALPVTYSNVEPSDFVGSSDKQEGEWPTGLRLS
RIGGIH

FLuc SecM(*Ec*) *N* = 575

MEDAKNIKKGPAPFYPLEDGTAGEQLHKAMKRYALVPGTIAFTDAHIEVDITYAEYFEMSVRLAEAMKRYGLNTNHRIVVC
SENSLQFFMPVLGALFIGVAVAPANDIYNERELLNSMGISQPTVVFVSKKGLQKILNVQKKLPIIQKIIIMDSKTDYQGFQS
MYTFVTSHLPPGFNEYDFVPESFDRDKTIALIMNSSGSTGLPKGVALPHRTACVRFSHARDPIFGNQIIPDTAILSVVPFHH
GFGMFTTLGYLICGFRVVLMYRFEEELFLRSLQDYKIQSALLVPTLFSFFAKSTLIDKYDLSNLHEIASGGAPLSKEVGEAVA
KRFHLPGIRQGYGLTETTSAILITPEGDDKPGAVGKVVPFFEAKVVDLDTGKTLGVNQRGELCVRGPMIMSGYVNNPEAT
NALIDKDGWLHSGDIAYWDEDEHFFIVDRLKSLIKYKGYQVAPAELESILLQHPNIFDAGVAGLPDDDAGELPAAVVVLE

HGKTMTEKEIVDYVASQVTTAKKLRGGVVFVDEVPKGLTGKLDARKIREILIKAKKGGKIAVSGSGSGSGFSTPVWISQAQ
GIRAGPDYIKRAVGLPGDKVTYDPVSKELTIQPGCSSGQACENALPVTYSNVEPSDFVGSSDKQEGEWPTGLRLSRIGGI
H

FLuc SecM(*Ec*) $N$ = 570

MEDAKNIKKGPAPFYPLEDGTAGEQLHKAMKRYALVPGTIAFTDAHIEVDITYAEYFEMSVRLAEAMKRYGLNTNHRIVVC
SENSLQFFMPVLGALFIGVAVAPANDIYNERELLNSMGISQPTVVFVSKKGLQKILNVQKKLPIIQKIIIMDSKTDYQGFQS
MYTFVTSHLPPGFNEYDFVPESFDRDKTIALIMNSSGSTGLPKGVALPHRTACVRFSHARDPIFGNQIIPDTAILSVVPFHH
GFGMFTTLGYLICGFRVVLMYRFEEELFLRSLQDYKIQSALLVPTLFSFFAKSTLIDKYDLSNLHEIASGGAPLSKEVGEAVA
KRFHLPGIRQGYGLTETTSAILITPEGDDKPGAVGKVVPFFEAKVVDLDTGKTLGVNQRGELCVRGPMIMSGYVNNPEAT
NALIDKDGWLHSGDIAYWDEDEHFFIVDRLKSLIKYKGYQVAPAELESILLQHPNIFDAGVAGLPDDDAGELPAAVVVLE
HGKTMTEKEIVDYVASQVTTAKKLRGGVVFVDEVPKGLTGKLDARKIREILIKAKKGGKIAVSGSFSTPVWISQAQGIRAGP
DYIKRAVGLPGDKVTYDPVSKELTIQPGCSSGQACENALPVTYSNVEPSDFVGSSDKQEGEWPTGLRLSRIGGIH

FLuc SecM(*Ec*) $N$ = 565

MEDAKNIKKGPAPFYPLEDGTAGEQLHKAMKRYALVPGTIAFTDAHIEVDITYAEYFEMSVRLAEAMKRYGLNTNHRIVVC
SENSLQFFMPVLGALFIGVAVAPANDIYNERELLNSMGISQPTVVFVSKKGLQKILNVQKKLPIIQKIIIMDSKTDYQGFQS
MYTFVTSHLPPGFNEYDFVPESFDRDKTIALIMNSSGSTGLPKGVALPHRTACVRFSHARDPIFGNQIIPDTAILSVVPFHH
GFGMFTTLGYLICGFRVVLMYRFEEELFLRSLQDYKIQSALLVPTLFSFFAKSTLIDKYDLSNLHEIASGGAPLSKEVGEAVA
KRFHLPGIRQGYGLTETTSAILITPEGDDKPGAVGKVVPFFEAKVVDLDTGKTLGVNQRGELCVRGPMIMSGYVNNPEAT
NALIDKDGWLHSGDIAYWDEDEHFFIVDRLKSLIKYKGYQVAPAELESILLQHPNIFDAGVAGLPDDDAGELPAAVVVLE
HGKTMTEKEIVDYVASQVTTAKKLRGGVVFVDEVPKGLTGKLDARKIREILIKAKKGSGSFSTPVWISQAQGIRAGPDYIKR
AVGLPGDKVTYDPVSKELTIQPGCSSGQACENALPVTYSNVEPSDFVGSSDKQEGEWPTGLRLSRIGGIH

FLuc SecM(*Ec*) $N$ = 562

MEDAKNIKKGPAPFYPLEDGTAGEQLHKAMKRYALVPGTIAFTDAHIEVDITYAEYFEMSVRLAEAMKRYGLNTNHRIVVC
SENSLQFFMPVLGALFIGVAVAPANDIYNERELLNSMGISQPTVVFVSKKGLQKILNVQKKLPIIQKIIIMDSKTDYQGFQS
MYTFVTSHLPPGFNEYDFVPESFDRDKTIALIMNSSGSTGLPKGVALPHRTACVRFSHARDPIFGNQIIPDTAILSVVPFHH
GFGMFTTLGYLICGFRVVLMYRFEEELFLRSLQDYKIQSALLVPTLFSFFAKSTLIDKYDLSNLHEIASGGAPLSKEVGEAVA
KRFHLPGIRQGYGLTETTSAILITPEGDDKPGAVGKVVPFFEAKVVDLDTGKTLGVNQRGELCVRGPMIMSGYVNNPEAT
NALIDKDGWLHSGDIAYWDEDEHFFIVDRLKSLIKYKGYQVAPAELESILLQHPNIFDAGVAGLPDDDAGELPAAVVVLE
HGKTMTEKEIVDYVASQVTTAKKLRGGVVFVDEVPKGLTGKLDARKIREILIKASGSFSTPVWISQAQGIRAGPDYIKRAVG
LPGDKVTYDPVSKELTIQPGCSSGQACENALPVTYSNVEPSDFVGSSDKQEGEWPTGLRLSRIGGIH

FLuc SecM(*Ec*) $N$ = 560

MEDAKNIKKGPAPFYPLEDGTAGEQLHKAMKRYALVPGTIAFTDAHIEVDITYAEYFEMSVRLAEAMKRYGLNTNHRIVVC
SENSLQFFMPVLGALFIGVAVAPANDIYNERELLNSMGISQPTVVFVSKKGLQKILNVQKKLPIIQKIIIMDSKTDYQGFQS
MYTFVTSHLPPGFNEYDFVPESFDRDKTIALIMNSSGSTGLPKGVALPHRTACVRFSHARDPIFGNQIIPDTAILSVVPFHH
GFGMFTTLGYLICGFRVVLMYRFEEELFLRSLQDYKIQSALLVPTLFSFFAKSTLIDKYDLSNLHEIASGGAPLSKEVGEAVA
KRFHLPGIRQGYGLTETTSAILITPEGDDKPGAVGKVVPFFEAKVVDLDTGKTLGVNQRGELCVRGPMIMSGYVNNPEAT
NALIDKDGWLHSGDIAYWDEDEHFFIVDRLKSLIKYKGYQVAPAELESILLQHPNIFDAGVAGLPDDDAGELPAAVVVLE
HGKTMTEKEIVDYVASQVTTAKKLRGGVVFVDEVPKGLTGKLDARKIREILISGSFSTPVWISQAQGIRAGPDYIKRAVGLP
GDKVTYDPVSKELTIQPGCSSGQACENALPVTYSNVEPSDFVGSSDKQEGEWPTGLRLSRIGGIH

FLuc SecM(*Ec*) $N$ = 558

MEDAKNIKKGPAPFYPLEDGTAGEQLHKAMKRYALVPGTIAFTDAHIEVDITYAEYFEMSVRLAEAMKRYGLNTNHRIVVC
SENSLQFFMPVLGALFIGVAVAPANDIYNERELLNSMGISQPTVVFVSKKGLQKILNVQKKLPIIQKIIIMDSKTDYQGFQS
MYTFVTSHLPPGFNEYDFVPESFDRDKTIALIMNSSGSTGLPKGVALPHRTACVRFSHARDPIFGNQIIPDTAILSVVPFHH
GFGMFTTLGYLICGFRVVLMYRFEEELFLRSLQDYKIQSALLVPTLFSFFAKSTLIDKYDLSNLHEIASGGAPLSKEVGEAVA
KRFHLPGIRQGYGLTETTSAILITPEGDDKPGAVGKVVPFFEAKVVDLDTGKTLGVNQRGELCVRGPMIMSGYVNNPEAT
NALIDKDGWLHSGDIAYWDEDEHFFIVDRLKSLIKYKGYQVAPAELESILLQHPNIFDAGVAGLPDDDAGELPAAVVVLE

HGKTMTEKEIVDYVASQVTTAKKLRGGVVFVDEVPKGLTGKLDARKIREISGSFSTPVWISQAQGIRAGPDYIKRAVGLPG
DKVTYDPVSKELTIQPGCSSGQACENALPVTYSNVEPSDFVGSSDKQEGEWPTGLRLSRIGGIH

FLuc SecM(*Ec*) *N* = 555

MEDAKNIKKGPAPFYPLEDGTAGEQLHKAMKRYALVPGTIAFTDAHIEVDITYAEYFEMSVRLAEAMKRYGLNTNHRIVVC
SENSLQFFMPVLGALFIGVAVAPANDIYNERELLNSMGISQPTVVFVSKKGLQKILNVQKKLPIIQKIIIMDSKTDYQGFQS
MYTFVTSHLPPGFNEYDFVPESFDRDKTIALIMNSSGSTGLPKGVALPHRTACVRFSHARDPIFGNQIIPDTAILSVVPFHH
GFGMFTTLGYLICGFRVVLMYRFEEELFLRSLQDYKIQSALLVPTLFSFFAKSTLIDKYDLSNLHEIASGGAPLSKEVGEAVA
KRFHLPGIRQGYGLTETTSAILITPEGDDKPGAVGKVVPFFEAKVVDLDTGKTLGVNQRGELCVRGPMIMSGYVNNPEAT
NALIDKDGWLHSGDIAYWDEDEHFFIVDRLKSLIKYKGYQVAPAELESILLQHPNIFDAGVAGLPDDDAGELPAAVVVLE
HGKTMTEKEIVDYVASQVTTAKKLRGGVVFVDEVPKGLTGKLDARKISGSFSTPVWISQAQGIRAGPDYIKRAVGLPGDKV
TYDPVSKELTIQPGCSSGQACENALPVTYSNVEPSDFVGSSDKQEGEWPTGLRLSRIGGIH

FLuc SecM(*Ec*) *N* = 550

MEDAKNIKKGPAPFYPLEDGTAGEQLHKAMKRYALVPGTIAFTDAHIEVDITYAEYFEMSVRLAEAMKRYGLNTNHRIVVC
SENSLQFFMPVLGALFIGVAVAPANDIYNERELLNSMGISQPTVVFVSKKGLQKILNVQKKLPIIQKIIIMDSKTDYQGFQS
MYTFVTSHLPPGFNEYDFVPESFDRDKTIALIMNSSGSTGLPKGVALPHRTACVRFSHARDPIFGNQIIPDTAILSVVPFHH
GFGMFTTLGYLICGFRVVLMYRFEEELFLRSLQDYKIQSALLVPTLFSFFAKSTLIDKYDLSNLHEIASGGAPLSKEVGEAVA
KRFHLPGIRQGYGLTETTSAILITPEGDDKPGAVGKVVPFFEAKVVDLDTGKTLGVNQRGELCVRGPMIMSGYVNNPEAT
NALIDKDGWLHSGDIAYWDEDEHFFIVDRLKSLIKYKGYQVAPAELESILLQHPNIFDAGVAGLPDDDAGELPAAVVVLE
HGKTMTEKEIVDYVASQVTTAKKLRGGVVFVDEVPKGLTGKLSGSFSTPVWISQAQGIRAGPDYIKRAVGLPGDKVTYDP
VSKELTIQPGCSSGQACENALPVTYSNVEPSDFVGSSDKQEGEWPTGLRLSRIGGIH

FLuc SecM(*Ec*) *N* = 545

MEDAKNIKKGPAPFYPLEDGTAGEQLHKAMKRYALVPGTIAFTDAHIEVDITYAEYFEMSVRLAEAMKRYGLNTNHRIVVC
SENSLQFFMPVLGALFIGVAVAPANDIYNERELLNSMGISQPTVVFVSKKGLQKILNVQKKLPIIQKIIIMDSKTDYQGFQS
MYTFVTSHLPPGFNEYDFVPESFDRDKTIALIMNSSGSTGLPKGVALPHRTACVRFSHARDPIFGNQIIPDTAILSVVPFHH
GFGMFTTLGYLICGFRVVLMYRFEEELFLRSLQDYKIQSALLVPTLFSFFAKSTLIDKYDLSNLHEIASGGAPLSKEVGEAVA
KRFHLPGIRQGYGLTETTSAILITPEGDDKPGAVGKVVPFFEAKVVDLDTGKTLGVNQRGELCVRGPMIMSGYVNNPEAT
NALIDKDGWLHSGDIAYWDEDEHFFIVDRLKSLIKYKGYQVAPAELESILLQHPNIFDAGVAGLPDDDAGELPAAVVVLE
HGKTMTEKEIVDYVASQVTTAKKLRGGVVFVDEVPKGSGSFSTPVWISQAQGIRAGPDYIKRAVGLPGDKVTYDPVSKELT
IQPGCSSGQACENALPVTYSNVEPSDFVGSSDKQEGEWPTGLRLSRIGGIH

FLuc SecM(*Ec*) *N* = 540

MEDAKNIKKGPAPFYPLEDGTAGEQLHKAMKRYALVPGTIAFTDAHIEVDITYAEYFEMSVRLAEAMKRYGLNTNHRIVVC
SENSLQFFMPVLGALFIGVAVAPANDIYNERELLNSMGISQPTVVFVSKKGLQKILNVQKKLPIIQKIIIMDSKTDYQGFQS
MYTFVTSHLPPGFNEYDFVPESFDRDKTIALIMNSSGSTGLPKGVALPHRTACVRFSHARDPIFGNQIIPDTAILSVVPFHH
GFGMFTTLGYLICGFRVVLMYRFEEELFLRSLQDYKIQSALLVPTLFSFFAKSTLIDKYDLSNLHEIASGGAPLSKEVGEAVA
KRFHLPGIRQGYGLTETTSAILITPEGDDKPGAVGKVVPFFEAKVVDLDTGKTLGVNQRGELCVRGPMIMSGYVNNPEAT
NALIDKDGWLHSGDIAYWDEDEHFFIVDRLKSLIKYKGYQVAPAELESILLQHPNIFDAGVAGLPDDDAGELPAAVVVLE
HGKTMTEKEIVDYVASQVTTAKKLRGGVVFVDSGSFSTPVWISQAQGIRAGPDYIKRAVGLPGDKVTYDPVSKELTIQPGC
SSGQACENALPVTYSNVEPSDFVGSSDKQEGEWPTGLRLSRIGGIH

FLuc SecM(*Ec*) *N* = 535

MEDAKNIKKGPAPFYPLEDGTAGEQLHKAMKRYALVPGTIAFTDAHIEVDITYAEYFEMSVRLAEAMKRYGLNTNHRIVVC
SENSLQFFMPVLGALFIGVAVAPANDIYNERELLNSMGISQPTVVFVSKKGLQKILNVQKKLPIIQKIIIMDSKTDYQGFQS
MYTFVTSHLPPGFNEYDFVPESFDRDKTIALIMNSSGSTGLPKGVALPHRTACVRFSHARDPIFGNQIIPDTAILSVVPFHH
GFGMFTTLGYLICGFRVVLMYRFEEELFLRSLQDYKIQSALLVPTLFSFFAKSTLIDKYDLSNLHEIASGGAPLSKEVGEAVA
KRFHLPGIRQGYGLTETTSAILITPEGDDKPGAVGKVVPFFEAKVVDLDTGKTLGVNQRGELCVRGPMIMSGYVNNPEAT
NALIDKDGWLHSGDIAYWDEDEHFFIVDRLKSLIKYKGYQVAPAELESILLQHPNIFDAGVAGLPDDDAGELPAAVVVLE

HGKTMTEKEIVDYVASQVTTAKKLRGGSGSFSTPVWISQAQGIRAGPDYIKRAVGLPGDKVTYDPVSKELTIQPGCSSGQ
ACENALPVTYSNVEPSDFVGSSDKQEGEWPTGLRLSRIGGIH

FLuc SecM(*Ec*) *N* = 530

MEDAKNIKKGPAPFYPLEDGTAGEQLHKAMKRYALVPGTIAFTDAHIEVDITYAEYFEMSVRLAEAMKRYGLNTNHRIVVC
SENSLQFFMPVLGALFIGVAVAPANDIYNERELLNSMGISQPTVVFVSKKGLQKILNVQKKLPIIQKIIIMDSKTDYQGFQS
MYTFVTSHLPPGFNEYDFVPESFDRDKTIALIMNSSGSTGLPKGVALPHRTACVRFSHARDPIFGNQIIPDTAILSVVPFHH
GFGMFTTLGYLICGFRVVLMYRFEEELFLRSLQDYKIQSALLVPTLFSFFAKSTLIDKYDLSNLHEIASGGAPLSKEVGEAVA
KRFHLPGIRQGYGLTETTSAILITPEGDDKPGAVGKVVPFFEAKVVDLDTGKTLGVNQRGELCVRGPMIMSGYVNNPEAT
NALIDKDGWLHSGDIAYWDEDEHFFIVDRLKSLIKYKGYQVAPAELESILLQHPNIFDAGVAGLPDDDAGELPAAVVVLE
HGKTMTEKEIVDYVASQVTTAKSGSFSTPVWISQAQGIRAGPDYIKRAVGLPGDKVTYDPVSKELTIQPGCSSGQACENA
LPVTYSNVEPSDFVGSSDKQEGEWPTGLRLSRIGGIH

FLuc SecM(*Ec*) *N* = 525

MEDAKNIKKGPAPFYPLEDGTAGEQLHKAMKRYALVPGTIAFTDAHIEVDITYAEYFEMSVRLAEAMKRYGLNTNHRIVVC
SENSLQFFMPVLGALFIGVAVAPANDIYNERELLNSMGISQPTVVFVSKKGLQKILNVQKKLPIIQKIIIMDSKTDYQGFQS
MYTFVTSHLPPGFNEYDFVPESFDRDKTIALIMNSSGSTGLPKGVALPHRTACVRFSHARDPIFGNQIIPDTAILSVVPFHH
GFGMFTTLGYLICGFRVVLMYRFEEELFLRSLQDYKIQSALLVPTLFSFFAKSTLIDKYDLSNLHEIASGGAPLSKEVGEAVA
KRFHLPGIRQGYGLTETTSAILITPEGDDKPGAVGKVVPFFEAKVVDLDTGKTLGVNQRGELCVRGPMIMSGYVNNPEAT
NALIDKDGWLHSGDIAYWDEDEHFFIVDRLKSLIKYKGYQVAPAELESILLQHPNIFDAGVAGLPDDDAGELPAAVVVLE
HGKTMTEKEIVDYVASQSGSFSTPVWISQAQGIRAGPDYIKRAVGLPGDKVTYDPVSKELTIQPGCSSGQACENALPVTY
SNVEPSDFVGSSDKQEGEWPTGLRLSRIGGIH

FLuc SecM(*Ec*) *N* = 520

MEDAKNIKKGPAPFYPLEDGTAGEQLHKAMKRYALVPGTIAFTDAHIEVDITYAEYFEMSVRLAEAMKRYGLNTNHRIVVC
SENSLQFFMPVLGALFIGVAVAPANDIYNERELLNSMGISQPTVVFVSKKGLQKILNVQKKLPIIQKIIIMDSKTDYQGFQS
MYTFVTSHLPPGFNEYDFVPESFDRDKTIALIMNSSGSTGLPKGVALPHRTACVRFSHARDPIFGNQIIPDTAILSVVPFHH
GFGMFTTLGYLICGFRVVLMYRFEEELFLRSLQDYKIQSALLVPTLFSFFAKSTLIDKYDLSNLHEIASGGAPLSKEVGEAVA
KRFHLPGIRQGYGLTETTSAILITPEGDDKPGAVGKVVPFFEAKVVDLDTGKTLGVNQRGELCVRGPMIMSGYVNNPEAT
NALIDKDGWLHSGDIAYWDEDEHFFIVDRLKSLIKYKGYQVAPAELESILLQHPNIFDAGVAGLPDDDAGELPAAVVVLE
HGKTMTEKEIVDSGSFSTPVWISQAQGIRAGPDYIKRAVGLPGDKVTYDPVSKELTIQPGCSSGQACENALPVTYSNVEP
SDFVGSSDKQEGEWPTGLRLSRIGGIH

FLuc SecM(*Ec*) *N* = 515

MEDAKNIKKGPAPFYPLEDGTAGEQLHKAMKRYALVPGTIAFTDAHIEVDITYAEYFEMSVRLAEAMKRYGLNTNHRIVVC
SENSLQFFMPVLGALFIGVAVAPANDIYNERELLNSMGISQPTVVFVSKKGLQKILNVQKKLPIIQKIIIMDSKTDYQGFQS
MYTFVTSHLPPGFNEYDFVPESFDRDKTIALIMNSSGSTGLPKGVALPHRTACVRFSHARDPIFGNQIIPDTAILSVVPFHH
GFGMFTTLGYLICGFRVVLMYRFEEELFLRSLQDYKIQSALLVPTLFSFFAKSTLIDKYDLSNLHEIASGGAPLSKEVGEAVA
KRFHLPGIRQGYGLTETTSAILITPEGDDKPGAVGKVVPFFEAKVVDLDTGKTLGVNQRGELCVRGPMIMSGYVNNPEAT
NALIDKDGWLHSGDIAYWDEDEHFFIVDRLKSLIKYKGYQVAPAELESILLQHPNIFDAGVAGLPDDDAGELPAAVVVLE
HGKTMTESGSFSTPVWISQAQGIRAGPDYIKRAVGLPGDKVTYDPVSKELTIQPGCSSGQACENALPVTYSNVEPSDFV
GSSDKQEGEWPTGLRLSRIGGIH

FLuc SecM(*Ec*) *N* = 510

MEDAKNIKKGPAPFYPLEDGTAGEQLHKAMKRYALVPGTIAFTDAHIEVDITYAEYFEMSVRLAEAMKRYGLNTNHRIVVC
SENSLQFFMPVLGALFIGVAVAPANDIYNERELLNSMGISQPTVVFVSKKGLQKILNVQKKLPIIQKIIIMDSKTDYQGFQS
MYTFVTSHLPPGFNEYDFVPESFDRDKTIALIMNSSGSTGLPKGVALPHRTACVRFSHARDPIFGNQIIPDTAILSVVPFHH
GFGMFTTLGYLICGFRVVLMYRFEEELFLRSLQDYKIQSALLVPTLFSFFAKSTLIDKYDLSNLHEIASGGAPLSKEVGEAVA
KRFHLPGIRQGYGLTETTSAILITPEGDDKPGAVGKVVPFFEAKVVDLDTGKTLGVNQRGELCVRGPMIMSGYVNNPEAT
NALIDKDGWLHSGDIAYWDEDEHFFIVDRLKSLIKYKGYQVAPAELESILLQHPNIFDAGVAGLPDDDAGELPAAVVVLE

HGSGSFSTPVWISQAQGIRAGPDYIKRAVGLPGDKVTYDPVSKELTIQPGCSSGQACENALPVTYSNVEPSDFVGSSDK
QEGEWPTGLRLSRIGGIH

FLuc SecM(*Ec*) *N* = 505

MEDAKNIKKGPAPFYPLEDGTAGEQLHKAMKRYALVPGTIAFTDAHIEVDITYAEYFEMSVRLAEAMKRYGLNTNHRIVVC
SENSLQFFMPVLGALFIGVAVAPANDIYNERELLNSMGISQPTVVFVSKKGLQKILNVQKKLPIIQKIIIMDSKTDYQGFQS
MYTFVTSHLPPGFNEYDFVPESFDRDKTIALIMNSSGSTGLPKGVALPHRTACVRFSHARDPIFGNQIIPDTAILSVVPFHH
GFGMFTTLGYLICGFRVVLMYRFEEELFLRSLQDYKIQSALLVPTLFSFFAKSTLIDKYDLSNLHEIASGGAPLSKEVGEAVA
KRFHLPGIRQGYGLTETTSAILITPEGDDKPGAVGKVVPFFEAKVVDLDTGKTLGVNQRGELCVRGPMIMSGYVNNPEAT
NALIDKDGWLHSGDIAYWDEDEHFFIVDRLKSLIKYKGYQVAPAELESILLQHPNIFDAGVAGLPDDDAGELPAAVVSGS
FSTPVWISQAQGIRAGPDYIKRAVGLPGDKVTYDPVSKELTIQPGCSSGQACENALPVTYSNVEPSDFVGSSDKQEGEW
PTGLRLSRIGGIH

FLuc SecM(*Ec*) *N* = 500

MEDAKNIKKGPAPFYPLEDGTAGEQLHKAMKRYALVPGTIAFTDAHIEVDITYAEYFEMSVRLAEAMKRYGLNTNHRIVVC
SENSLQFFMPVLGALFIGVAVAPANDIYNERELLNSMGISQPTVVFVSKKGLQKILNVQKKLPIIQKIIIMDSKTDYQGFQS
MYTFVTSHLPPGFNEYDFVPESFDRDKTIALIMNSSGSTGLPKGVALPHRTACVRFSHARDPIFGNQIIPDTAILSVVPFHH
GFGMFTTLGYLICGFRVVLMYRFEEELFLRSLQDYKIQSALLVPTLFSFFAKSTLIDKYDLSNLHEIASGGAPLSKEVGEAVA
KRFHLPGIRQGYGLTETTSAILITPEGDDKPGAVGKVVPFFEAKVVDLDTGKTLGVNQRGELCVRGPMIMSGYVNNPEAT
NALIDKDGWLHSGDIAYWDEDEHFFIVDRLKSLIKYKGYQVAPAELESILLQHPNIFDAGVAGLPDDDAGELSGSFSTPV
WISQAQGIRAGPDYIKRAVGLPGDKVTYDPVSKELTIQPGCSSGQACENALPVTYSNVEPSDFVGSSDKQEGEWPTGLR
LSRIGGIH

FLuc SecM(*Ec*) *N* = 495

MEDAKNIKKGPAPFYPLEDGTAGEQLHKAMKRYALVPGTIAFTDAHIEVDITYAEYFEMSVRLAEAMKRYGLNTNHRIVVC
SENSLQFFMPVLGALFIGVAVAPANDIYNERELLNSMGISQPTVVFVSKKGLQKILNVQKKLPIIQKIIIMDSKTDYQGFQS
MYTFVTSHLPPGFNEYDFVPESFDRDKTIALIMNSSGSTGLPKGVALPHRTACVRFSHARDPIFGNQIIPDTAILSVVPFHH
GFGMFTTLGYLICGFRVVLMYRFEEELFLRSLQDYKIQSALLVPTLFSFFAKSTLIDKYDLSNLHEIASGGAPLSKEVGEAVA
KRFHLPGIRQGYGLTETTSAILITPEGDDKPGAVGKVVPFFEAKVVDLDTGKTLGVNQRGELCVRGPMIMSGYVNNPEAT
NALIDKDGWLHSGDIAYWDEDEHFFIVDRLKSLIKYKGYQVAPAELESILLQHPNIFDAGVAGLPDDSGSFSTPVWISQA
QGIRAGPDYIKRAVGLPGDKVTYDPVSKELTIQPGCSSGQACENALPVTYSNVEPSDFVGSSDKQEGEWPTGLRLSRIG
GIH

FLuc SecM(*Ec*) *N* = 490

MEDAKNIKKGPAPFYPLEDGTAGEQLHKAMKRYALVPGTIAFTDAHIEVDITYAEYFEMSVRLAEAMKRYGLNTNHRIVVC
SENSLQFFMPVLGALFIGVAVAPANDIYNERELLNSMGISQPTVVFVSKKGLQKILNVQKKLPIIQKIIIMDSKTDYQGFQS
MYTFVTSHLPPGFNEYDFVPESFDRDKTIALIMNSSGSTGLPKGVALPHRTACVRFSHARDPIFGNQIIPDTAILSVVPFHH
GFGMFTTLGYLICGFRVVLMYRFEEELFLRSLQDYKIQSALLVPTLFSFFAKSTLIDKYDLSNLHEIASGGAPLSKEVGEAVA
KRFHLPGIRQGYGLTETTSAILITPEGDDKPGAVGKVVPFFEAKVVDLDTGKTLGVNQRGELCVRGPMIMSGYVNNPEAT
NALIDKDGWLHSGDIAYWDEDEHFFIVDRLKSLIKYKGYQVAPAELESILLQHPNIFDAGVASGSFSTPVWISQAQGIRAG
PDYIKRAVGLPGDKVTYDPVSKELTIQPGCSSGQACENALPVTYSNVEPSDFVGSSDKQEGEWPTGLRLSRIGGIH

FLuc SecM(*Ec*) *N* = 485

MEDAKNIKKGPAPFYPLEDGTAGEQLHKAMKRYALVPGTIAFTDAHIEVDITYAEYFEMSVRLAEAMKRYGLNTNHRIVVC
SENSLQFFMPVLGALFIGVAVAPANDIYNERELLNSMGISQPTVVFVSKKGLQKILNVQKKLPIIQKIIIMDSKTDYQGFQS
MYTFVTSHLPPGFNEYDFVPESFDRDKTIALIMNSSGSTGLPKGVALPHRTACVRFSHARDPIFGNQIIPDTAILSVVPFHH
GFGMFTTLGYLICGFRVVLMYRFEEELFLRSLQDYKIQSALLVPTLFSFFAKSTLIDKYDLSNLHEIASGGAPLSKEVGEAVA
KRFHLPGIRQGYGLTETTSAILITPEGDDKPGAVGKVVPFFEAKVVDLDTGKTLGVNQRGELCVRGPMIMSGYVNNPEAT
NALIDKDGWLHSGDIAYWDEDEHFFIVDRLKSLIKYKGYQVAPAELESILLQHPNIFSGSFSTPVWISQAQGIRAGPDYIK
RAVGLPGDKVTYDPVSKELTIQPGCSSGQACENALPVTYSNVEPSDFVGSSDKQEGEWPTGLRLSRIGGIH

FLuc SecM(*Ec*) *N* = 480

MEDAKNIKKGPAPFYPLEDGTAGEQLHKAMKRYALVPGTIAFTDAHIEVDITYAEYFEMSVRLAEAMKRYGLNTNHRIVVC SENSLQFFMPVLGALFIGVAVAPANDIYNERELLNSMGISQPTVVFVSKKGLQKILNVQKKLPIIQKIIIMDSKTDYQGFQS MYTFVTSHLPPGFNEYDFVPESFDRDKTIALIMNSSGSTGLPKGVALPHRTACVRFSHARDPIFGNQIIPDTAILSVVPFHH GFGMFTTLGYLICGFRVVLMYRFEEELFLRSLQDYKIQSALLVPTLFSFFAKSTLIDKYDLSNLHEIASGGAPLSKEVGEAVA KRFHLPGIRQGYGLTETTSAILITPEGDDKPGAVGKVVPFFEAKVVDLDTGKTLGVNQRGELCVRGPMIMSGYVNNPEAT NALIDKDGWLHSGDIAYWDEDEHFFIVDRLKSLIKYKGYQVAPAELESILLQSGSFSTPVWISQAQGIRAGPDYIKRAVGL PGDKVTYDPVSKELTIQPGCSSGQACENALPVTYSNVEPSDFVGSSDKQEGEWPTGLRLSRIGGIH

FLuc SecM(*Ec*) *N* = 475

MEDAKNIKKGPAPFYPLEDGTAGEQLHKAMKRYALVPGTIAFTDAHIEVDITYAEYFEMSVRLAEAMKRYGLNTNHRIVVC SENSLQFFMPVLGALFIGVAVAPANDIYNERELLNSMGISQPTVVFVSKKGLQKILNVQKKLPIIQKIIIMDSKTDYQGFQS MYTFVTSHLPPGFNEYDFVPESFDRDKTIALIMNSSGSTGLPKGVALPHRTACVRFSHARDPIFGNQIIPDTAILSVVPFHH GFGMFTTLGYLICGFRVVLMYRFEEELFLRSLQDYKIQSALLVPTLFSFFAKSTLIDKYDLSNLHEIASGGAPLSKEVGEAVA KRFHLPGIRQGYGLTETTSAILITPEGDDKPGAVGKVVPFFEAKVVDLDTGKTLGVNQRGELCVRGPMIMSGYVNNPEAT NALIDKDGWLHSGDIAYWDEDEHFFIVDRLKSLIKYKGYQVAPAELESGSFSTPVWISQAQGIRAGPDYIKRAVGLPGDK VTYDPVSKELTIQPGCSSGQACENALPVTYSNVEPSDFVGSSDKQEGEWPTGLRLSRIGGIH

FLuc SecM(*Ec*) *N* = 472

MEDAKNIKKGPAPFYPLEDGTAGEQLHKAMKRYALVPGTIAFTDAHIEVDITYAEYFEMSVRLAEAMKRYGLNTNHRIVVC SENSLQFFMPVLGALFIGVAVAPANDIYNERELLNSMGISQPTVVFVSKKGLQKILNVQKKLPIIQKIIIMDSKTDYQGFQS MYTFVTSHLPPGFNEYDFVPESFDRDKTIALIMNSSGSTGLPKGVALPHRTACVRFSHARDPIFGNQIIPDTAILSVVPFHH GFGMFTTLGYLICGFRVVLMYRFEEELFLRSLQDYKIQSALLVPTLFSFFAKSTLIDKYDLSNLHEIASGGAPLSKEVGEAVA KRFHLPGIRQGYGLTETTSAILITPEGDDKPGAVGKVVPFFEAKVVDLDTGKTLGVNQRGELCVRGPMIMSGYVNNPEAT NALIDKDGWLHSGDIAYWDEDEHFFIVDRLKSLIKYKGYQVAPASGSFSTPVWISQAQGIRAGPDYIKRAVGLPGDKVTY DPVSKELTIQPGCSSGQACENALPVTYSNVEPSDFVGSSDKQEGEWPTGLRLSRIGGIH

FLuc SecM(*Ec*) *N* = 470

MEDAKNIKKGPAPFYPLEDGTAGEQLHKAMKRYALVPGTIAFTDAHIEVDITYAEYFEMSVRLAEAMKRYGLNTNHRIVVC SENSLQFFMPVLGALFIGVAVAPANDIYNERELLNSMGISQPTVVFVSKKGLQKILNVQKKLPIIQKIIIMDSKTDYQGFQS MYTFVTSHLPPGFNEYDFVPESFDRDKTIALIMNSSGSTGLPKGVALPHRTACVRFSHARDPIFGNQIIPDTAILSVVPFHH GFGMFTTLGYLICGFRVVLMYRFEEELFLRSLQDYKIQSALLVPTLFSFFAKSTLIDKYDLSNLHEIASGGAPLSKEVGEAVA KRFHLPGIRQGYGLTETTSAILITPEGDDKPGAVGKVVPFFEAKVVDLDTGKTLGVNQRGELCVRGPMIMSGYVNNPEAT NALIDKDGWLHSGDIAYWDEDEHFFIVDRLKSLIKYKGYQVASGSFSTPVWISQAQGIRAGPDYIKRAVGLPGDKVTYDP VSKELTIQPGCSSGQACENALPVTYSNVEPSDFVGSSDKQEGEWPTGLRLSRIGGIH

FLuc SecM(*Ec*) *N* = 468

MEDAKNIKKGPAPFYPLEDGTAGEQLHKAMKRYALVPGTIAFTDAHIEVDITYAEYFEMSVRLAEAMKRYGLNTNHRIVVC SENSLQFFMPVLGALFIGVAVAPANDIYNERELLNSMGISQPTVVFVSKKGLQKILNVQKKLPIIQKIIIMDSKTDYQGFQS MYTFVTSHLPPGFNEYDFVPESFDRDKTIALIMNSSGSTGLPKGVALPHRTACVRFSHARDPIFGNQIIPDTAILSVVPFHH GFGMFTTLGYLICGFRVVLMYRFEEELFLRSLQDYKIQSALLVPTLFSFFAKSTLIDKYDLSNLHEIASGGAPLSKEVGEAVA KRFHLPGIRQGYGLTETTSAILITPEGDDKPGAVGKVVPFFEAKVVDLDTGKTLGVNQRGELCVRGPMIMSGYVNNPEAT NALIDKDGWLHSGDIAYWDEDEHFFIVDRLKSLIKYKGYQSGSFSTPVWISQAQGIRAGPDYIKRAVGLPGDKVTYDPVS KELTIQPGCSSGQACENALPVTYSNVEPSDFVGSSDKQEGEWPTGLRLSRIGGIH

FLuc SecM(*Ec*) *N* = 465

MEDAKNIKKGPAPFYPLEDGTAGEQLHKAMKRYALVPGTIAFTDAHIEVDITYAEYFEMSVRLAEAMKRYGLNTNHRIVVC SENSLQFFMPVLGALFIGVAVAPANDIYNERELLNSMGISQPTVVFVSKKGLQKILNVQKKLPIIQKIIIMDSKTDYQGFQS MYTFVTSHLPPGFNEYDFVPESFDRDKTIALIMNSSGSTGLPKGVALPHRTACVRFSHARDPIFGNQIIPDTAILSVVPFHH GFGMFTTLGYLICGFRVVLMYRFEEELFLRSLQDYKIQSALLVPTLFSFFAKSTLIDKYDLSNLHEIASGGAPLSKEVGEAVA

KRFHLPGIRQGYGLTETTSAILITPEGDDKPGAVGKVVPFFEAKVVDLDTGKTLGVNQRGELCVRGPMIMSGYVNNPEAT
NALIDKDGWLHSGDIAYWDEDEHFFIVDRLKSLIKYKSGSFSTPVWISQAQGIRAGPDYIKRAVGLPGDKVTYDPVSKELT
IQPGCSSGQACENALPVTYSNVEPSDFVGSSDKQEGEWPTGLRLSRIGGIH

FLuc SecM(*Ec*) *N* = 460

MEDAKNIKKGPAPFYPLEDGTAGEQLHKAMKRYALVPGTIAFTDAHIEVDITYAEYFEMSVRLAEAMKRYGLNTNHRIVVC
SENSLQFFMPVLGALFIGVAVAPANDIYNERELLNSMGISQPTVVFVSKKGLQKILNVQKKLPIIQKIIIMDSKTDYQGFQS
MYTFVTSHLPPGFNEYDFVPESFDRDKTIALIMNSSGSTGLPKGVALPHRTACVRFSHARDPIFGNQIIPDTAILSVVPFHH
GFGMFTTLGYLICGFRVVLMYRFEEELFLRSLQDYKIQSALLVPTLFSFFAKSTLIDKYDLSNLHEIASGGAPLSKEVGEAVA
KRFHLPGIRQGYGLTETTSAILITPEGDDKPGAVGKVVPFFEAKVVDLDTGKTLGVNQRGELCVRGPMIMSGYVNNPEAT
NALIDKDGWLHSGDIAYWDEDEHFFIVDRLKSSGSFSTPVWISQAQGIRAGPDYIKRAVGLPGDKVTYDPVSKELTIQPG
CSSGQACENALPVTYSNVEPSDFVGSSDKQEGEWPTGLRLSRIGGIH

FLuc SecM(*Ec*) *N* = 455

MEDAKNIKKGPAPFYPLEDGTAGEQLHKAMKRYALVPGTIAFTDAHIEVDITYAEYFEMSVRLAEAMKRYGLNTNHRIVVC
SENSLQFFMPVLGALFIGVAVAPANDIYNERELLNSMGISQPTVVFVSKKGLQKILNVQKKLPIIQKIIIMDSKTDYQGFQS
MYTFVTSHLPPGFNEYDFVPESFDRDKTIALIMNSSGSTGLPKGVALPHRTACVRFSHARDPIFGNQIIPDTAILSVVPFHH
GFGMFTTLGYLICGFRVVLMYRFEEELFLRSLQDYKIQSALLVPTLFSFFAKSTLIDKYDLSNLHEIASGGAPLSKEVGEAVA
KRFHLPGIRQGYGLTETTSAILITPEGDDKPGAVGKVVPFFEAKVVDLDTGKTLGVNQRGELCVRGPMIMSGYVNNPEAT
NALIDKDGWLHSGDIAYWDEDEHFFIVSGSFSTPVWISQAQGIRAGPDYIKRAVGLPGDKVTYDPVSKELTIQPGCSSG
QACENALPVTYSNVEPSDFVGSSDKQEGEWPTGLRLSRIGGIH

FLuc SecM(*Ec*) *N* = 450

MEDAKNIKKGPAPFYPLEDGTAGEQLHKAMKRYALVPGTIAFTDAHIEVDITYAEYFEMSVRLAEAMKRYGLNTNHRIVVC
SENSLQFFMPVLGALFIGVAVAPANDIYNERELLNSMGISQPTVVFVSKKGLQKILNVQKKLPIIQKIIIMDSKTDYQGFQS
MYTFVTSHLPPGFNEYDFVPESFDRDKTIALIMNSSGSTGLPKGVALPHRTACVRFSHARDPIFGNQIIPDTAILSVVPFHH
GFGMFTTLGYLICGFRVVLMYRFEEELFLRSLQDYKIQSALLVPTLFSFFAKSTLIDKYDLSNLHEIASGGAPLSKEVGEAVA
KRFHLPGIRQGYGLTETTSAILITPEGDDKPGAVGKVVPFFEAKVVDLDTGKTLGVNQRGELCVRGPMIMSGYVNNPEAT
NALIDKDGWLHSGDIAYWDEDESGSFSTPVWISQAQGIRAGPDYIKRAVGLPGDKVTYDPVSKELTIQPGCSSGQACEN
ALPVTYSNVEPSDFVGSSDKQEGEWPTGLRLSRIGGIH

FLuc SecM(*Ec*) *N* = 445

MEDAKNIKKGPAPFYPLEDGTAGEQLHKAMKRYALVPGTIAFTDAHIEVDITYAEYFEMSVRLAEAMKRYGLNTNHRIVVC
SENSLQFFMPVLGALFIGVAVAPANDIYNERELLNSMGISQPTVVFVSKKGLQKILNVQKKLPIIQKIIIMDSKTDYQGFQS
MYTFVTSHLPPGFNEYDFVPESFDRDKTIALIMNSSGSTGLPKGVALPHRTACVRFSHARDPIFGNQIIPDTAILSVVPFHH
GFGMFTTLGYLICGFRVVLMYRFEEELFLRSLQDYKIQSALLVPTLFSFFAKSTLIDKYDLSNLHEIASGGAPLSKEVGEAVA
KRFHLPGIRQGYGLTETTSAILITPEGDDKPGAVGKVVPFFEAKVVDLDTGKTLGVNQRGELCVRGPMIMSGYVNNPEAT
NALIDKDGWLHSGDIAYSGSFSTPVWISQAQGIRAGPDYIKRAVGLPGDKVTYDPVSKELTIQPGCSSGQACENALPVTY
SNVEPSDFVGSSDKQEGEWPTGLRLSRIGGIH

FLuc SecM(*Ec*) *N* = 440

MEDAKNIKKGPAPFYPLEDGTAGEQLHKAMKRYALVPGTIAFTDAHIEVDITYAEYFEMSVRLAEAMKRYGLNTNHRIVVC
SENSLQFFMPVLGALFIGVAVAPANDIYNERELLNSMGISQPTVVFVSKKGLQKILNVQKKLPIIQKIIIMDSKTDYQGFQS
MYTFVTSHLPPGFNEYDFVPESFDRDKTIALIMNSSGSTGLPKGVALPHRTACVRFSHARDPIFGNQIIPDTAILSVVPFHH
GFGMFTTLGYLICGFRVVLMYRFEEELFLRSLQDYKIQSALLVPTLFSFFAKSTLIDKYDLSNLHEIASGGAPLSKEVGEAVA
KRFHLPGIRQGYGLTETTSAILITPEGDDKPGAVGKVVPFFEAKVVDLDTGKTLGVNQRGELCVRGPMIMSGYVNNPEAT
NALIDKDGWLHSSGSFSTPVWISQAQGIRAGPDYIKRAVGLPGDKVTYDPVSKELTIQPGCSSGQACENALPVTYSNVE
PSDFVGSSDKQEGEWPTGLRLSRIGGIH

FLuc SecM(*Ec*) *N* = 435

MEDAKNIKKGPAPFYPLEDGTAGEQLHKAMKRYALVPGTIAFTDAHIEVDITYAEYFEMSVRLAEAMKRYGLNTNHRIVVC
SENSLQFFMPVLGALFIGVAVAPANDIYNERELLNSMGISQPTVVFVSKKGLQKILNVQKKLPIIQKIIIMDSKTDYQGFQS
MYTFVTSHLPPGFNEYDFVPESFDRDKTIALIMNSSGSTGLPKGVALPHRTACVRFSHARDPIFGNQIIPDTAILSVVPFHH
GFGMFTTLGYLICGFRVVLMYRFEEELFLRSLQDYKIQSALLVPTLFSFFAKSTLIDKYDLSNLHEIASGGAPLSKEVGEAVA
KRFHLPGIRQGYGLTETTSAILITPEGDDKPGAVGKVVPFFEAKVVDLDTGKTLGVNQRGELCVRGPMIMSGYVNNPEAT
NALIDKDSGSFSTPVWISQAQGIRAGPDYIKRAVGLPGDKVTYDPVSKELTIQPGCSSGQACENALPVTYSNVEPSDFVG
SSDKQEGEWPTGLRLSRIGGIH

FLuc SecM(*Ec*) *N* = 430

MEDAKNIKKGPAPFYPLEDGTAGEQLHKAMKRYALVPGTIAFTDAHIEVDITYAEYFEMSVRLAEAMKRYGLNTNHRIVVC
SENSLQFFMPVLGALFIGVAVAPANDIYNERELLNSMGISQPTVVFVSKKGLQKILNVQKKLPIIQKIIIMDSKTDYQGFQS
MYTFVTSHLPPGFNEYDFVPESFDRDKTIALIMNSSGSTGLPKGVALPHRTACVRFSHARDPIFGNQIIPDTAILSVVPFHH
GFGMFTTLGYLICGFRVVLMYRFEEELFLRSLQDYKIQSALLVPTLFSFFAKSTLIDKYDLSNLHEIASGGAPLSKEVGEAVA
KRFHLPGIRQGYGLTETTSAILITPEGDDKPGAVGKVVPFFEAKVVDLDTGKTLGVNQRGELCVRGPMIMSGYVNNPEAT
NASGSFSTPVWISQAQGIRAGPDYIKRAVGLPGDKVTYDPVSKELTIQPGCSSGQACENALPVTYSNVEPSDFVGSSDK
QEGEWPTGLRLSRIGGIH

FLuc SecM(*Ec*) *N* = 427

MEDAKNIKKGPAPFYPLEDGTAGEQLHKAMKRYALVPGTIAFTDAHIEVDITYAEYFEMSVRLAEAMKRYGLNTNHRIVVC
SENSLQFFMPVLGALFIGVAVAPANDIYNERELLNSMGISQPTVVFVSKKGLQKILNVQKKLPIIQKIIIMDSKTDYQGFQS
MYTFVTSHLPPGFNEYDFVPESFDRDKTIALIMNSSGSTGLPKGVALPHRTACVRFSHARDPIFGNQIIPDTAILSVVPFHH
GFGMFTTLGYLICGFRVVLMYRFEEELFLRSLQDYKIQSALLVPTLFSFFAKSTLIDKYDLSNLHEIASGGAPLSKEVGEAVA
KRFHLPGIRQGYGLTETTSAILITPEGDDKPGAVGKVVPFFEAKVVDLDTGKTLGVNQRGELCVRGPMIMSGYVNNPEAS
GSFSTPVWISQAQGIRAGPDYIKRAVGLPGDKVTYDPVSKELTIQPGCSSGQACENALPVTYSNVEPSDFVGSSDKQEG
EWPTGLRLSRIGGIH

FLuc SecM(*Ec*) *N* = 425

MEDAKNIKKGPAPFYPLEDGTAGEQLHKAMKRYALVPGTIAFTDAHIEVDITYAEYFEMSVRLAEAMKRYGLNTNHRIVVC
SENSLQFFMPVLGALFIGVAVAPANDIYNERELLNSMGISQPTVVFVSKKGLQKILNVQKKLPIIQKIIIMDSKTDYQGFQS
MYTFVTSHLPPGFNEYDFVPESFDRDKTIALIMNSSGSTGLPKGVALPHRTACVRFSHARDPIFGNQIIPDTAILSVVPFHH
GFGMFTTLGYLICGFRVVLMYRFEEELFLRSLQDYKIQSALLVPTLFSFFAKSTLIDKYDLSNLHEIASGGAPLSKEVGEAVA
KRFHLPGIRQGYGLTETTSAILITPEGDDKPGAVGKVVPFFEAKVVDLDTGKTLGVNQRGELCVRGPMIMSGYVNNPSGS
FSTPVWISQAQGIRAGPDYIKRAVGLPGDKVTYDPVSKELTIQPGCSSGQACENALPVTYSNVEPSDFVGSSDKQEGEW
PTGLRLSRIGGIH

FLuc SecM(*Ec*) *N* = 423

MEDAKNIKKGPAPFYPLEDGTAGEQLHKAMKRYALVPGTIAFTDAHIEVDITYAEYFEMSVRLAEAMKRYGLNTNHRIVVC
SENSLQFFMPVLGALFIGVAVAPANDIYNERELLNSMGISQPTVVFVSKKGLQKILNVQKKLPIIQKIIIMDSKTDYQGFQS
MYTFVTSHLPPGFNEYDFVPESFDRDKTIALIMNSSGSTGLPKGVALPHRTACVRFSHARDPIFGNQIIPDTAILSVVPFHH
GFGMFTTLGYLICGFRVVLMYRFEEELFLRSLQDYKIQSALLVPTLFSFFAKSTLIDKYDLSNLHEIASGGAPLSKEVGEAVA
KRFHLPGIRQGYGLTETTSAILITPEGDDKPGAVGKVVPFFEAKVVDLDTGKTLGVNQRGELCVRGPMIMSGYVNSGSFS
TPVWISQAQGIRAGPDYIKRAVGLPGDKVTYDPVSKELTIQPGCSSGQACENALPVTYSNVEPSDFVGSSDKQEGEWPT
GLRLSRIGGIH

FLuc SecM(*Ec*) *N* = 420

MEDAKNIKKGPAPFYPLEDGTAGEQLHKAMKRYALVPGTIAFTDAHIEVDITYAEYFEMSVRLAEAMKRYGLNTNHRIVVC
SENSLQFFMPVLGALFIGVAVAPANDIYNERELLNSMGISQPTVVFVSKKGLQKILNVQKKLPIIQKIIIMDSKTDYQGFQS
MYTFVTSHLPPGFNEYDFVPESFDRDKTIALIMNSSGSTGLPKGVALPHRTACVRFSHARDPIFGNQIIPDTAILSVVPFHH
GFGMFTTLGYLICGFRVVLMYRFEEELFLRSLQDYKIQSALLVPTLFSFFAKSTLIDKYDLSNLHEIASGGAPLSKEVGEAVA
KRFHLPGIRQGYGLTETTSAILITPEGDDKPGAVGKVVPFFEAKVVDLDTGKTLGVNQRGELCVRGPMIMSGSGSFSTPV

WISQAQGIRAGPDYIKRAVGLPGDKVTYDPVSKELTIQPGCSSGQACENALPVTYSNVEPSDFVGSSDKQEGEWPTGLRLSRIGGIH

FLuc SecM(*Ec*) *N* = 415

MEDAKNIKKGPAPFYPLEDGTAGEQLHKAMKRYALVPGTIAFTDAHIEVDITYAEYFEMSVRLAEAMKRYGLNTNHRIVVCSENSLQFFMPVLGALFIGVAVAPANDIYNERELLNSMGISQPTVVFVSKKGLQKILNVQKKLPIIQKIIIMDSKTDYQGFQSMYTFVTSHLPPGFNEYDFVPESFDRDKTIALIMNSSGSTGLPKGVALPHRTACVRFSHARDPIFGNQIIPDTAILSVVPFHHGFGMFTTLGYLICGFRVVLMYRFEEELFLRSLQDYKIQSALLVPTLFSFFAKSTLIDKYDLSNLHEIASGGAPLSKEVGEAVAKRFHLPGIRQGYGLTETTSAILITPEGDDKPGAVGKVVPFFEAKVVDLDTGKTLGVNQRGELCVRGPSGSFSTPVWISQAQGIRAGPDYIKRAVGLPGDKVTYDPVSKELTIQPGCSSGQACENALPVTYSNVEPSDFVGSSDKQEGEWPTGLRLSRIGGIH

FLuc SecM(*Ec*) *N* = 412

MEDAKNIKKGPAPFYPLEDGTAGEQLHKAMKRYALVPGTIAFTDAHIEVDITYAEYFEMSVRLAEAMKRYGLNTNHRIVVCSENSLQFFMPVLGALFIGVAVAPANDIYNERELLNSMGISQPTVVFVSKKGLQKILNVQKKLPIIQKIIIMDSKTDYQGFQSMYTFVTSHLPPGFNEYDFVPESFDRDKTIALIMNSSGSTGLPKGVALPHRTACVRFSHARDPIFGNQIIPDTAILSVVPFHHGFGMFTTLGYLICGFRVVLMYRFEEELFLRSLQDYKIQSALLVPTLFSFFAKSTLIDKYDLSNLHEIASGGAPLSKEVGEAVAKRFHLPGIRQGYGLTETTSAILITPEGDDKPGAVGKVVPFFEAKVVDLDTGKTLGVNQRGELCVSGSFSTPVWISQAQGIRAGPDYIKRAVGLPGDKVTYDPVSKELTIQPGCSSGQACENALPVTYSNVEPSDFVGSSDKQEGEWPTGLRLSRIGGIH

FLuc SecM(*Ec*) *N* = 410

MEDAKNIKKGPAPFYPLEDGTAGEQLHKAMKRYALVPGTIAFTDAHIEVDITYAEYFEMSVRLAEAMKRYGLNTNHRIVVCSENSLQFFMPVLGALFIGVAVAPANDIYNERELLNSMGISQPTVVFVSKKGLQKILNVQKKLPIIQKIIIMDSKTDYQGFQSMYTFVTSHLPPGFNEYDFVPESFDRDKTIALIMNSSGSTGLPKGVALPHRTACVRFSHARDPIFGNQIIPDTAILSVVPFHHGFGMFTTLGYLICGFRVVLMYRFEEELFLRSLQDYKIQSALLVPTLFSFFAKSTLIDKYDLSNLHEIASGGAPLSKEVGEAVAKRFHLPGIRQGYGLTETTSAILITPEGDDKPGAVGKVVPFFEAKVVDLDTGKTLGVNQRGELSGSFSTPVWISQAQGIRAGPDYIKRAVGLPGDKVTYDPVSKELTIQPGCSSGQACENALPVTYSNVEPSDFVGSSDKQEGEWPTGLRLSRIGGIH

FLuc SecM(*Ec*) *N* = 408

MEDAKNIKKGPAPFYPLEDGTAGEQLHKAMKRYALVPGTIAFTDAHIEVDITYAEYFEMSVRLAEAMKRYGLNTNHRIVVCSENSLQFFMPVLGALFIGVAVAPANDIYNERELLNSMGISQPTVVFVSKKGLQKILNVQKKLPIIQKIIIMDSKTDYQGFQSMYTFVTSHLPPGFNEYDFVPESFDRDKTIALIMNSSGSTGLPKGVALPHRTACVRFSHARDPIFGNQIIPDTAILSVVPFHHGFGMFTTLGYLICGFRVVLMYRFEEELFLRSLQDYKIQSALLVPTLFSFFAKSTLIDKYDLSNLHEIASGGAPLSKEVGEAVAKRFHLPGIRQGYGLTETTSAILITPEGDDKPGAVGKVVPFFEAKVVDLDTGKTLGVNQRGSGSFSTPVWISQAQGIRAGPDYIKRAVGLPGDKVTYDPVSKELTIQPGCSSGQACENALPVTYSNVEPSDFVGSSDKQEGEWPTGLRLSRIGGIH

FLuc SecM(*Ec*) *N* = 405

MEDAKNIKKGPAPFYPLEDGTAGEQLHKAMKRYALVPGTIAFTDAHIEVDITYAEYFEMSVRLAEAMKRYGLNTNHRIVVCSENSLQFFMPVLGALFIGVAVAPANDIYNERELLNSMGISQPTVVFVSKKGLQKILNVQKKLPIIQKIIIMDSKTDYQGFQSMYTFVTSHLPPGFNEYDFVPESFDRDKTIALIMNSSGSTGLPKGVALPHRTACVRFSHARDPIFGNQIIPDTAILSVVPFHHGFGMFTTLGYLICGFRVVLMYRFEEELFLRSLQDYKIQSALLVPTLFSFFAKSTLIDKYDLSNLHEIASGGAPLSKEVGEAVAKRFHLPGIRQGYGLTETTSAILITPEGDDKPGAVGKVVPFFEAKVVDLDTGKTLGVNSGSFSTPVWISQAQGIRAGPDYIKRAVGLPGDKVTYDPVSKELTIQPGCSSGQACENALPVTYSNVEPSDFVGSSDKQEGEWPTGLRLSRIGGIH

FLuc SecM(*Ec*) *N* = 400

MEDAKNIKKGPAPFYPLEDGTAGEQLHKAMKRYALVPGTIAFTDAHIEVDITYAEYFEMSVRLAEAMKRYGLNTNHRIVVCSENSLQFFMPVLGALFIGVAVAPANDIYNERELLNSMGISQPTVVFVSKKGLQKILNVQKKLPIIQKIIIMDSKTDYQGFQSMYTFVTSHLPPGFNEYDFVPESFDRDKTIALIMNSSGSTGLPKGVALPHRTACVRFSHARDPIFGNQIIPDTAILSVVPFHHGFGMFTTLGYLICGFRVVLMYRFEEELFLRSLQDYKIQSALLVPTLFSFFAKSTLIDKYDLSNLHEIASGGAPLSKEVGEAVAKRFHLPGIRQGYGLTETTSAILITPEGDDKPGAVGKVVPFFEAKVVDLDTGKSGSFSTPVWISQAQGIRAGPDYIKRAVGLPGDKVTYDPVSKELTIQPGCSSGQACENALPVTYSNVEPSDFVGSSDKQEGEWPTGLRLSRIGGIH

FLuc SecM(*Ec*) *N* = 395

MEDAKNIKKGPAPFYPLEDGTAGEQLHKAMKRYALVPGTIAFTDAHIEVDITYAEYFEMSVRLAEAMKRYGLNTNHRIVVC
SENSLQFFMPVLGALFIGVAVAPANDIYNERELLNSMGISQPTVVFVSKKGLQKILNVQKKLPIIQKIIIMDSKTDYQGFQS
MYTFVTSHLPPGFNEYDFVPESFDRDKTIALIMNSSGSTGLPKGVALPHRTACVRFSHARDPIFGNQIIPDTAILSVVPFHH
GFGMFTTLGYLICGFRVVLMYRFEEELFLRSLQDYKIQSALLVPTLFSFFAKSTLIDKYDLSNLHEIASGGAPLSKEVGEAVA
KRFHLPGIRQGYGLTETTSAILITPEGDDKPGAVGKVVPFFEAKVVDSGSFSTPVWISQAQGIRAGPDYIKRAVGLPGDKVT
YDPVSKELTIQPGCSSGQACENALPVTYSNVEPSDFVGSSDKQEGEWPTGLRLSRIGGIH

FLuc SecM(*Ec*) *N* = 390

MEDAKNIKKGPAPFYPLEDGTAGEQLHKAMKRYALVPGTIAFTDAHIEVDITYAEYFEMSVRLAEAMKRYGLNTNHRIVVC
SENSLQFFMPVLGALFIGVAVAPANDIYNERELLNSMGISQPTVVFVSKKGLQKILNVQKKLPIIQKIIIMDSKTDYQGFQS
MYTFVTSHLPPGFNEYDFVPESFDRDKTIALIMNSSGSTGLPKGVALPHRTACVRFSHARDPIFGNQIIPDTAILSVVPFHH
GFGMFTTLGYLICGFRVVLMYRFEEELFLRSLQDYKIQSALLVPTLFSFFAKSTLIDKYDLSNLHEIASGGAPLSKEVGEAVA
KRFHLPGIRQGYGLTETTSAILITPEGDDKPGAVGKVVPFFESGSFSTPVWISQAQGIRAGPDYIKRAVGLPGDKVTYDPVS
KELTIQPGCSSGQACENALPVTYSNVEPSDFVGSSDKQEGEWPTGLRLSRIGGIH

FLuc SecM(*Ec*) *N* = 385

MEDAKNIKKGPAPFYPLEDGTAGEQLHKAMKRYALVPGTIAFTDAHIEVDITYAEYFEMSVRLAEAMKRYGLNTNHRIVVC
SENSLQFFMPVLGALFIGVAVAPANDIYNERELLNSMGISQPTVVFVSKKGLQKILNVQKKLPIIQKIIIMDSKTDYQGFQS
MYTFVTSHLPPGFNEYDFVPESFDRDKTIALIMNSSGSTGLPKGVALPHRTACVRFSHARDPIFGNQIIPDTAILSVVPFHH
GFGMFTTLGYLICGFRVVLMYRFEEELFLRSLQDYKIQSALLVPTLFSFFAKSTLIDKYDLSNLHEIASGGAPLSKEVGEAVA
KRFHLPGIRQGYGLTETTSAILITPEGDDKPGAVGKVSGSFSTPVWISQAQGIRAGPDYIKRAVGLPGDKVTYDPVSKELTI
QPGCSSGQACENALPVTYSNVEPSDFVGSSDKQEGEWPTGLRLSRIGGIH

FLuc SecM(*Ec*) *N* = 382

MEDAKNIKKGPAPFYPLEDGTAGEQLHKAMKRYALVPGTIAFTDAHIEVDITYAEYFEMSVRLAEAMKRYGLNTNHRIVVC
SENSLQFFMPVLGALFIGVAVAPANDIYNERELLNSMGISQPTVVFVSKKGLQKILNVQKKLPIIQKIIIMDSKTDYQGFQS
MYTFVTSHLPPGFNEYDFVPESFDRDKTIALIMNSSGSTGLPKGVALPHRTACVRFSHARDPIFGNQIIPDTAILSVVPFHH
GFGMFTTLGYLICGFRVVLMYRFEEELFLRSLQDYKIQSALLVPTLFSFFAKSTLIDKYDLSNLHEIASGGAPLSKEVGEAVA
KRFHLPGIRQGYGLTETTSAILITPEGDDKPGAVSGSFSTPVWISQAQGIRAGPDYIKRAVGLPGDKVTYDPVSKELTIQPG
CSSGQACENALPVTYSNVEPSDFVGSSDKQEGEWPTGLRLSRIGGIH

FLuc SecM(*Ec*) *N* = 380

MEDAKNIKKGPAPFYPLEDGTAGEQLHKAMKRYALVPGTIAFTDAHIEVDITYAEYFEMSVRLAEAMKRYGLNTNHRIVVC
SENSLQFFMPVLGALFIGVAVAPANDIYNERELLNSMGISQPTVVFVSKKGLQKILNVQKKLPIIQKIIIMDSKTDYQGFQS
MYTFVTSHLPPGFNEYDFVPESFDRDKTIALIMNSSGSTGLPKGVALPHRTACVRFSHARDPIFGNQIIPDTAILSVVPFHH
GFGMFTTLGYLICGFRVVLMYRFEEELFLRSLQDYKIQSALLVPTLFSFFAKSTLIDKYDLSNLHEIASGGAPLSKEVGEAVA
KRFHLPGIRQGYGLTETTSAILITPEGDDKPGSGSFSTPVWISQAQGIRAGPDYIKRAVGLPGDKVTYDPVSKELTIQPGCS
SGQACENALPVTYSNVEPSDFVGSSDKQEGEWPTGLRLSRIGGIH

FLuc SecM(*Ec*) *N* = 378

MEDAKNIKKGPAPFYPLEDGTAGEQLHKAMKRYALVPGTIAFTDAHIEVDITYAEYFEMSVRLAEAMKRYGLNTNHRIVVC
SENSLQFFMPVLGALFIGVAVAPANDIYNERELLNSMGISQPTVVFVSKKGLQKILNVQKKLPIIQKIIIMDSKTDYQGFQS
MYTFVTSHLPPGFNEYDFVPESFDRDKTIALIMNSSGSTGLPKGVALPHRTACVRFSHARDPIFGNQIIPDTAILSVVPFHH
GFGMFTTLGYLICGFRVVLMYRFEEELFLRSLQDYKIQSALLVPTLFSFFAKSTLIDKYDLSNLHEIASGGAPLSKEVGEAVA
KRFHLPGIRQGYGLTETTSAILITPEGDDKSGSFSTPVWISQAQGIRAGPDYIKRAVGLPGDKVTYDPVSKELTIQPGCSSG
QACENALPVTYSNVEPSDFVGSSDKQEGEWPTGLRLSRIGGIH

FLuc SecM(*Ec*) *N* = 375

MEDAKNIKKGPAPFYPLEDGTAGEQLHKAMKRYALVPGTIAFTDAHIEVDITYAEYFEMSVRLAEAMKRYGLNTNHRIVVC
SENSLQFFMPVLGALFIGVAVAPANDIYNERELLNSMGISQPTVVFVSKKGLQKILNVQKKLPIIQKIIIMDSKTDYQGFQS
MYTFVTSHLPPGFNEYDFVPESFDRDKTIALIMNSSGSTGLPKGVALPHRTACVRFSHARDPIFGNQIIPDTAILSVVPFHH
GFGMFTTLGYLICGFRVVLMYRFEEELFLRSLQDYKIQSALLVPTLFSFFAKSTLIDKYDLSNLHEIASGGAPLSKEVGEAVA
KRFHLPGIRQGYGLTETTSAILITPEGSGSFSTPVWISQAQGIRAGPDYIKRAVGLPGDKVTYDPVSKELTIQPGCSSGQAC
ENALPVTYSNVEPSDFVGSSDKQEGEWPTGLRLSRIGGIH

FLuc SecM(*Ec*) $N$ = 370

MEDAKNIKKGPAPFYPLEDGTAGEQLHKAMKRYALVPGTIAFTDAHIEVDITYAEYFEMSVRLAEAMKRYGLNTNHRIVVC
SENSLQFFMPVLGALFIGVAVAPANDIYNERELLNSMGISQPTVVFVSKKGLQKILNVQKKLPIIQKIIIMDSKTDYQGFQS
MYTFVTSHLPPGFNEYDFVPESFDRDKTIALIMNSSGSTGLPKGVALPHRTACVRFSHARDPIFGNQIIPDTAILSVVPFHH
GFGMFTTLGYLICGFRVVLMYRFEEELFLRSLQDYKIQSALLVPTLFSFFAKSTLIDKYDLSNLHEIASGGAPLSKEVGEAVA
KRFHLPGIRQGYGLTETTSAILSGSFSTPVWISQAQGIRAGPDYIKRAVGLPGDKVTYDPVSKELTIQPGCSSGQACENAL
PVTYSNVEPSDFVGSSDKQEGEWPTGLRLSRIGGIH

FLuc SecM(*Ec*) $N$ = 365

MEDAKNIKKGPAPFYPLEDGTAGEQLHKAMKRYALVPGTIAFTDAHIEVDITYAEYFEMSVRLAEAMKRYGLNTNHRIVVC
SENSLQFFMPVLGALFIGVAVAPANDIYNERELLNSMGISQPTVVFVSKKGLQKILNVQKKLPIIQKIIIMDSKTDYQGFQS
MYTFVTSHLPPGFNEYDFVPESFDRDKTIALIMNSSGSTGLPKGVALPHRTACVRFSHARDPIFGNQIIPDTAILSVVPFHH
GFGMFTTLGYLICGFRVVLMYRFEEELFLRSLQDYKIQSALLVPTLFSFFAKSTLIDKYDLSNLHEIASGGAPLSKEVGEAVA
KRFHLPGIRQGYGLTETSGSFSTPVWISQAQGIRAGPDYIKRAVGLPGDKVTYDPVSKELTIQPGCSSGQACENALPVTYS
NVEPSDFVGSSDKQEGEWPTGLRLSRIGGIH

FLuc SecM(*Ec*) $N$ = 360

MEDAKNIKKGPAPFYPLEDGTAGEQLHKAMKRYALVPGTIAFTDAHIEVDITYAEYFEMSVRLAEAMKRYGLNTNHRIVVC
SENSLQFFMPVLGALFIGVAVAPANDIYNERELLNSMGISQPTVVFVSKKGLQKILNVQKKLPIIQKIIIMDSKTDYQGFQS
MYTFVTSHLPPGFNEYDFVPESFDRDKTIALIMNSSGSTGLPKGVALPHRTACVRFSHARDPIFGNQIIPDTAILSVVPFHH
GFGMFTTLGYLICGFRVVLMYRFEEELFLRSLQDYKIQSALLVPTLFSFFAKSTLIDKYDLSNLHEIASGGAPLSKEVGEAVA
KRFHLPGIRQGYSGSFSTPVWISQAQGIRAGPDYIKRAVGLPGDKVTYDPVSKELTIQPGCSSGQACENALPVTYSNVEP
SDFVGSSDKQEGEWPTGLRLSRIGGIH

FLuc SecM(*Ec*) $N$ = 355

MEDAKNIKKGPAPFYPLEDGTAGEQLHKAMKRYALVPGTIAFTDAHIEVDITYAEYFEMSVRLAEAMKRYGLNTNHRIVVC
SENSLQFFMPVLGALFIGVAVAPANDIYNERELLNSMGISQPTVVFVSKKGLQKILNVQKKLPIIQKIIIMDSKTDYQGFQS
MYTFVTSHLPPGFNEYDFVPESFDRDKTIALIMNSSGSTGLPKGVALPHRTACVRFSHARDPIFGNQIIPDTAILSVVPFHH
GFGMFTTLGYLICGFRVVLMYRFEEELFLRSLQDYKIQSALLVPTLFSFFAKSTLIDKYDLSNLHEIASGGAPLSKEVGEAVA
KRFHLPGSGSFSTPVWISQAQGIRAGPDYIKRAVGLPGDKVTYDPVSKELTIQPGCSSGQACENALPVTYSNVEPSDFVG
SSDKQEGEWPTGLRLSRIGGIH

FLuc SecM(*Ec*) $N$ = 350

MEDAKNIKKGPAPFYPLEDGTAGEQLHKAMKRYALVPGTIAFTDAHIEVDITYAEYFEMSVRLAEAMKRYGLNTNHRIVVC
SENSLQFFMPVLGALFIGVAVAPANDIYNERELLNSMGISQPTVVFVSKKGLQKILNVQKKLPIIQKIIIMDSKTDYQGFQS
MYTFVTSHLPPGFNEYDFVPESFDRDKTIALIMNSSGSTGLPKGVALPHRTACVRFSHARDPIFGNQIIPDTAILSVVPFHH
GFGMFTTLGYLICGFRVVLMYRFEEELFLRSLQDYKIQSALLVPTLFSFFAKSTLIDKYDLSNLHEIASGGAPLSKEVGEAVA
KRSGSFSTPVWISQAQGIRAGPDYIKRAVGLPGDKVTYDPVSKELTIQPGCSSGQACENALPVTYSNVEPSDFVGSSDKQ
EGEWPTGLRLSRIGGIH

FLuc SecM(*Ec*) $N$ = 345

MEDAKNIKKGPAPFYPLEDGTAGEQLHKAMKRYALVPGTIAFTDAHIEVDITYAEYFEMSVRLAEAMKRYGLNTNHRIVVC
SENSLQFFMPVLGALFIGVAVAPANDIYNERELLNSMGISQPTVVFVSKKGLQKILNVQKKLPIIQKIIIMDSKTDYQGFQS
MYTFVTSHLPPGFNEYDFVPESFDRDKTIALIMNSSGSTGLPKGVALPHRTACVRFSHARDPIFGNQIIPDTAILSVVPFHH

GFGMFTTLGYLICGFRVVLMYRFEEELFLRSLQDYKIQSALLVPTLFSFFAKSTLIDKYDLSNLHEIASGGAPLSKEVGESGS
FSTPVWISQAQGIRAGPDYIKRAVGLPGDKVTYDPVSKELTIQPGCSSGQACENALPVTYSNVEPSDFVGSSDKQEGEW
PTGLRLSRIGGIH

FLuc SecM(*Ec*) *N* = 340

MEDAKNIKKGPAPFYPLEDGTAGEQLHKAMKRYALVPGTIAFTDAHIEVDITYAEYFEMSVRLAEAMKRYGLNTNHRIVVC
SENSLQFFMPVLGALFIGVAVAPANDIYNERELLNSMGISQPTVVFVSKKGLQKILNVQKKLPIIQKIIIMDSKTDYQGFQS
MYTFVTSHLPPGFNEYDFVPESFDRDKTIALIMNSSGSTGLPKGVALPHRTACVRFSHARDPIFGNQIIPDTAILSVVPFHH
GFGMFTTLGYLICGFRVVLMYRFEEELFLRSLQDYKIQSALLVPTLFSFFAKSTLIDKYDLSNLHEIASGGAPLSSGSFSTPV
WISQAQGIRAGPDYIKRAVGLPGDKVTYDPVSKELTIQPGCSSGQACENALPVTYSNVEPSDFVGSSDKQEGEWPTGLR
LSRIGGIH

FLuc SecM(*Ec*) *N* = 335

MEDAKNIKKGPAPFYPLEDGTAGEQLHKAMKRYALVPGTIAFTDAHIEVDITYAEYFEMSVRLAEAMKRYGLNTNHRIVVC
SENSLQFFMPVLGALFIGVAVAPANDIYNERELLNSMGISQPTVVFVSKKGLQKILNVQKKLPIIQKIIIMDSKTDYQGFQS
MYTFVTSHLPPGFNEYDFVPESFDRDKTIALIMNSSGSTGLPKGVALPHRTACVRFSHARDPIFGNQIIPDTAILSVVPFHH
GFGMFTTLGYLICGFRVVLMYRFEEELFLRSLQDYKIQSALLVPTLFSFFAKSTLIDKYDLSNLHEIASGSGSFSTPVWISQA
QGIRAGPDYIKRAVGLPGDKVTYDPVSKELTIQPGCSSGQACENALPVTYSNVEPSDFVGSSDKQEGEWPTGLRLSRIG
GIH

FLuc SecM(*Ec*) *N* = 330

MEDAKNIKKGPAPFYPLEDGTAGEQLHKAMKRYALVPGTIAFTDAHIEVDITYAEYFEMSVRLAEAMKRYGLNTNHRIVVC
SENSLQFFMPVLGALFIGVAVAPANDIYNERELLNSMGISQPTVVFVSKKGLQKILNVQKKLPIIQKIIIMDSKTDYQGFQS
MYTFVTSHLPPGFNEYDFVPESFDRDKTIALIMNSSGSTGLPKGVALPHRTACVRFSHARDPIFGNQIIPDTAILSVVPFHH
GFGMFTTLGYLICGFRVVLMYRFEEELFLRSLQDYKIQSALLVPTLFSFFAKSTLIDKYDLSNLHSGSFSTPVWISQAQGIR
AGPDYIKRAVGLPGDKVTYDPVSKELTIQPGCSSGQACENALPVTYSNVEPSDFVGSSDKQEGEWPTGLRLSRIGGIH

FLuc SecM(*Ec*) *N* = 325

MEDAKNIKKGPAPFYPLEDGTAGEQLHKAMKRYALVPGTIAFTDAHIEVDITYAEYFEMSVRLAEAMKRYGLNTNHRIVVC
SENSLQFFMPVLGALFIGVAVAPANDIYNERELLNSMGISQPTVVFVSKKGLQKILNVQKKLPIIQKIIIMDSKTDYQGFQS
MYTFVTSHLPPGFNEYDFVPESFDRDKTIALIMNSSGSTGLPKGVALPHRTACVRFSHARDPIFGNQIIPDTAILSVVPFHH
GFGMFTTLGYLICGFRVVLMYRFEEELFLRSLQDYKIQSALLVPTLFSFFAKSTLIDKYDSGSFSTPVWISQAQGIRAGPDYI
KRAVGLPGDKVTYDPVSKELTIQPGCSSGQACENALPVTYSNVEPSDFVGSSDKQEGEWPTGLRLSRIGGIH

FLuc SecM(*Ec*) *N* = 320

MEDAKNIKKGPAPFYPLEDGTAGEQLHKAMKRYALVPGTIAFTDAHIEVDITYAEYFEMSVRLAEAMKRYGLNTNHRIVVC
SENSLQFFMPVLGALFIGVAVAPANDIYNERELLNSMGISQPTVVFVSKKGLQKILNVQKKLPIIQKIIIMDSKTDYQGFQS
MYTFVTSHLPPGFNEYDFVPESFDRDKTIALIMNSSGSTGLPKGVALPHRTACVRFSHARDPIFGNQIIPDTAILSVVPFHH
GFGMFTTLGYLICGFRVVLMYRFEEELFLRSLQDYKIQSALLVPTLFSFFAKSTLSGSFSTPVWISQAQGIRAGPDYIKRAV
GLPGDKVTYDPVSKELTIQPGCSSGQACENALPVTYSNVEPSDFVGSSDKQEGEWPTGLRLSRIGGIH

FLuc SecM(*Ec*) *N* = 317

MEDAKNIKKGPAPFYPLEDGTAGEQLHKAMKRYALVPGTIAFTDAHIEVDITYAEYFEMSVRLAEAMKRYGLNTNHRIVVC
SENSLQFFMPVLGALFIGVAVAPANDIYNERELLNSMGISQPTVVFVSKKGLQKILNVQKKLPIIQKIIIMDSKTDYQGFQS
MYTFVTSHLPPGFNEYDFVPESFDRDKTIALIMNSSGSTGLPKGVALPHRTACVRFSHARDPIFGNQIIPDTAILSVVPFHH
GFGMFTTLGYLICGFRVVLMYRFEEELFLRSLQDYKIQSALLVPTLFSFFAKSGSFSTPVWISQAQGIRAGPDYIKRAVGLP
GDKVTYDPVSKELTIQPGCSSGQACENALPVTYSNVEPSDFVGSSDKQEGEWPTGLRLSRIGGIH

FLuc SecM(*Ec*) *N* = 315

MEDAKNIKKGPAPFYPLEDGTAGEQLHKAMKRYALVPGTIAFTDAHIEVDITYAEYFEMSVRLAEAMKRYGLNTNHRIVVC
SENSLQFFMPVLGALFIGVAVAPANDIYNERELLNSMGISQPTVVFVSKKGLQKILNVQKKLPIIQKIIIMDSKTDYQGFQS
MYTFVTSHLPPGFNEYDFVPESFDRDKTIALIMNSSGSTGLPKGVALPHRTACVRFSHARDPIFGNQIIPDTAILSVVPFHH
GFGMFTTLGYLICGFRVVLMYRFEEELFLRSLQDYKIQSALLVPTLFSFFSGSFSTPVWISQAQGIRAGPDYIKRAVGLPGD
KVTYDPVSKELTIQPGCSSGQACENALPVTYSNVEPSDFVGSSDKQEGEWPTGLRLSRIGGIH

FLuc SecM(*Ec*) $N = 313$

MEDAKNIKKGPAPFYPLEDGTAGEQLHKAMKRYALVPGTIAFTDAHIEVDITYAEYFEMSVRLAEAMKRYGLNTNHRIVVC
SENSLQFFMPVLGALFIGVAVAPANDIYNERELLNSMGISQPTVVFVSKKGLQKILNVQKKLPIIQKIIIMDSKTDYQGFQS
MYTFVTSHLPPGFNEYDFVPESFDRDKTIALIMNSSGSTGLPKGVALPHRTACVRFSHARDPIFGNQIIPDTAILSVVPFHH
GFGMFTTLGYLICGFRVVLMYRFEEELFLRSLQDYKIQSALLVPTLFSSGSFSTPVWISQAQGIRAGPDYIKRAVGLPGDKV
TYDPVSKELTIQPGCSSGQACENALPVTYSNVEPSDFVGSSDKQEGEWPTGLRLSRIGGIH

FLuc SecM(*Ec*) $N = 310$

MEDAKNIKKGPAPFYPLEDGTAGEQLHKAMKRYALVPGTIAFTDAHIEVDITYAEYFEMSVRLAEAMKRYGLNTNHRIVVC
SENSLQFFMPVLGALFIGVAVAPANDIYNERELLNSMGISQPTVVFVSKKGLQKILNVQKKLPIIQKIIIMDSKTDYQGFQS
MYTFVTSHLPPGFNEYDFVPESFDRDKTIALIMNSSGSTGLPKGVALPHRTACVRFSHARDPIFGNQIIPDTAILSVVPFHH
GFGMFTTLGYLICGFRVVLMYRFEEELFLRSLQDYKIQSALLVPTSGSFSTPVWISQAQGIRAGPDYIKRAVGLPGDKVTY
DPVSKELTIQPGCSSGQACENALPVTYSNVEPSDFVGSSDKQEGEWPTGLRLSRIGGIH

FLuc SecM(*Ec*) $N = 305$

MEDAKNIKKGPAPFYPLEDGTAGEQLHKAMKRYALVPGTIAFTDAHIEVDITYAEYFEMSVRLAEAMKRYGLNTNHRIVVC
SENSLQFFMPVLGALFIGVAVAPANDIYNERELLNSMGISQPTVVFVSKKGLQKILNVQKKLPIIQKIIIMDSKTDYQGFQS
MYTFVTSHLPPGFNEYDFVPESFDRDKTIALIMNSSGSTGLPKGVALPHRTACVRFSHARDPIFGNQIIPDTAILSVVPFHH
GFGMFTTLGYLICGFRVVLMYRFEEELFLRSLQDYKIQSASGSFSTPVWISQAQGIRAGPDYIKRAVGLPGDKVTYDPVSK
ELTIQPGCSSGQACENALPVTYSNVEPSDFVGSSDKQEGEWPTGLRLSRIGGIH

FLuc SecM(*Ec*) $N = 302$

MEDAKNIKKGPAPFYPLEDGTAGEQLHKAMKRYALVPGTIAFTDAHIEVDITYAEYFEMSVRLAEAMKRYGLNTNHRIVVC
SENSLQFFMPVLGALFIGVAVAPANDIYNERELLNSMGISQPTVVFVSKKGLQKILNVQKKLPIIQKIIIMDSKTDYQGFQS
MYTFVTSHLPPGFNEYDFVPESFDRDKTIALIMNSSGSTGLPKGVALPHRTACVRFSHARDPIFGNQIIPDTAILSVVPFHH
GFGMFTTLGYLICGFRVVLMYRFEEELFLRSLQDYKISGSFSTPVWISQAQGIRAGPDYIKRAVGLPGDKVTYDPVSKELTI
QPGCSSGQACENALPVTYSNVEPSDFVGSSDKQEGEWPTGLRLSRIGGIH

FLuc SecM(*Ec*) $N = 300$

MEDAKNIKKGPAPFYPLEDGTAGEQLHKAMKRYALVPGTIAFTDAHIEVDITYAEYFEMSVRLAEAMKRYGLNTNHRIVVC
SENSLQFFMPVLGALFIGVAVAPANDIYNERELLNSMGISQPTVVFVSKKGLQKILNVQKKLPIIQKIIIMDSKTDYQGFQS
MYTFVTSHLPPGFNEYDFVPESFDRDKTIALIMNSSGSTGLPKGVALPHRTACVRFSHARDPIFGNQIIPDTAILSVVPFHH
GFGMFTTLGYLICGFRVVLMYRFEEELFLRSLQDYSGSFSTPVWISQAQGIRAGPDYIKRAVGLPGDKVTYDPVSKELTIQ
PGCSSGQACENALPVTYSNVEPSDFVGSSDKQEGEWPTGLRLSRIGGIH

FLuc SecM(*Ec*) $N = 298$

MEDAKNIKKGPAPFYPLEDGTAGEQLHKAMKRYALVPGTIAFTDAHIEVDITYAEYFEMSVRLAEAMKRYGLNTNHRIVVC
SENSLQFFMPVLGALFIGVAVAPANDIYNERELLNSMGISQPTVVFVSKKGLQKILNVQKKLPIIQKIIIMDSKTDYQGFQS
MYTFVTSHLPPGFNEYDFVPESFDRDKTIALIMNSSGSTGLPKGVALPHRTACVRFSHARDPIFGNQIIPDTAILSVVPFHH
GFGMFTTLGYLICGFRVVLMYRFEEELFLRSLQSGSFSTPVWISQAQGIRAGPDYIKRAVGLPGDKVTYDPVSKELTIQPG
CSSGQACENALPVTYSNVEPSDFVGSSDKQEGEWPTGLRLSRIGGIH

FLuc SecM(*Ec*) $N = 295$

MEDAKNIKKGPAPFYPLEDGTAGEQLHKAMKRYALVPGTIAFTDAHIEVDITYAEYFEMSVRLAEAMKRYGLNTNHRIVVC
SENSLQFFMPVLGALFIGVAVAPANDIYNERELLNSMGISQPTVVFVSKKGLQKILNVQKKLPIIQKIIIMDSKTDYQGFQS
MYTFVTSHLPPGFNEYDFVPESFDRDKTIALIMNSSGSTGLPKGVALPHRTACVRFSHARDPIFGNQIIPDTAILSVVPFHH
GFGMFTTLGYLICGFRVVLMYRFEEELFLRSGSFSTPVWISQAQGIRAGPDYIKRAVGLPGDKVTYDPVSKELTIQPGCSS
GQACENALPVTYSNVEPSDFVGSSDKQEGEWPTGLRLSRIGGIH

FLuc SecM(*Ec*) *N* = 290

MEDAKNIKKGPAPFYPLEDGTAGEQLHKAMKRYALVPGTIAFTDAHIEVDITYAEYFEMSVRLAEAMKRYGLNTNHRIVVC
SENSLQFFMPVLGALFIGVAVAPANDIYNERELLNSMGISQPTVVFVSKKGLQKILNVQKKLPIIQKIIIMDSKTDYQGFQS
MYTFVTSHLPPGFNEYDFVPESFDRDKTIALIMNSSGSTGLPKGVALPHRTACVRFSHARDPIFGNQIIPDTAILSVVPFHH
GFGMFTTLGYLICGFRVVLMYRFEESGSFSTPVWISQAQGIRAGPDYIKRAVGLPGDKVTYDPVSKELTIQPGCSSGQAC
ENALPVTYSNVEPSDFVGSSDKQEGEWPTGLRLSRIGGIH

FLuc SecM(*Ec*) *N* = 285

MEDAKNIKKGPAPFYPLEDGTAGEQLHKAMKRYALVPGTIAFTDAHIEVDITYAEYFEMSVRLAEAMKRYGLNTNHRIVVC
SENSLQFFMPVLGALFIGVAVAPANDIYNERELLNSMGISQPTVVFVSKKGLQKILNVQKKLPIIQKIIIMDSKTDYQGFQS
MYTFVTSHLPPGFNEYDFVPESFDRDKTIALIMNSSGSTGLPKGVALPHRTACVRFSHARDPIFGNQIIPDTAILSVVPFHH
GFGMFTTLGYLICGFRVVLMSGSFSTPVWISQAQGIRAGPDYIKRAVGLPGDKVTYDPVSKELTIQPGCSSGQACENALP
VTYSNVEPSDFVGSSDKQEGEWPTGLRLSRIGGIH

FLuc SecM(*Ec*) *N* = 280

MEDAKNIKKGPAPFYPLEDGTAGEQLHKAMKRYALVPGTIAFTDAHIEVDITYAEYFEMSVRLAEAMKRYGLNTNHRIVVC
SENSLQFFMPVLGALFIGVAVAPANDIYNERELLNSMGISQPTVVFVSKKGLQKILNVQKKLPIIQKIIIMDSKTDYQGFQS
MYTFVTSHLPPGFNEYDFVPESFDRDKTIALIMNSSGSTGLPKGVALPHRTACVRFSHARDPIFGNQIIPDTAILSVVPFHH
GFGMFTTLGYLICGFSGSFSTPVWISQAQGIRAGPDYIKRAVGLPGDKVTYDPVSKELTIQPGCSSGQACENALPVTYSN
VEPSDFVGSSDKQEGEWPTGLRLSRIGGIH

FLuc SecM(*Ec*) *N* = 275

MEDAKNIKKGPAPFYPLEDGTAGEQLHKAMKRYALVPGTIAFTDAHIEVDITYAEYFEMSVRLAEAMKRYGLNTNHRIVVC
SENSLQFFMPVLGALFIGVAVAPANDIYNERELLNSMGISQPTVVFVSKKGLQKILNVQKKLPIIQKIIIMDSKTDYQGFQS
MYTFVTSHLPPGFNEYDFVPESFDRDKTIALIMNSSGSTGLPKGVALPHRTACVRFSHARDPIFGNQIIPDTAILSVVPFHH
GFGMFTTLGYSGSFSTPVWISQAQGIRAGPDYIKRAVGLPGDKVTYDPVSKELTIQPGCSSGQACENALPVTYSNVEPSD
FVGSSDKQEGEWPTGLRLSRIGGIH

FLuc SecM(*Ec*) *N* = 270

MEDAKNIKKGPAPFYPLEDGTAGEQLHKAMKRYALVPGTIAFTDAHIEVDITYAEYFEMSVRLAEAMKRYGLNTNHRIVVC
SENSLQFFMPVLGALFIGVAVAPANDIYNERELLNSMGISQPTVVFVSKKGLQKILNVQKKLPIIQKIIIMDSKTDYQGFQS
MYTFVTSHLPPGFNEYDFVPESFDRDKTIALIMNSSGSTGLPKGVALPHRTACVRFSHARDPIFGNQIIPDTAILSVVPFHH
GFGMFSGSFSTPVWISQAQGIRAGPDYIKRAVGLPGDKVTYDPVSKELTIQPGCSSGQACENALPVTYSNVEPSDFVGS
SDKQEGEWPTGLRLSRIGGIH

FLuc SecM(*Ec*) *N* = 265

MEDAKNIKKGPAPFYPLEDGTAGEQLHKAMKRYALVPGTIAFTDAHIEVDITYAEYFEMSVRLAEAMKRYGLNTNHRIVVC
SENSLQFFMPVLGALFIGVAVAPANDIYNERELLNSMGISQPTVVFVSKKGLQKILNVQKKLPIIQKIIIMDSKTDYQGFQS
MYTFVTSHLPPGFNEYDFVPESFDRDKTIALIMNSSGSTGLPKGVALPHRTACVRFSHARDPIFGNQIIPDTAILSVVPFHH
SGSFSTPVWISQAQGIRAGPDYIKRAVGLPGDKVTYDPVSKELTIQPGCSSGQACENALPVTYSNVEPSDFVGSSDKQE
GEWPTGLRLSRIGGIH

FLuc SecM(*Ec*) *N* = 260

MEDAKNIKKGPAPFYPLEDGTAGEQLHKAMKRYALVPGTIAFTDAHIEVDITYAEYFEMSVRLAEAMKRYGLNTNHRIVVC
SENSLQFFMPVLGALFIGVAVAPANDIYNERELLNSMGISQPTVVFVSKKGLQKILNVQKKLPIIQKIIIMDSKTDYQGFQS
MYTFVTSHLPPGFNEYDFVPESFDRDKTIALIMNSSGSTGLPKGVALPHRTACVRFSHARDPIFGNQIIPDTAILSVSGSFS
TPVWISQAQGIRAGPDYIKRAVGLPGDKVTYDPVSKELTIQPGCSSGQACENALPVTYSNVEPSDFVGSSDKQEGEWPT
GLRLSRIGGIH

FLuc SecM(*Ec*) *N* = 255

MEDAKNIKKGPAPFYPLEDGTAGEQLHKAMKRYALVPGTIAFTDAHIEVDITYAEYFEMSVRLAEAMKRYGLNTNHRIVVC
SENSLQFFMPVLGALFIGVAVAPANDIYNERELLNSMGISQPTVVFVSKKGLQKILNVQKKLPIIQKIIIMDSKTDYQGFQS
MYTFVTSHLPPGFNEYDFVPESFDRDKTIALIMNSSGSTGLPKGVALPHRTACVRFSHARDPIFGNQIIPDTSGSFSTPVWI
SQAQGIRAGPGSSDKQEGEWPTGLRLSRIGGIH

FLuc SecM(*Ec*) *N* = 250

MEDAKNIKKGPAPFYPLEDGTAGEQLHKAMKRYALVPGTIAFTDAHIEVDITYAEYFEMSVRLAEAMKRYGLNTNHRIVVC
SENSLQFFMPVLGALFIGVAVAPANDIYNERELLNSMGISQPTVVFVSKKGLQKILNVQKKLPIIQKIIIMDSKTDYQGFQS
MYTFVTSHLPPGFNEYDFVPESFDRDKTIALIMNSSGSTGLPKGVALPHRTACVRFSHARDPIFGNQSGSFSTPVWISQA
QGIRAGPGSSDKQEGEWPTGLRLSRIGGIH

FLuc SecM(*Ec*) *N* = 245

MEDAKNIKKGPAPFYPLEDGTAGEQLHKAMKRYALVPGTIAFTDAHIEVDITYAEYFEMSVRLAEAMKRYGLNTNHRIVVC
SENSLQFFMPVLGALFIGVAVAPANDIYNERELLNSMGISQPTVVFVSKKGLQKILNVQKKLPIIQKIIIMDSKTDYQGFQS
MYTFVTSHLPPGFNEYDFVPESFDRDKTIALIMNSSGSTGLPKGVALPHRTACVRFSHARDPSGSFSTPVWISQAQGIRA
GPGSSDKQEGEWPTGLRLSRIGGIH

FLuc SecM(*Ec*) *N* = 240

MEDAKNIKKGPAPFYPLEDGTAGEQLHKAMKRYALVPGTIAFTDAHIEVDITYAEYFEMSVRLAEAMKRYGLNTNHRIVVC
SENSLQFFMPVLGALFIGVAVAPANDIYNERELLNSMGISQPTVVFVSKKGLQKILNVQKKLPIIQKIIIMDSKTDYQGFQS
MYTFVTSHLPPGFNEYDFVPESFDRDKTIALIMNSSGSTGLPKGVALPHRTACVRFSSGSFSTPVWISQAQGIRAGPGSS
DKQEGEWPTGLRLSRIGGIH

FLuc SecM(*Ec*) *N* = 235

MEDAKNIKKGPAPFYPLEDGTAGEQLHKAMKRYALVPGTIAFTDAHIEVDITYAEYFEMSVRLAEAMKRYGLNTNHRIVVC
SENSLQFFMPVLGALFIGVAVAPANDIYNERELLNSMGISQPTVVFVSKKGLQKILNVQKKLPIIQKIIIMDSKTDYQGFQS
MYTFVTSHLPPGFNEYDFVPESFDRDKTIALIMNSSGSTGLPKGVALPHRTASGSFSTPVWISQAQGIRAGPGSSDKQEG
EWPTGLRLSRIGGIH

FLuc SecM(*Ec*) *N* = 230

MEDAKNIKKGPAPFYPLEDGTAGEQLHKAMKRYALVPGTIAFTDAHIEVDITYAEYFEMSVRLAEAMKRYGLNTNHRIVVC
SENSLQFFMPVLGALFIGVAVAPANDIYNERELLNSMGISQPTVVFVSKKGLQKILNVQKKLPIIQKIIIMDSKTDYQGFQS
MYTFVTSHLPPGFNEYDFVPESFDRDKTIALIMNSSGSTGLPKGVALSGSFSTPVWISQAQGIRAGPGSSDKQEGEWPT
GLRLSRIGGIH

FLuc SecM(*Ec*) *N* = 225

MEDAKNIKKGPAPFYPLEDGTAGEQLHKAMKRYALVPGTIAFTDAHIEVDITYAEYFEMSVRLAEAMKRYGLNTNHRIVVC
SENSLQFFMPVLGALFIGVAVAPANDIYNERELLNSMGISQPTVVFVSKKGLQKILNVQKKLPIIQKIIIMDSKTDYQGFQS
MYTFVTSHLPPGFNEYDFVPESFDRDKTIALIMNSSGSTGLPSGSFSTPVWISQAQGIRAGPGSSDKQEGEWPTGLRLSR
IGGIH

FLuc SecM(*Ec*) *N* = 220

MEDAKNIKKGPAPFYPLEDGTAGEQLHKAMKRYALVPGTIAFTDAHIEVDITYAEYFEMSVRLAEAMKRYGLNTNHRIVVC
SENSLQFFMPVLGALFIGVAVAPANDIYNERELLNSMGISQPTVVFVSKKGLQKILNVQKKLPIIQKIIIMDSKTDYQGFQS
MYTFVTSHLPPGFNEYDFVPESFDRDKTIALIMNSSGSGSFSTPVWISQAQGIRAGPGSSDKQEGEWPTGLRLSRIGGIH

FLuc SecM(*Ec*) *N* = 215

MEDAKNIKKGPAPFYPLEDGTAGEQLHKAMKRYALVPGTIAFTDAHIEVDITYAEYFEMSVRLAEAMKRYGLNTNHRIVVC
SENSLQFFMPVLGALFIGVAVAPANDIYNERELLNSMGISQPTVVFVSKKGLQKILNVQKKLPIIQKIIIMDSKTDYQGFQS
MYTFVTSHLPPGFNEYDFVPESFDRDKTIALISGSFSTPVWISQAQGIRAGPGSSDKQEGEWPTGLRLSRIGGIH

FLuc SecM(*Ec*) *N* = 210

MEDAKNIKKGPAPFYPLEDGTAGEQLHKAMKRYALVPGTIAFTDAHIEVDITYAEYFEMSVRLAEAMKRYGLNTNHRIVVC
SENSLQFFMPVLGALFIGVAVAPANDIYNERELLNSMGISQPTVVFVSKKGLQKILNVQKKLPIIQKIIIMDSKTDYQGFQS
MYTFVTSHLPPGFNEYDFVPESFDRDKSGSFSTPVWISQAQGIRAGPGSSDKQEGEWPTGLRLSRIGGIH

FLuc SecM(*Ec*) *N* = 205

MEDAKNIKKGPAPFYPLEDGTAGEQLHKAMKRYALVPGTIAFTDAHIEVDITYAEYFEMSVRLAEAMKRYGLNTNHRIVVC
SENSLQFFMPVLGALFIGVAVAPANDIYNERELLNSMGISQPTVVFVSKKGLQKILNVQKKLPIIQKIIIMDSKTDYQGFQS
MYTFVTSHLPPGFNEYDFVPESSGSFSTPVWISQAQGIRAGPGSSDKQEGEWPTGLRLSRIGGIH

FLuc SecM(*Ec*) *N* = 200

MEDAKNIKKGPAPFYPLEDGTAGEQLHKAMKRYALVPGTIAFTDAHIEVDITYAEYFEMSVRLAEAMKRYGLNTNHRIVVC
SENSLQFFMPVLGALFIGVAVAPANDIYNERELLNSMGISQPTVVFVSKKGLQKILNVQKKLPIIQKIIIMDSKTDYQGFQS
MYTFVTSHLPPGFNEYDSGSFSTPVWISQAQGIRAGPGSSDKQEGEWPTGLRLSRIGGIH

FLuc SecM(*Ec*) *N* = 195

MEDAKNIKKGPAPFYPLEDGTAGEQLHKAMKRYALVPGTIAFTDAHIEVDITYAEYFEMSVRLAEAMKRYGLNTNHRIVVC
SENSLQFFMPVLGALFIGVAVAPANDIYNERELLNSMGISQPTVVFVSKKGLQKILNVQKKLPIIQKIIIMDSKTDYQGFQS
MYTFVTSHLPPGSGSFSTPVWISQAQGIRAGPGSSDKQEGEWPTGLRLSRIGGIH

FLuc SecM(*Ec*) *N* = 190

MEDAKNIKKGPAPFYPLEDGTAGEQLHKAMKRYALVPGTIAFTDAHIEVDITYAEYFEMSVRLAEAMKRYGLNTNHRIVVC
SENSLQFFMPVLGALFIGVAVAPANDIYNERELLNSMGISQPTVVFVSKKGLQKILNVQKKLPIIQKIIIMDSKTDYQGFQS
MYTFVTSSGSFSTPVWISQAQGIRAGPGSSDKQEGEWPTGLRLSRIGGIH

FLuc SecM(*Ec*) *N* = 185

MEDAKNIKKGPAPFYPLEDGTAGEQLHKAMKRYALVPGTIAFTDAHIEVDITYAEYFEMSVRLAEAMKRYGLNTNHRIVVC
SENSLQFFMPVLGALFIGVAVAPANDIYNERELLNSMGISQPTVVFVSKKGLQKILNVQKKLPIIQKIIIMDSKTDYQGFQS
MYSGSFSTPVWISQAQGIRAGPGSSDKQEGEWPTGLRLSRIGGIH

FLuc SecM(*Ec*) *N* = 180

MEDAKNIKKGPAPFYPLEDGTAGEQLHKAMKRYALVPGTIAFTDAHIEVDITYAEYFEMSVRLAEAMKRYGLNTNHRIVVC
SENSLQFFMPVLGALFIGVAVAPANDIYNERELLNSMGISQPTVVFVSKKGLQKILNVQKKLPIIQKIIIMDSKTDYQGSGSF
STPVWISQAQGIRAGPGSSDKQEGEWPTGLRLSRIGGIH

FLuc SecM(*Ec*) *N* = 175

MEDAKNIKKGPAPFYPLEDGTAGEQLHKAMKRYALVPGTIAFTDAHIEVDITYAEYFEMSVRLAEAMKRYGLNTNHRIVVC
SENSLQFFMPVLGALFIGVAVAPANDIYNERELLNSMGISQPTVVFVSKKGLQKILNVQKKLPIIQKIIIMDSKSGSFSTPVW
ISQAQGIRAGPGSSDKQEGEWPTGLRLSRIGGIH

FLuc SecM(*Ec*) *N* = 170

MEDAKNIKKGPAPFYPLEDGTAGEQLHKAMKRYALVPGTIAFTDAHIEVDITYAEYFEMSVRLAEAMKRYGLNTNHRIVVC
SENSLQFFMPVLGALFIGVAVAPANDIYNERELLNSMGISQPTVVFVSKKGLQKILNVQKKLPIIQKIISGSFSTPVWISQAQ
GIRAGPGSSDKQEGEWPTGLRLSRIGGIH

FLuc SecM(*Ec*) *N* = 167

MEDAKNIKKGPAPFYPLEDGTAGEQLHKAMKRYALVPGTIAFTDAHIEVDITYAEYFEMSVRLAEAMKRYGLNTNHRIVVC
SENSLQFFMPVLGALFIGVAVAPANDIYNERELLNSMGISQPTVVFVSKKGLQKILNVQKKLPIIQSGSFSTPVWISQAQGI
RAGPGSSDKQEGEWPTGLRLSRIGGIH

FLuc SecM(*Ec*) *N* = 165

MEDAKNIKKGPAPFYPLEDGTAGEQLHKAMKRYALVPGTIAFTDAHIEVDITYAEYFEMSVRLAEAMKRYGLNTNHRIVVC
SENSLQFFMPVLGALFIGVAVAPANDIYNERELLNSMGISQPTVVFVSKKGLQKILNVQKKLPISGSFSTPVWISQAQGIRA
GPGSSDKQEGEWPTGLRLSRIGGIH

FLuc SecM(*Ec*) *N* = 163

MEDAKNIKKGPAPFYPLEDGTAGEQLHKAMKRYALVPGTIAFTDAHIEVDITYAEYFEMSVRLAEAMKRYGLNTNHRIVVC
SENSLQFFMPVLGALFIGVAVAPANDIYNERELLNSMGISQPTVVFVSKKGLQKILNVQKKLSGSFSTPVWISQAQGIRAG
PGSSDKQEGEWPTGLRLSRIGGIH

FLuc SecM(*Ec*) *N* = 160

MEDAKNIKKGPAPFYPLEDGTAGEQLHKAMKRYALVPGTIAFTDAHIEVDITYAEYFEMSVRLAEAMKRYGLNTNHRIVVC
SENSLQFFMPVLGALFIGVAVAPANDIYNERELLNSMGISQPTVVFVSKKGLQKILNVQSGSFSTPVWISQAQGIRAGPG
SSDKQEGEWPTGLRLSRIGGIH

FLuc SecM(*Ec*) *N* = 155

MEDAKNIKKGPAPFYPLEDGTAGEQLHKAMKRYALVPGTIAFTDAHIEVDITYAEYFEMSVRLAEAMKRYGLNTNHRIVVC
SENSLQFFMPVLGALFIGVAVAPANDIYNERELLNSMGISQPTVVFVSKKGLQKSGSFSTPVWISQAQGIRAGPGSSDKQ
EGEWPTGLRLSRIGGIH

FLuc SecM(*Ec*) *N* = 150

MEDAKNIKKGPAPFYPLEDGTAGEQLHKAMKRYALVPGTIAFTDAHIEVDITYAEYFEMSVRLAEAMKRYGLNTNHRIVVC
SENSLQFFMPVLGALFIGVAVAPANDIYNERELLNSMGISQPTVVFVSKSGSFSTPVWISQAQGIRAGPGSSDKQEGEW
PTGLRLSRIGGIH

FLuc SecM(*Ec*) *N* = 145

MEDAKNIKKGPAPFYPLEDGTAGEQLHKAMKRYALVPGTIAFTDAHIEVDITYAEYFEMSVRLAEAMKRYGLNTNHRIVVC
SENSLQFFMPVLGALFIGVAVAPANDIYNERELLNSMGISQPTVSGSFSTPVWISQAQGIRAGPGSSDKQEGEWPTGLRL
SRIGGIH

FLuc SecM(*Ec*) *N* = 140

MEDAKNIKKGPAPFYPLEDGTAGEQLHKAMKRYALVPGTIAFTDAHIEVDITYAEYFEMSVRLAEAMKRYGLNTNHRIVVC
SENSLQFFMPVLGALFIGVAVAPANDIYNERELLNSMGISGSFSTPVWISQAQGIRAGPGSSDKQEGEWPTGLRLSRIGG
IH

FLuc SecM(*Ec*) *N* = 135

MEDAKNIKKGPAPFYPLEDGTAGEQLHKAMKRYALVPGTIAFTDAHIEVDITYAEYFEMSVRLAEAMKRYGLNTNHRIVVC
SENSLQFFMPVLGALFIGVAVAPANDIYNERELLSGSFSTPVWISQAQGIRAGPGSSDKQEGEWPTGLRLSRIGGIH

FLuc SecM(*Ec*) *N* = 130

MEDAKNIKKGPAPFYPLEDGTAGEQLHKAMKRYALVPGTIAFTDAHIEVDITYAEYFEMSVRLAEAMKRYGLNTNHRIVVCSENSLQFFMPVLGALFIGVAVAPANDIYNSGSFSTPVWISQAQGIRAGPGSSDKQEGEWPTGLRLSRIGGIH

FLuc SecM(*Ec*) *N* = 127

MEDAKNIKKGPAPFYPLEDGTAGEQLHKAMKRYALVPGTIAFTDAHIEVDITYAEYFEMSVRLAEAMKRYGLNTNHRIVVCSENSLQFFMPVLGALFIGVAVAPANDSGSFSTPVWISQAQGIRAGPGSSDKQEGEWPTGLRLSRIGGIH

FLuc SecM(*Ec*) *N* = 125

MEDAKNIKKGPAPFYPLEDGTAGEQLHKAMKRYALVPGTIAFTDAHIEVDITYAEYFEMSVRLAEAMKRYGLNTNHRIVVCSENSLQFFMPVLGALFIGVAVAPASGSFSTPVWISQAQGIRAGPGSSDKQEGEWPTGLRLSRIGGIH

FLuc SecM(*Ec*) *N* = 123

MEDAKNIKKGPAPFYPLEDGTAGEQLHKAMKRYALVPGTIAFTDAHIEVDITYAEYFEMSVRLAEAMKRYGLNTNHRIVVCSENSLQFFMPVLGALFIGVAVASGSFSTPVWISQAQGIRAGPGSSDKQEGEWPTGLRLSRIGGIH

FLuc SecM(*Ec*) *N* = 120

MEDAKNIKKGPAPFYPLEDGTAGEQLHKAMKRYALVPGTIAFTDAHIEVDITYAEYFEMSVRLAEAMKRYGLNTNHRIVVCSENSLQFFMPVLGALFIGVSGSFSTPVWISQAQGIRAGPGSSDKQEGEWPTGLRLSRIGGIH

FLuc SecM(*Ec*) *N* = 115

MEDAKNIKKGPAPFYPLEDGTAGEQLHKAMKRYALVPGTIAFTDAHIEVDITYAEYFEMSVRLAEAMKRYGLNTNHRIVVCSENSLQFFMPVLGASGSFSTPVWISQAQGIRAGPGSSDKQEGEWPTGLRLSRIGGIH

FLuc SecM(*Ec*) *N* = 110

MEDAKNIKKGPAPFYPLEDGTAGEQLHKAMKRYALVPGTIAFTDAHIEVDITYAEYFEMSVRLAEAMKRYGLNTNHRIVVCSENSLQFFMSGSFSTPVWISQAQGIRAGPGSSDKQEGEWPTGLRLSRIGGIH

FLuc SecM(*Ec*) *N* = 105

MEDAKNIKKGPAPFYPLEDGTAGEQLHKAMKRYALVPGTIAFTDAHIEVDITYAEYFEMSVRLAEAMKRYGLNTNHRIVVCSENSSGSFSTPVWISQAQGIRAGPGSSDKQEGEWPTGLRLSRIGGIH

FLuc SecM(*Ec*) *N* = 100

MEDAKNIKKGPAPFYPLEDGTAGEQLHKAMKRYALVPGTIAFTDAHIEVDITYAEYFEMSVRLAEAMKRYGLNTNHRIVVSGSFSTPVWISQAQGIRAGPGSSDKQEGEWPTGLRLSRIGGIH

FLuc SecM(*Ec*) *N* = 95

MEDAKNIKKGPAPFYPLEDGTAGEQLHKAMKRYALVPGTIAFTDAHIEVDITYAEYFEMSVRLAEAMKRYGLNTNSGSFSTPVWISQAQGIRAGPGSSDKQEGEWPTGLRLSRIGGIH

FLuc SecM(*Ec*) *N* = 90

MEDAKNIKKGPAPFYPLEDGTAGEQLHKAMKRYALVPGTIAFTDAHIEVDITYAEYFEMSVRLAEAMKRYSGSFSTPVWISQAQGIRAGPGSSDKQEGEWPTGLRLSRIGGIH

FLuc SecM(*Ec*) *N* = 85

MEDAKNIKKGPAPFYPLEDGTAGEQLHKAMKRYALVPGTIAFTDAHIEVDITYAEYFEMSVRLAESGSFSTPVWISQAQGIRAGPGSSDKQEGEWPTGLRLSRIGGIH

FLuc SecM(*Ec*) *N* = 80

MEDAKNIKKGPAPFYPLEDGTAGEQLHKAMKRYALVPGTIAFTDAHIEVDITYAEYFEMSSGSFSTPVWISQAQGIRAGPGSSDKQEGEWPTGLRLSRIGGIH

FLuc SecM(*Ec*) *N* = 75

MEDAKNIKKGPAPFYPLEDGTAGEQLHKAMKRYALVPGTIAFTDAHIEVDITYAESGSFSTPVWISQAQGIRAGPGSSDKQEGEWPTGLRLSRIGGIH

FLuc SecM(*Ec*) *N* = 70

MEDAKNIKKGPAPFYPLEDGTAGEQLHKAMKRYALVPGTIAFTDAHIEVDSGSFSTPVWISQAQGIRAGPGSSDKQEGEWPTGLRLSRIGGIH

**FLuc CTD-only constructs (FLuc residues 1-422 deleted)**

FLuc SecM(*Ec*) CTD *N* = 500

MPMPIAYWDEDEHFFIVDRLKSLIKYKGYQVAPAELESILLQHPNIFDAGVAGLPDDDAGELSGSFSTPVWISQAQGIRAGPDYIKRAVGLPGDKVTYDPVSKELTIQPGCSSGQACENALPVTYSNVEPSDFVGSSDKQEGEWPTGLRLSRIGGIH

FLuc SecM(*Ec*) CTD *N* = 505

MPMPIAYWDEDEHFFIVDRLKSLIKYKGYQVAPAELESILLQHPNIFDAGVAGLPDDDAGELPAAVVSGSFSTPVWISQAQGIRAGPDYIKRAVGLPGDKVTYDPVSKELTIQPGCSSGQACENALPVTYSNVEPSDFVGSSDKQEGEWPTGLRLSRIGGIH

FLuc SecM(*Ec*) CTD *N* = 510

MPMPIAYWDEDEHFFIVDRLKSLIKYKGYQVAPAELESILLQHPNIFDAGVAGLPDDDAGELPAAVVVLEHGSGSFSTPVWISQAQGIRAGPDYIKRAVGLPGDKVTYDPVSKELTIQPGCSSGQACENALPVTYSNVEPSDFVGSSDKQEGEWPTGLRLSRIGGIH

FLuc SecM(*Ec*) CTD *N* = 515

MPMPIAYWDEDEHFFIVDRLKSLIKYKGYQVAPAELESILLQHPNIFDAGVAGLPDDDAGELPAAVVVLEHGKTMTESGSFSTPVWISQAQGIRAGPDYIKRAVGLPGDKVTYDPVSKELTIQPGCSSGQACENALPVTYSNVEPSDFVGSSDKQEGEWPTGLRLSRIGGIH

FLuc SecM(*Ec*) CTD *N* = 520

MPMPIAYWDEDEHFFIVDRLKSLIKYKGYQVAPAELESILLQHPNIFDAGVAGLPDDDAGELPAAVVVLEHGKTMTEKEIVDSGSFSTPVWISQAQGIRAGPDYIKRAVGLPGDKVTYDPVSKELTIQPGCSSGQACENALPVTYSNVEPSDFVGSSDKQEGEWPTGLRLSRIGGIH

FLuc SecM(*Ec*) CTD *N* = 525

MPMPIAYWDEDEHFFIVDRLKSLIKYKGYQVAPAELESILLQHPNIFDAGVAGLPDDDAGELPAAVVVLEHGKTMTEKEIVDYVASQSGSFSTPVWISQAQGIRAGPDYIKRAVGLPGDKVTYDPVSKELTIQPGCSSGQACENALPVTYSNVEPSDFVGSSDKQEGEWPTGLRLSRIGGIH

FLuc SecM(*Ec*) CTD *N* = 530

MPMPIAYWDEDEHFFIVDRLKSLIKYKGYQVAPAELESILLQHPNIFDAGVAGLPDDDAGELPAAVVVLEHGKTMTEKEIVDYVASQVTTAKSGSFSTPVWISQAQGIRAGPDYIKRAVGLPGDKVTYDPVSKELTIQPGCSSGQACENALPVTYSNVEPSDFVGSSDKQEGEWPTGLRLSRIGGIH

FLuc SecM(*Ec*) CTD *N* = 535

MPMPIAYWDEDEHFFIVDRLKSLIKYKGYQVAPAELESILLQHPNIFDAGVAGLPDDDAGELPAAVVVLEHGKTMTEKEIV
DYVASQVTTAKKLRGGSGSFSTPVWISQAQGIRAGPDYIKRAVGLPGDKVTYDPVSKELTIQPGCSSGQACENALPVTYS
NVEPSDFVGSSDKQEGEWPTGLRLSRIGGIH

FLuc SecM(*Ec*) CTD *N* = 540

MPMPIAYWDEDEHFFIVDRLKSLIKYKGYQVAPAELESILLQHPNIFDAGVAGLPDDDAGELPAAVVVLEHGKTMTEKEIV
DYVASQVTTAKKLRGGVVFVDSGSFSTPVWISQAQGIRAGPDYIKRAVGLPGDKVTYDPVSKELTIQPGCSSGQACENAL
PVTYSNVEPSDFVGSSDKQEGEWPTGLRLSRIGGIH

FLuc SecM(*Ec*) CTD *N* = 545

MPMPIAYWDEDEHFFIVDRLKSLIKYKGYQVAPAELESILLQHPNIFDAGVAGLPDDDAGELPAAVVVLEHGKTMTEKEIV
DYVASQVTTAKKLRGGVVFVDEVPKGSGSFSTPVWISQAQGIRAGPDYIKRAVGLPGDKVTYDPVSKELTIQPGCSSGQA
CENALPVTYSNVEPSDFVGSSDKQEGEWPTGLRLSRIGGIH

FLuc SecM(*Ec*) CTD *N* = 550

MPMPIAYWDEDEHFFIVDRLKSLIKYKGYQVAPAELESILLQHPNIFDAGVAGLPDDDAGELPAAVVVLEHGKTMTEKEIV
DYVASQVTTAKKLRGGVVFVDEVPKGLTGKLSGSFSTPVWISQAQGIRAGPDYIKRAVGLPGDKVTYDPVSKELTIQPGCS
SGQACENALPVTYSNVEPSDFVGSSDKQEGEWPTGLRLSRIGGIH

FLuc SecM(*Ec*) CTD *N* = 555

MPMPIAYWDEDEHFFIVDRLKSLIKYKGYQVAPAELESILLQHPNIFDAGVAGLPDDDAGELPAAVVVLEHGKTMTEKEIV
DYVASQVTTAKKLRGGVVFVDEVPKGLTGKLDARKISGSFSTPVWISQAQGIRAGPDYIKRAVGLPGDKVTYDPVSKELTI
QPGCSSGQACENALPVTYSNVEPSDFVGSSDKQEGEWPTGLRLSRIGGIH

FLuc SecM(*Ec*) CTD *N* = 560

MPMPIAYWDEDEHFFIVDRLKSLIKYKGYQVAPAELESILLQHPNIFDAGVAGLPDDDAGELPAAVVVLEHGKTMTEKEIV
DYVASQVTTAKKLRGGVVFVDEVPKGLTGKLDARKIREILISGSFSTPVWISQAQGIRAGPDYIKRAVGLPGDKVTYDPVSK
ELTIQPGCSSGQACENALPVTYSNVEPSDFVGSSDKQEGEWPTGLRLSRIGGIH

FLuc SecM(*Ec*) CTD *N* = 565

MPMPIAYWDEDEHFFIVDRLKSLIKYKGYQVAPAELESILLQHPNIFDAGVAGLPDDDAGELPAAVVVLEHGKTMTEKEIV
DYVASQVTTAKKLRGGVVFVDEVPKGLTGKLDARKIREILIKAKKGSGSFSTPVWISQAQGIRAGPDYIKRAVGLPGDKVTY
DPVSKELTIQPGCSSGQACENALPVTYSNVEPSDFVGSSDKQEGEWPTGLRLSRIGGIH

FLuc SecM(*Ec*) CTD *N* = 570

MPMPIAYWDEDEHFFIVDRLKSLIKYKGYQVAPAELESILLQHPNIFDAGVAGLPDDDAGELPAAVVVLEHGKTMTEKEIV
DYVASQVTTAKKLRGGVVFVDEVPKGLTGKLDARKIREILIKAKKGGKIAVSGSFSTPVWISQAQGIRAGPDYIKRAVGLPG
DKVTYDPVSKELTIQPGCSSGQACENALPVTYSNVEPSDFVGSSDKQEGEWPTGLRLSRIGGIH

FLuc SecM(*Ec*) CTD *N* = 575

MPMPIAYWDEDEHFFIVDRLKSLIKYKGYQVAPAELESILLQHPNIFDAGVAGLPDDDAGELPAAVVVLEHGKTMTEKEIV
DYVASQVTTAKKLRGGVVFVDEVPKGLTGKLDARKIREILIKAKKGGKIAVSGSGSGSFSTPVWISQAQGIRAGPDYIKRA
VGLPGDKVTYDPVSKELTIQPGCSSGQACENALPVTYSNVEPSDFVGSSDKQEGEWPTGLRLSRIGGIH

FLuc SecM(*Ec*) CTD *N* = 580

MPMPIAYWDEDEHFFIVDRLKSLIKYKGYQVAPAELESILLQHPNIFDAGVAGLPDDDAGELPAAVVVLEHGKTMTEKEIV
DYVASQVTTAKKLRGGVVFVDEVPKGLTGKLDARKIREILIKAKKGGKIAVSGSGSGSGSGSGSFSTPVWISQAQGIRAGP
DYIKRAVGLPGDKVTYDPVSKELTIQPGCSSGQACENALPVTYSNVEPSDFVGSSDKQEGEWPTGLRLSRIGGIH

FLuc SecM(*Ec*) CTD *N* = 585

MPMPIAYWDEDEHFFIVDRLKSLIKYKGYQVAPAELESILLQHPNIFDAGVAGLPDDDAGELPAAVVVLEHGKTMTEKEIVDYVASQVTTAKKLRGGVVFVDEVPKGLTGKLDARKIREILIKAKKGGKIAVSGSGSGSGSGSGSGSGSGFSTPVWISQAQGIRAGPDYIKRAVGLPGDKVTYDPVSKELTIQPGCSSGQACENALPVTYSNVEPSDFVGSSDKQEGEWPTGLRLSRIGGIH

FLuc SecM(*Ec*) CTD *N* = 592

MPMPIAYWDEDEHFFIVDRLKSLIKYKGYQVAPAELESILLQHPNIFDAGVAGLPDDDAGELPAAVVVLEHGKTMTEKEIVDYVASQVTTAKKLRGGVVFVDEVPKGLTGKLDARKIREILIKAKKGGKIAVSGSGSGSGSGSGSGSGSGSGSGSGGFSTPVWISQAQGIRAGPDYIKRAVGLPGDKVTYDPVSKELTIQPGCSSGQACENALPVTYSNVEPSDFVGSSDKQEGEWPTGLRLSRIGGIH

FLuc SecM(*Ec*) CTD *N* = 600

MPMPIAYWDEDEHFFIVDRLKSLIKYKGYQVAPAELESILLQHPNIFDAGVAGLPDDDAGELPAAVVVLEHGKTMTEKEIVDYVASQVTTAKKLRGGVVFVDEVPKGLTGKLDARKIREILIKAKKGGKIAVSGSGSGSGSGSGSGSGSGSGSGSGGSGSGSGSGFSTPVWISQAQGIRAGPDYIKRAVGLPGDKVTYDPVSKELTIQPGCSSGQACENALPVTYSNVEPSDFVGSSDKQEGEWPTGLRLSRIGGIH

FLuc SecM(*Ec*) CTD *N* = 605

MPMPIAYWDEDEHFFIVDRLKSLIKYKGYQVAPAELESILLQHPNIFDAGVAGLPDDDAGELPAAVVVLEHGKTMTEKEIVDYVASQVTTAKKLRGGVVFVDEVPKGLTGKLDARKIREILIKAKKGGKIAVSGSGSGSGSGSGSGSGSGSGSGSGGSGSGSGSGSGSGSFSTPVWISQAQGIRAGPDYIKRAVGLPGDKVTYDPVSKELTIQPGCSSGQACENALPVTYSNVEPSDFVGSSDKQEGEWPTGLRLSRIGGIH

FLuc SecM(*Ec*) CTD *N* = 619

MPMPIAYWDEDEHFFIVDRLKSLIKYKGYQVAPAELESILLQHPNIFDAGVAGLPDDDAGELPAAVVVLEHGKTMTEKEIVDYVASQVTTAKKLRGGVVFVDEVPKGLTGKLDARKIREILIKAKKGGKIAVSGSGSGSGSGSGSGSGSGSGSGSGGSGSGSGSGSGSGSGSGSGSGSGSGSGSGSFSTPVWISQAQGIRAGPDYIKRAVGLPGDKVTYDPVSKELTIQPGCSSGQACENALPVTYSNVEPSDFVGSSDKQEGEWPTGLRLSRIGGIH

**FLuc N-terminal deletions**

FLuc SecM(*Ec*) Δ4-213 *N* = 300

MEDTACVRFSHARDPIFGNQIIPDTAILSVVPFHHGFGMFTTLGYLICGFRVVLMYRFEEELFLRSLQDYSGSFSTPVWISQAQGIRAGPDYIKRAVGLPGDKVTYDPVSKELTIQPGCSSGQACENALPVTYSNVEPSDFVGSSDKQEGEWPTGLRLSRIGGIH

FLuc SecM(*Ec*) Δ4-173 *N* = 300

MEDPGFNEYDFVPESFDRDKTIALIMNSSGSTGLPKGVALPHRTACVRFSHARDPIFGNQIIPDTAILSVVPFHHGFGMFTTLGYLICGFRVVLMYRFEEELFLRSLQDYSGSFSTPVWISQAQGIRAGPDYIKRAVGLPGDKVTYDPVSKELTIQPGCSSGQACENALPVTYSNVEPSDFVGSSDKQEGEWPTGLRLSRIGGIH

FLuc SecM(*Ec*) Δ4-153 *N* = 300

MEDSKTDYQGFQSMYTFVTSHLPPGFNEYDFVPESFDRDKTIALIMNSSGSTGLPKGVALPHRTACVRFSHARDPIFGNQIIPDTAILSVVPFHHGFGMFTTLGYLICGFRVVLMYRFEEELFLRSLQDYSGSFSTPVWISQAQGIRAGPDYIKRAVGLPGDKVTYDPVSKELTIQPGCSSGQACENALPVTYSNVEPSDFVGSSDKQEGEWPTGLRLSRIGGIH

FLuc SecM(*Ec*) Δ4-138 *N* = 300

MEDVQKKLPIIQKIIIMDSKTDYQGFQSMYTFVTSHLPPGFNEYDFVPESFDRDKTIALIMNSSGSTGLPKGVALPHRTACVRFSHARDPIFGNQIIPDTAILSVVPFHHGFGMFTTLGYLICGFRVVLMYRFEEELFLRSLQDYSGSFSTPVWISQAQGIRAGPDYIKRAVGLPGDKVTYDPVSKELTIQPGCSSGQACENALPVTYSNVEPSDFVGSSDKQEGEWPTGLRLSRIGGIH

FLuc SecM(*Ec*) Δ4-123 *N* = 300

MEDTVVFVSKKGLQKILNVQKKLPIIQKIIIMDSKTDYQGFQSMYTFVTSHLPPGFNEYDFVPESFDRDKTIALIMNSSGSTGLPKGVALPHRTACVRFSHARDPIFGNQIIPDTAILSVVPFHHGFGMFTTLGYLICGFRVVLMYRFEEELFLRSLQDYSGSFSTPVWISQAQGIRAGPDYIKRAVGLPGDKVTYDPVSKELTIQPGCSSGQACENALPVTYSNVEPSDFVGSSDKQEGEWPTGLRLSRIGGIH

FLuc SecM(*Ec*) Δ4-113 *N* = 300

MEDLLNSMGISQPTVVFVSKKGLQKILNVQKKLPIIQKIIIMDSKTDYQGFQSMYTFVTSHLPPGFNEYDFVPESFDRDKTIALIMNSSGSTGLPKGVALPHRTACVRFSHARDPIFGNQIIPDTAILSVVPFHHGFGMFTTLGYLICGFRVVLMYRFEEELFLRSLQDYSGSFSTPVWISQAQGIRAGPDYIKRAVGLPGDKVTYDPVSKELTIQPGCSSGQACENALPVTYSNVEPSDFVGSSDKQEGEWPTGLRLSRIGGIH

FLuc SecM(*Ec*) Δ4-103 *N* = 300

MEDPANDIYNERELLNSMGISQPTVVFVSKKGLQKILNVQKKLPIIQKIIIMDSKTDYQGFQSMYTFVTSHLPPGFNEYDFVPESFDRDKTIALIMNSSGSTGLPKGVALPHRTACVRFSHARDPIFGNQIIPDTAILSVVPFHHGFGMFTTLGYLICGFRVVLMYRFEEELFLRSLQDYSGSFSTPVWISQAQGIRAGPDYIKRAVGLPGDKVTYDPVSKELTIQPGCSSGQACENALPVTYSNVEPSDFVGSSDKQEGEWPTGLRLSRIGGIH

FLuc SecM(*Ec*) Δ4-93 *N* = 300

MEDGALFIGVAVAPANDIYNERELLNSMGISQPTVVFVSKKGLQKILNVQKKLPIIQKIIIMDSKTDYQGFQSMYTFVTSHLPPGFNEYDFVPESFDRDKTIALIMNSSGSTGLPKGVALPHRTACVRFSHARDPIFGNQIIPDTAILSVVPFHHGFGMFTTLGYLICGFRVVLMYRFEEELFLRSLQDYSGSFSTPVWISQAQGIRAGPDYIKRAVGLPGDKVTYDPVSKELTIQPGCSSGQACENALPVTYSNVEPSDFVGSSDKQEGEWPTGLRLSRIGGIH

FLuc SecM(*Ec*) Δ4-83 *N* = 300

MEDNSLQFFMPVLGALFIGVAVAPANDIYNERELLNSMGISQPTVVFVSKKGLQKILNVQKKLPIIQKIIIMDSKTDYQGFQSMYTFVTSHLPPGFNEYDFVPESFDRDKTIALIMNSSGSTGLPKGVALPHRTACVRFSHARDPIFGNQIIPDTAILSVVPFHHGFGMFTTLGYLICGFRVVLMYRFEEELFLRSLQDYSGSFSTPVWISQAQGIRAGPDYIKRAVGLPGDKVTYDPVSKELTIQPGCSSGQACENALPVTYSNVEPSDFVGSSDKQEGEWPTGLRLSRIGGIH

FLuc SecM(*Ec*) Δ4-73 *N* = 300

MEDTNHRIVVCSENSLQFFMPVLGALFIGVAVAPANDIYNERELLNSMGISQPTVVFVSKKGLQKILNVQKKLPIIQKIIIMDSKTDYQGFQSMYTFVTSHLPPGFNEYDFVPESFDRDKTIALIMNSSGSTGLPKGVALPHRTACVRFSHARDPIFGNQIIPDTAILSVVPFHHGFGMFTTLGYLICGFRVVLMYRFEEELFLRSLQDYSGSFSTPVWISQAQGIRAGPDYIKRAVGLPGDKVTYDPVSKELTIQPGCSSGQACENALPVTYSNVEPSDFVGSSDKQEGEWPTGLRLSRIGGIH

FLuc SecM(*Ec*) Δ4-63 *N* = 300

MEDAEAMKRYGLNTNHRIVVCSENSLQFFMPVLGALFIGVAVAPANDIYNERELLNSMGISQPTVVFVSKKGLQKILNVQKKLPIIQKIIIMDSKTDYQGFQSMYTFVTSHLPPGFNEYDFVPESFDRDKTIALIMNSSGSTGLPKGVALPHRTACVRFSHARDPIFGNQIIPDTAILSVVPFHHGFGMFTTLGYLICGFRVVLMYRFEEELFLRSLQDYSGSFSTPVWISQAQGIRAGPDYIKRAVGLPGDKVTYDPVSKELTIQPGCSSGQACENALPVTYSNVEPSDFVGSSDKQEGEWPTGLRLSRIGGIH

FLuc SecM(*Ec*) Δ4-53 *N* = 300

MEDAEYFEMSVRLAEAMKRYGLNTNHRIVVCSENSLQFFMPVLGALFIGVAVAPANDIYNERELLNSMGISQPTVVFVSKKGLQKILNVQKKLPIIQKIIIMDSKTDYQGFQSMYTFVTSHLPPGFNEYDFVPESFDRDKTIALIMNSSGSTGLPKGVALPHRTACVRFSHARDPIFGNQIIPDTAILSVVPFHHGFGMFTTLGYLICGFRVVLMYRFEEELFLRSLQDYSGSFSTPVWISQAQ

GIRAGPDYIKRAVGLPGDKVTYDPVSKELTIQPGCSSGQACENALPVTYSNVEPSDFVGSSDKQEGEWPTGLRLSRIGGI H

FLuc SecM(*Ec*) Δ4-43 *N* = 300

MEDDAHIEVDITYAEYFEMSVRLAEAMKRYGLNTNHRIVVCSENSLQFFMPVLGALFIGVAVAPANDIYNERELLNSMGIS QPTVVFVSKKGLQKILNVQKKLPIIQKIIIMDSKTDYQGFQSMYTFVTSHLPPGFNEYDFVPESFDRDKTIALIMNSSGSTGL PKGVALPHRTACVRFSHARDPIFGNQIIPDTAILSVVPFHHGFGMFTTLGYLICGFRVVLMYRFEEELFLRSLQDYSGSFST PVWISQAQGIRAGPDYIKRAVGLPGDKVTYDPVSKELTIQPGCSSGQACENALPVTYSNVEPSDFVGSSDKQEGEWPTG LRLSRIGGIH

FLuc SecM(*Ec*) Δ4-33 *N* = 300

MEDALVPGTIAFTDAHIEVDITYAEYFEMSVRLAEAMKRYGLNTNHRIVVCSENSLQFFMPVLGALFIGVAVAPANDIYNER ELLNSMGISQPTVVFVSKKGLQKILNVQKKLPIIQKIIIMDSKTDYQGFQSMYTFVTSHLPPGFNEYDFVPESFDRDKTIALI MNSSGSTGLPKGVALPHRTACVRFSHARDPIFGNQIIPDTAILSVVPFHHGFGMFTTLGYLICGFRVVLMYRFEEELFLRS LQDYSGSFSTPVWISQAQGIRAGPDYIKRAVGLPGDKVTYDPVSKELTIQPGCSSGQACENALPVTYSNVEPSDFVGSSD KQEGEWPTGLRLSRIGGIH

FLuc SecM(*Ec*) Δ4-23 *N* = 300

MEDEQLHKAMKRYALVPGTIAFTDAHIEVDITYAEYFEMSVRLAEAMKRYGLNTNHRIVVCSENSLQFFMPVLGALFIGVA VAPANDIYNERELLNSMGISQPTVVFVSKKGLQKILNVQKKLPIIQKIIIMDSKTDYQGFQSMYTFVTSHLPPGFNEYDFVP ESFDRDKTIALIMNSSGSTGLPKGVALPHRTACVRFSHARDPIFGNQIIPDTAILSVVPFHHGFGMFTTLGYLICGFRVVLM YRFEEELFLRSLQDYSGSFSTPVWISQAQGIRAGPDYIKRAVGLPGDKVTYDPVSKELTIQPGCSSGQACENALPVTYSNV EPSDFVGSSDKQEGEWPTGLRLSRIGGIH

FLuc SecM(*Ec*) Δ4-13 *N* = 300

MEDFYPLEDGTAGEQLHKAMKRYALVPGTIAFTDAHIEVDITYAEYFEMSVRLAEAMKRYGLNTNHRIVVCSENSLQFFM PVLGALFIGVAVAPANDIYNERELLNSMGISQPTVVFVSKKGLQKILNVQKKLPIIQKIIIMDSKTDYQGFQSMYTFVTSHLP PGFNEYDFVPESFDRDKTIALIMNSSGSTGLPKGVALPHRTACVRFSHARDPIFGNQIIPDTAILSVVPFHHGFGMFTTLG YLICGFRVVLMYRFEEELFLRSLQDYSGSFSTPVWISQAQGIRAGPDYIKRAVGLPGDKVTYDPVSKELTIQPGCSSGQAC ENALPVTYSNVEPSDFVGSSDKQEGEWPTGLRLSRIGGIH

**FLuc GS-scanning constructs**

FLuc SecM(*Ec*) *N* = 300 15GS

MEDAKNIKKGPAPFYPLEDGTAGEQLHKAMKRYALVPGTIAFTDAHIEVDITYAEYFEMSVRLAEAMKRYGLNTNHRIVVC SENSLQFFMPVLGALFIGVAVAPANDIYNERELLNSMGISQPTVVFVSKKGLQKILNVQKKLPIIQKIIIMDSKTDYQGFQS MYTFVTSHLPPGFNEYDFVPESFDRDKTIALIMNSSGSTGLPKGVALPHRTACVRFSHARDPIFGNQIIPDTAILSVVPFHH GFGMFTTLGYLICGFRVVLMSGSGGSGSGSGSGSGSGSFSTPVWISQAQGIRAGPDYIKRAVGLPGDKVTYDPVSKELTI QPGCSSGQACENALPVTYSNVEPSDFVGSSDKQEGEWPTGLRLSRIGGIH

FLuc SecM(*Ec*) *N* = 300 20GS

MEDAKNIKKGPAPFYPLEDGTAGEQLHKAMKRYALVPGTIAFTDAHIEVDITYAEYFEMSVRLAEAMKRYGLNTNHRIVVC SENSLQFFMPVLGALFIGVAVAPANDIYNERELLNSMGISQPTVVFVSKKGLQKILNVQKKLPIIQKIIIMDSKTDYQGFQS MYTFVTSHLPPGFNEYDFVPESFDRDKTIALIMNSSGSTGLPKGVALPHRTACVRFSHARDPIFGNQIIPDTAILSVVPFHH GFGMFTTLGYLICGFGSGSGSGSGGSGSGSGSGSGSGSFSTPVWISQAQGIRAGPDYIKRAVGLPGDKVTYDPVSKELT IQPGCSSGQACENALPVTYSNVEPSDFVGSSDKQEGEWPTGLRLSRIGGIH

FLuc SecM(*Ec*) *N* = 300 25GS

MEDAKNIKKGPAPFYPLEDGTAGEQLHKAMKRYALVPGTIAFTDAHIEVDITYAEYFEMSVRLAEAMKRYGLNTNHRIVVC
SENSLQFFMPVLGALFIGVAVAPANDIYNERELLNSMGISQPTVVFVSKKGLQKILNVQKKLPIIQKIIIMDSKTDYQGFQS
MYTFVTSHLPPGFNEYDFVPESFDRDKTIALIMNSSGSTGLPKGVALPHRTACVRFSHARDPIFGNQIIPDTAILSVVPFHH
GFGMFTTLGYSGSGSGSGSGSGSGGSGSGSGSGSGSGSFSTPVWISQAQGIRAGPDYIKRAVGLPGDKVTYDPVSKEL
TIQPGCSSGQACENALPVTYSNVEPSDFVGSSDKQEGEWPTGLRLSRIGGIH

FLuc SecM(*Ec*) *N* = 300 30GS

MEDAKNIKKGPAPFYPLEDGTAGEQLHKAMKRYALVPGTIAFTDAHIEVDITYAEYFEMSVRLAEAMKRYGLNTNHRIVVC
SENSLQFFMPVLGALFIGVAVAPANDIYNERELLNSMGISQPTVVFVSKKGLQKILNVQKKLPIIQKIIIMDSKTDYQGFQS
MYTFVTSHLPPGFNEYDFVPESFDRDKTIALIMNSSGSTGLPKGVALPHRTACVRFSHARDPIFGNQIIPDTAILSVVPFHH
GFGMFGSGSGSGSGSGSGSGSGSGGSGSGSGSGSGSGSFSTPVWISQAQGIRAGPDYIKRAVGLPGDKVTYDPVSK
ELTIQPGCSSGQACENALPVTYSNVEPSDFVGSSDKQEGEWPTGLRLSRIGGIH

FLuc SecM(*Ec*) *N* = 400 3GS

MEDAKNIKKGPAPFYPLEDGTAGEQLHKAMKRYALVPGTIAFTDAHIEVDITYAEYFEMSVRLAEAMKRYGLNTNHRIVVC
SENSLQFFMPVLGALFIGVAVAPANDIYNERELLNSMGISQPTVVFVSKKGLQKILNVQKKLPIIQKIIIMDSKTDYQGFQS
MYTFVTSHLPPGFNEYDFVPESFDRDKTIALIMNSSGSTGLPKGVALPHRTACVRFSHARDPIFGNQIIPDTAILSVVPFHH
GFGMFTTLGYLICGFRVVLMYRFEEELFLRSLQDYKIQSALLVPTLFSFFAKSTLIDKYDLSNLHEIASGGAPLSKEVGEAVA
KRFHLPGIRQGYGLTETTSAILITPEGDDKPGAVGKVVPFFEAKVVDLDGSGSGSFSTPVWISQAQGIRAGPDYIKRAVGL
PGDKVTYDPVSKELTIQPGCSSGQACENALPVTYSNVEPSDFVGSSDKQEGEWPTGLRLSRIGGIH

FLuc SecM(*Ec*) *N* = 400 6GS

MEDAKNIKKGPAPFYPLEDGTAGEQLHKAMKRYALVPGTIAFTDAHIEVDITYAEYFEMSVRLAEAMKRYGLNTNHRIVVC
SENSLQFFMPVLGALFIGVAVAPANDIYNERELLNSMGISQPTVVFVSKKGLQKILNVQKKLPIIQKIIIMDSKTDYQGFQS
MYTFVTSHLPPGFNEYDFVPESFDRDKTIALIMNSSGSTGLPKGVALPHRTACVRFSHARDPIFGNQIIPDTAILSVVPFHH
GFGMFTTLGYLICGFRVVLMYRFEEELFLRSLQDYKIQSALLVPTLFSFFAKSTLIDKYDLSNLHEIASGGAPLSKEVGEAVA
KRFHLPGIRQGYGLTETTSAILITPEGDDKPGAVGKVVPFFEAKVVSGSGSGSGSFSTPVWISQAQGIRAGPDYIKRAVGL
PGDKVTYDPVSKELTIQPGCSSGQACENALPVTYSNVEPSDFVGSSDKQEGEWPTGLRLSRIGGIH

FLuc SecM(*Ec*) *N* = 400 9GS

MEDAKNIKKGPAPFYPLEDGTAGEQLHKAMKRYALVPGTIAFTDAHIEVDITYAEYFEMSVRLAEAMKRYGLNTNHRIVVC
SENSLQFFMPVLGALFIGVAVAPANDIYNERELLNSMGISQPTVVFVSKKGLQKILNVQKKLPIIQKIIIMDSKTDYQGFQS
MYTFVTSHLPPGFNEYDFVPESFDRDKTIALIMNSSGSTGLPKGVALPHRTACVRFSHARDPIFGNQIIPDTAILSVVPFHH
GFGMFTTLGYLICGFRVVLMYRFEEELFLRSLQDYKIQSALLVPTLFSFFAKSTLIDKYDLSNLHEIASGGAPLSKEVGEAVA
KRFHLPGIRQGYGLTETTSAILITPEGDDKPGAVGKVVPFFEAGSGSGSGSGSGSFSTPVWISQAQGIRAGPDYIKRAVGL
PGDKVTYDPVSKELTIQPGCSSGQACENALPVTYSNVEPSDFVGSSDKQEGEWPTGLRLSRIGGIH

FLuc SecM(*Ec*) *N* = 400 12GS

MEDAKNIKKGPAPFYPLEDGTAGEQLHKAMKRYALVPGTIAFTDAHIEVDITYAEYFEMSVRLAEAMKRYGLNTNHRIVVC
SENSLQFFMPVLGALFIGVAVAPANDIYNERELLNSMGISQPTVVFVSKKGLQKILNVQKKLPIIQKIIIMDSKTDYQGFQS
MYTFVTSHLPPGFNEYDFVPESFDRDKTIALIMNSSGSTGLPKGVALPHRTACVRFSHARDPIFGNQIIPDTAILSVVPFHH
GFGMFTTLGYLICGFRVVLMYRFEEELFLRSLQDYKIQSALLVPTLFSFFAKSTLIDKYDLSNLHEIASGGAPLSKEVGEAVA
KRFHLPGIRQGYGLTETTSAILITPEGDDKPGAVGKVVPFGGSGSGSGSGSGSGSFSTPVWISQAQGIRAGPDYIKRAVG
LPGDKVTYDPVSKELTIQPGCSSGQACENALPVTYSNVEPSDFVGSSDKQEGEWPTGLRLSRIGGIH

FLuc SecM(*Ec*) *N* = 400 15GS

MEDAKNIKKGPAPFYPLEDGTAGEQLHKAMKRYALVPGTIAFTDAHIEVDITYAEYFEMSVRLAEAMKRYGLNTNHRIVVC
SENSLQFFMPVLGALFIGVAVAPANDIYNERELLNSMGISQPTVVFVSKKGLQKILNVQKKLPIIQKIIIMDSKTDYQGFQS
MYTFVTSHLPPGFNEYDFVPESFDRDKTIALIMNSSGSTGLPKGVALPHRTACVRFSHARDPIFGNQIIPDTAILSVVPFHH
GFGMFTTLGYLICGFRVVLMYRFEEELFLRSLQDYKIQSALLVPTLFSFFAKSTLIDKYDLSNLHEIASGGAPLSKEVGEAVA

KRFHLPGIRQGYGLTETTSAILITPEGDDKPGAVGKVSGSGGSGSGSGSGSGSFSTPVWISQAQGIRAGPDYIKRAVG
LPGDKVTYDPVSKELTIQPGCSSGQACENALPVTYSNVEPSDFVGSSDKQEGEWPTGLRLSRIGGIH

FLuc SecM(*Ec*) *N* = 400 20GS

MEDAKNIKKGPAPFYPLEDGTAGEQLHKAMKRYALVPGTIAFTDAHIEVDITYAEYFEMSVRLAEAMKRYGLNTNHRIVVC
SENSLQFFMPVLGALFIGVAVAPANDIYNERELLNSMGISQPTVVFVSKKGLQKILNVQKKLPIIQKIIIMDSKTDYQGFQS
MYTFVTSHLPPGFNEYDFVPESFDRDKTIALIMNSSGSTGLPKGVALPHRTACVRFSHARDPIFGNQIIPDTAILSVVPFHH
GFGMFTTLGYLICGFRVVLMYRFEEELFLRSLQDYKIQSALLVPTLFSFFAKSTLIDKYDLSNLHEIASGGAPLSKEVGEAVA
KRFHLPGIRQGYGLTETTSAILITPEGDDKPGGSGSGSGSGGSGSGSGSGSGSFSTPVWISQAQGIRAGPDYIKRAV
GLPGDKVTYDPVSKELTIQPGCSSGQACENALPVTYSNVEPSDFVGSSDKQEGEWPTGLRLSRIGGIH

FLuc SecM(*Ec*) *N* = 400 25GS

MEDAKNIKKGPAPFYPLEDGTAGEQLHKAMKRYALVPGTIAFTDAHIEVDITYAEYFEMSVRLAEAMKRYGLNTNHRIVVC
SENSLQFFMPVLGALFIGVAVAPANDIYNERELLNSMGISQPTVVFVSKKGLQKILNVQKKLPIIQKIIIMDSKTDYQGFQS
MYTFVTSHLPPGFNEYDFVPESFDRDKTIALIMNSSGSTGLPKGVALPHRTACVRFSHARDPIFGNQIIPDTAILSVVPFHH
GFGMFTTLGYLICGFRVVLMYRFEEELFLRSLQDYKIQSALLVPTLFSFFAKSTLIDKYDLSNLHEIASGGAPLSKEVGEAVA
KRFHLPGIRQGYGLTETTSAILITPEGSGSGSGSGSGSGGSGSGSGSGSGSGSFSTPVWISQAQGIRAGPDYIKRAVG
LPGDKVTYDPVSKELTIQPGCSSGQACENALPVTYSNVEPSDFVGSSDKQEGEWPTGLRLSRIGGIH

FLuc SecM(*Ec*) *N* = 400 30GS

MEDAKNIKKGPAPFYPLEDGTAGEQLHKAMKRYALVPGTIAFTDAHIEVDITYAEYFEMSVRLAEAMKRYGLNTNHRIVVC
SENSLQFFMPVLGALFIGVAVAPANDIYNERELLNSMGISQPTVVFVSKKGLQKILNVQKKLPIIQKIIIMDSKTDYQGFQS
MYTFVTSHLPPGFNEYDFVPESFDRDKTIALIMNSSGSTGLPKGVALPHRTACVRFSHARDPIFGNQIIPDTAILSVVPFHH
GFGMFTTLGYLICGFRVVLMYRFEEELFLRSLQDYKIQSALLVPTLFSFFAKSTLIDKYDLSNLHEIASGGAPLSKEVGEAVA
KRFHLPGIRQGYGLTETTSAILGSGSGSGSGSGSGSGSGGSGSGSGSGSGSGSFSTPVWISQAQGIRAGPDYIKRAV
GLPGDKVTYDPVSKELTIQPGCSSGQACENALPVTYSNVEPSDFVGSSDKQEGEWPTGLRLSRIGGIH

***Cryo-EM constructs***

**Color code:** FLuc 6x His Linker SecM (3W) AP C-tail

FLuc SecM(3W) *N* = 110

MHHHHHHEDAKNIKKGPAPFYPLEDGTAGEQLHKAMKRYALVPGTIAFTDAHIEVDITYAEYFEMSVRLAEAMKRYGLNT
NHRIVVCSENSLQFFMSGSFSTPVWIWWWPPIRGSPGSSDKQEGEWPTGLRLSRIGGIH

FLuc SecM(3W) *N* = 130

MHHHHHHEDAKNIKKGPAPFYPLEDGTAGEQLHKAMKRYALVPGTIAFTDAHIEVDITYAEYFEMSVRLAEAMKRYGLNT
NHRIVVCSENSLQFFMPVLGALFIGVAVAPANDIYNSGSFSTPVWIWWWPPIRGSPGSSDKQEGEWPTGLRLSRIGGIH

FLuc SecM(3W) *N* = 190

MHHHHHHEDAKNIKKGPAPFYPLEDGTAGEQLHKAMKRYALVPGTIAFTDAHIEVDITYAEYFEMSVRLAEAMKRYGLNT
NHRIVVCSENSLQFFMPVLGALFIGVAVAPANDIYNERELLNSMGISQPTVVFVSKKGLQKILNVQKKLPIIQKIIIMDSKTD
YQGFQSMYTFVTSSGSFSTPVWIWWWPPIRGSPGSSDKQEGEWPTGLRLSRIGGIH
